# Community structure of heritable viruses in a *Drosophila*-parasitoids complex

**DOI:** 10.1101/2023.07.29.551099

**Authors:** Julien Varaldi, David Lepetit, Nelly Burlet, Camille Faber, Bérénice Baretje, Rolland Allemand

## Abstract

The diversity and phenotypic impacts related to the presence of heritable bacteria in insects have been extensively studied in the last decades. On the contrary, heritable viruses have been overlooked for several reasons, including technical ones. This is regrettable because of the size of this gap knowledge and because case study indicate that viruses may have profound impact on the functionning of individuals and communities. Additionally, the factors that may shape viral communities are poorly known, except in some very specific viral-insect systems. Here we analyze the community structure of heritable viruses in a multi-hosts-multi-parasitoids community. Drosophilidae and their larval and pupal parasitoids were sampled in two locations, two years and two seasons. After two lab generations, putative DNA and RNA viruses were purified and sequenced. Our analysis revealed the presence of 53 viruses (including 37 new viruses), the great majority of which were RNA viruses. The "species" factor was by far the most significant one, explaining more than 50% of the variance in viral structure. Additionally, parasitoids had a higher number of heritable viruses compared to their hosts, suggesting that this lifestyle favours the association with viruses. Finally, our community-level survey challenged previous interpretation concerning the host range of some previously described viruses.

## 1 Introduction

Insects have a close relationship with a wide range of heritable bacteria that can shape their phenotype. Several genetic innovations have been made possible by the acquisition and "domestication" of these hereditary bacteria, which have, for example, played a critical role in the exploitation of several unbalanced diets, such as blood [1] or sap fluids [2]. In addition, a wide range of bacteria infecting insects induce reproductive manipulation, thanks to which they achieve high prevalence in populations despite bringing no benefit to their hosts [3]. Overall, the diversity of bacteria involved in these genetic innovations is now quite well known, thanks to decades of research efforts [2, 4, 5]. In some systems, factors shaping this global diversity have even been identified, with host species and host relatedness [6, 7] appearing as the most structuring ones (but see [8]), although ecology may matter for some specific bacterial lineages[9, 10].

On the contrary, and despite their ubiquity on earth, the diversity and potential phenotypic effects associated with heritable viruses are largely unknown. This gap in our knowledge is regrettable for several reasons. Firstly, because of the size of the gap. Indeed, recent developments in metagenomics, metatranscriptomics and large-scale database mining have uncovered thousands of unsuspected viral passengers [11–13], the majority of which are not known to be pathogenic to their host. Among them, there is no doubt that a significant fraction benefit from vertical transmission. Secondly, several case studies indicate that inherited viruses can play a key role in the functioning of organisms and ecosystems. For instance, some viruses can have conditional or long-term beneficial effects. Examples of conditional beneficial effect include the induction of thermal tolerance [14] or the protection against pathogenic viruses [15], while long-term beneficial effects can be illustrated by the independent cases of viral endogenization (integration into wasp genome) and domestication (retention by selection) observed in certain parasitic wasps [16]. In parasitic wasps, viruses are used as biological weapon to protect parastic eggs from the host immune response [17, 18]. Further, like bacteria, some heritable viruses can also manipulate the phenotype of their hosts in a number of ways, by interplaying with insect behaviour [19], [20] or insect physiology [21], with possible consequences at the community level [22].

In addition to our poor knowledge on virus diversity *per se* (except in very few model systems that have been deeply looked at, such as some mosquitoes [23] or the honey-bee [24]), the factors that may shape viral community structure, such as host species, space or time, are even less well identified. This may be partly explained by the fact that most of the studies exploring virus diversity do not provide a global unbiased picture of viral communities nested within the community of interacting organisms in which they live. Numerous studies focus on one or a very limited set of host species (for example: [25]), which may still enables testing of factors such as space and time [26, 27], but inevitably prevent testing the host species factor. Conversely, some recent large-scale studies using public databases have provided invaluable insights into viral diversity [11–13]. However, these findings only concern RNA viruses, as they are all based on homology searches using the universal RNA-dependent RNA polymerase (RdRp) gene as bait (a strategy that cannot be used to track DNA viruses, as they do not share a universal gene). In addition, the identification of factors shaping the viral communities described in these huge datasets was limited either because most viruses derived from metagenomic or environmental datasets thus preventing inference of host species [13], or because of a lack of information regarding the ecology of the species used [11, 12]. Altogether, these limitations impede our capacity to identify factors that may structure the community of viruses hidden within insect communities.

To study the structure of virus community, one requires a set of potentially interacting species in common geographical areas. Host-parasitoid communities nicely fits with this definition. In this paper, we studied the composition of viruses associated with Drosophilidae species and their parasitoids in two locations sampled during two consecutive years and two seasons. This insect community has been extensively studied in the past [28, 29] and a number of inherited viruses have already been described in one species or another [30], including some with fascinating phenotypic effects. In particular, a behaviour manipulating virus has been discovered in the larval parasitoid *Leptopilina boulardi*. This inherited virus forces females to accept laying eggs in already parasitized hosts (a behaviour called "superparasitism"), thus permitting the horizontal transmission of the virus between the parasitoids sharing the same *Drosophila* host [19, 20]. This behaviour manipulation allows the virus to reach high prevalence in wasp populations [31], and may contribute to the coexistence of *Leptopilina* species on the shared *Drosophila* ressource [22]. In our sampling, the community was composed of 9 *Drosophilidae* species that are potential hosts for six parasitoids, four attacking the larval stages, two attacking the pupal stage. By using a viral purification protocol combined with DNA and cDNA sequencing, we were able to identify both DNA and RNA viruses present in the host-parasitoid community. By applying this protocol to insects after two generations of rearing under laboratory conditions, this setup ensures that the viruses we are focusing on are transmitted along generations, either because of vertical transmission *per se* (transmitted through gametes), or because of pseudo-vertical transmission, for example through contamination of the medium in which the offspring develop. In addition, these two generations of rearing ensured that the viruses were capable of replicating in the insect species in which they were found, allowing us to confidently infer each virus host spectrum. This way, we identified a rich community of both RNA and DNA viruses and addressed the following questions: what factor(s) (or interaction of factors) structure the community of viruses? Is this geographical location, season, sampling year or species? In addition, we tested whether the relatedness of insect species influences the viral communities they host, similar to what has been regularly found for bacterial microbiomes. We did not include tests for ecological factors (i.e. environment, host range), as all the species were collected in the same environment (i.e. rotten fruit), and because the host range of most of the wasps is largely overlapping and still unclear for some wasp species (see table S1).

Next we also tested whether parasitoids have more heritable viruses than their hosts. This idea was motivated by the fact that throughout evolution, parasitoid wasps have maintained a special relationship with viruses. This is evident not only from the abundance and diversity of "free-living" heritable viruses that have been detected in the reproductive tract of parasitoid species [19, 32–35] but also from the abundance of endogenous viral elements found in the genomes of Hymenoptera with a parasitoid lifestyle [16]. These observations suggest that parasitoids are ideal hosts for viruses, perhaps because of their particular life cycle. Indeed, during oviposition, female parasitoids inject substances produced in their reproductive apparatus (typically in the venom gland) into their host [36]. Viruses could take advantage of this behavior to ensure their own transmission within the parasitoid population by targeting the tissues involved in the production of these substances [20, 35]. Localization in this tissue may not only favour vertical transmission along generations but also horizontal transmission in case of host sharing [20, 35]. In addition, because parasitoids necessarily develop from a hosts in every generation (whereas hosts are not attacked by parasitoids in every generation, and when they are, they usually die), we can speculate that parasitoids may occasionally pick up viruses from their hosts (whereas the reverse may be less frequent), ultimately increasing the corpus of viruses for parasitoids compared to free-living species. Finally, we also tested experimentally the transmission and phenotypic effect of one of the parasitoid viruses discovered in this survey. In total, our analysis uncovered 53 viruses, 37 of them being described for the first time. Thanks to the community-level approach, this dataset also provides valuable insight into the host range of previously described viruses, sometimes challenging previous interpretation made from monospecific datasets.

## 2 Material and methods

### 2.1 Sampling design and rearing conditions

Two sites (untreated with insecticides) were sampled in east/southeastern France (Igé, Burgundy; Gotheron, near Valence, fig. S1) for two consecutive years (2011 and 2012) in June and September. The climate of these two sites is quite different with Gotheron having a mediterranean climate, while Igé’s climate is continental. To collect the insects, 30 banana traps were used at each location and time. Ten traps were left for three to five days to collect Drosophilidae, 11 to 14 days to collect larval parasitoids, and 18 to 19 days to collect pupal parasitoids. Trapped insects were identified to the species level by visual inspection under a binocular microscope when necessary. As far as possible, for each species and each location and time point, 35 isofemale lines were established and maintained during two generations in the lab. All Drosophilidae species were reared using a standard medium [37] at 21C whereas all wasps were reared at 25C using *D. melanogaster* as host (StFoy strain). These two temperatures were chosen because they were satisfactory for the development of all insect species. In reality, the number of foundresses varied a lot between species (from 1 to 35). In order to maximize the quantity of material available in the end for sequencing, the second generation of lab rearing was also used to amplify the lines for the species having a reduced number of foundresses (below 35). For each sample, after these two generations of possible amplification, the insects emerging from all G2 vials (as far as possible 35) were pooled together and stored at -80°C in an RNA later solution with 0.2% Tween20 for later purification of viruses.

### 2.2 Purification of viral nucleic acids, barcoding and sequencing strategy

#### 2.2.1 Sequential Isolation of the Insect Genomic DNA and of the Viral Nucleic Acids

Adult insects stored at -80°C were crushed (between 80 mg to 4650 mg of insects depending on the species) with a “Tissue-Lyser” apparatus (Qiagen) in a buffer containing 20 mM Tris-HCl pH 7.5, 10 mM KCl, 4 mM MgCl2, 6 mM NaCl, 1 mM DTT and 0.1% Tween 20 v/v (buffer inspired from [38] and adapted to allow the activity of the DNase I). The crushed insects solution was filtered using syringes and filters until a 0.45 µm flow-through is obtained. Then an aliquot of 500 µl of the flow-through was taken to extract the insects genomic DNA by the Boom R. SiO2 technique [39, 40]. The insect genomic DNA was used as template for subsequent CO1-PCR. The remaining of the flow-through was treated 2 hours at room temperature with DNase I (0.5 units/ml, Sigma-Aldrich) and RNase A (400 mg/ml, Invitrogen) to remove unprotected nucleic acids. The putative viral nucleic acids (VNA) protected by their capsids from DNase I and RNase A digestion, were released from their capsid and purified using the SiO2 technique. At the end, the SiO2-eluate which contains a mixture of viral-DNA and viral-RNA is separated into 2 equal parts v/v. One part was treated by the Turbo DNA-free Kit (Ambion™) to obtain the “viral-RNA”; while the other part was treated by a RNase-A DNase-free (Fermentas) to obtain the “viral-DNA”.

#### 2.2.2 Cytochrome oxydase 1 - PCR

To check that insects have been correctly identified, a Cytochromoe Oxydase 1 (CO1) - PCR was performed on each insects genomic DNA using LCO1490 (GGTCAACAAATCATAAAGATATTGG) and HCO2198 (TAAACTTCAGGGT-GACCAAAAAATCA) primers with the AccuPrime™ Taq DNA Polymerase System (Invitrogen) with 30 or 34 PCR-cycles depending on the templates.

#### 2.2.3 Random-PCR amplification of the VNA

Random-PCR amplification of the Viral DNA or RNA were respectively achieved with the GenomePlex Complete Whole Genome Amplification (WGA) Kit and the TransPlex Whole Transcriptome Amplification (WTA) Kit (Sigma-Aldrich) according the manufacturer’s instructions, except that we used our own Amplification Master Mix, namely an AccuPrime Taq DNA Polymerase System (Invitrogen) with a 5’-phosphorylated universal primer (>PrimerUniversel_WG/TA: GTGGTGTGTTGGGTGTGTTTGGNNNNNNNNN). An equal volume (respectively 5 and 5.8 *µ*L for DNA or RNA extracts) for the various viral nucleic acids was used as PCR template and the number of PCR cycles was individually adjusted depending on the quantity of PCR products estimated from an agarose gel. Between 12 and 19 PCR cycles were needed for the DNA extracts, while between 13 and 20 PCR cycles were needed to equally random-PCR amplify the various samples.

#### 2.2.4 Parallel tagging, pooling and sequencing strategy

We used the “Parallel tagged sequencing” strategy of [41] in order to barcode the amplicons obtained from each CO1-PCR samples, each random-PCR amplification of viral-DNA samples (WGA) and each random-PCR amplification of viral-RNA samples (WTA; 55 of each, plus three negative controls = 168 samples in total, see file Genetic_construct.pdf on the github for details). In order to minimize mislabelling issues, all pairs of 8bp barcodes had at least three differences (median=6). The samples were pooled so that they had approximately the same number of molecules in the final pool used for the library preparation. The final sequencing was done on the ProfilExpert facility (Univ. Lyon 1) using 2 x 300pb Illumina paired-end reads. 11.88 Million pairedend reads were obtained.

#### 2.2.5 Demultiplexing

After quality check, reads were demultiplexed using cutadapt, allowing two mismatches in the 8bp tag. For CO1 reads, only reads starting with the expected primer sequences HCO (TAAACTTCAGGGTGACCAAAAAATCA) and LCO (GGTCAACAAATCATAAAGATATTGG) were retained (2 mismatches per full adaptor is tolerated without gaps and min_overlap=18), then trimmed on quality with stringent parameters to ensure high accuracy of insect species identification using trimmomatics (LEADING:20 TRAILING:20 SLIDINGWINDOW:10:15 MINLEN:220). After pairing reads using the python script fastqCombinePairedEnd.py, they were assigned at the species level using the Barcode of Life Data (BOLD) system. For the viral reads obtained after WGA or WTA, only those containing the expected Trans-plex/Omni-plex adaptor were retained (with 2 mismatches authorized without gaps and min_overlap=18). They were quality trimmed using trimmomatic v0.38 (LEADING:20 TRAILING:20 SLIDINGWINDOW:10:20 MINLEN:30 HEADCROP:10 CROP:250), and paired using the python fastqCombine-PairedEnd.py script. Singletons were also retained in downstream analysis. From the 11.88 millions reads, 5.12M contained a CO1 barcode, 4.16M contained a WGA barcode and 2.6M reads contained a WTA barcode. After all additional filtering steps, we obtained 38735 high-quality CO1 sequences (average=704 paired-reads per sample), 591257 paired-end + 2270517 single-reads for the WGA samples (average 6.7Mb per sample), and 349668 paired-end + 1660756 single-reads for the WTA samples (average 4.3Mb per sample).

### 2.3 Full genome sequencing of a new Filamentovirus in the wasp *L. heterotoma*

In the parasitic wasp *L. heterotoma*, a preliminary analysis of the metagenomic dataset revealed the presence of a set of 12 contigs related to the behaviour manipulating virus LbFV (25 to 48% identity at the protein level). Because LbFV is most likely the unique member of a new dsDNA viral family [42], and because of the originality of the phenotype affected by the virus (superparasitism behavior), we sought to obtain the complete genome sequence of this virus. To this end, we sequenced (without the WGA step), one of the samples positive for these 12 contigs (sample 43, Igé, Sept. 2012) using Illumina technology. The library construction and sequencing was done by MACROGEN company using the protocol for low-input material (Truseq Nano DNA library). 51 million of paired-end reads (150bp) of high quality (96.5 had a Phred-like score above 20) were obtained. After deduplication using fastuniq, reads were mapped to a database containing the genomic sequence of *L. heterotoma* and other insects of the community (*L. boulardi*, *D. melanogaster*, *D. simulans*, *D. subobscura*, *D. hydei*, *D. immmigrans*, *D. obscura* and *D. suzukii*) using hisat2, keeping only unmapped reads for the next step. After quality trimming using trimmomatic (v.0.38, parameters: LEADING:10 TRAILING:15 SLIDINGWINDOW:4:15 MINLEN:36), one sixth of the filtered dataset (i.e. N=8555009 paired-end reads) were assembled using MEGAHIT v1.2.9 (default parameters), ensuring around 1000x coverage on LhFV genome.

### 2.4 Inferring viral presence in each sample and each library type (wga/wta)

To quantify virus presence in each sample, a unique database on which reads could be mapped was required. To produce this database, we assembled the reads obtained from all WGA libraries on one side and all WTA librairies on the other side using MEGAHIT v1.2.9 (default parameters). During the preliminary analysis, we realized that WGA and WTA datasets were associated with large variation in coverage. Other experiments in the lab suggested that small quantities of DNA was sufficient to produce good quality Illumina sequences. Thus, we decided to directly construct a unique illumina library containing the pooled sample of all the purified nucleic acids obtained from all samples (except sample 43 used for LhFV sequencing) together with the product of their reverse transcription, without any amplification (neither WGA, nor WTA). This metagenomic sample (DNA/RNA) library was sequenced by MACROGEN company on an HiC machine and produced 211M high quality reads (96% with Phred-like score >20). After deduplication, host decontamination (mapping reads on the insect database as done for LhFV), and quality trimming, 192M reads were assembled using MEGAHIT using default parameters. Finally, the four assemblies obtained from the different analysis (wga, wta, metagenomic, LhFV) were merged using flye (with the –subassemblies option) in order to obtain a comprehensive view of the virome. This raw database contained 19604 contigs. Taxonomic assignation was performed by using the first hit of a mmseqs analysis performed on a local version of the refseq protein virus database augmented with additional virus proteins available in ncbi by querying "(virus[Title]) AND Drosophila " (2764 sequences), proteins from a few insect genomes (*Drosophila melanogaster*, *D. simulans*, *D. subobscura*, *D. obscura*, *D suzukii*, *Apis mellifera*, *Nasonia vitripennis*), and proteins from some bacteria and human (downloaded on the 12th may 2021, evalue threshold = 0.001).

The reads from each individual sample obtained after either whole transcriptome (with reverse transcription) or whole genome (without reverse transcription) amplification (WTA/WGA) were then mapped individually against this common database. Contig coverage and contig average sequencing depth were then calculated using samtools, and used to infer viral presence in each sample. 1358 and 281 contigs were respectively covered by the reads produced from at least one sample of, respectively, the wga and the wta datasets (a coverage threshold of 25% was used). After filtering out sequences assigned to non viral entities, or assigned to phages, transposons, polydnaviruses, to RNA viruses in the wga libraries or to DNA viruses in the wta libraries (since they are not expected there), while keeping unassigned contigs with high coding density (>50%, as expected for viral genomes), this led finally to 35 putative contigs for the WGA dataset and 183 for the WTA dataset. Raw reads have been deposited on the ncbi under the Bioproject PRJNA1000623, and the accession numbers associated with the assembled contigs are listed in tables S2 & S3 and are available as text files on our github.

### 2.5 Grouping contigs into putative viral genomes

We have tried to group contigs into single putative viral genomes for several reasons. Firstly, many viruses have a segmented genomic structure, which obviously necessitates clustering. In addition, we expected that at least some of the viruses present in our dataset would be poorly covered due to their low prevalence and/or low density in insects. This could lead to fragmented assemblies. To reach this goal, we used both phylogenetic information (contigs assigned to similar viruses are more likely to correspond to the same virus) and contig co-occurrence in insect samples (contigs that are always associated in the same samples are more likely to be part of the same virus). This grouping was done by eye inspection of the contigs heatmaps obtained after clustering contigs based on their co-occurrence in samples (fig. S2 & S3).

### 2.6 ORF prediction for virus genomes

We used getorf from the EMBOSS to predict ORFs with at least 50 amino acids (-minsize 150). We used two options for the -find option: valid ORFs should start with a methionine and finishing with a stop codon (find 1) or should simply be devoid of STOP codon (find 0). From all getorf ORFs, we wrote a custom R script ( orf_prediction.R available on our Github) that eliminates ORFs overlapping by more than 30 nucleotides while maximizing coding density (see file annotation.html on the github webpage). All orf predictions are given in the supplementary file S1.

### 2.7 Phylogenies

Orthologs of viral proteins of interest were retrieved from the mmseqs search evoked above. The sequences were aligned using muscle algorithm v3.8.31. Maximum likelihood phylogenetic reconstructions were then performed using PhyML (parameters:-daa-mLG-b-4-ve-c4-ae-fm). The branch supports were computed using approximate Likelihood Ratio Tests (aLRT).

### 2.8 A case study: prevalence, transmission and phenotypic effects for an Iflavirus infecting the parasitoid *L. heterotoma*

#### 2.8.1 Detection of viral infection

Six *L. heterotoma* isofemale lines originating from a different location (Lyon, France) were analyzed by rt-PCR for the presence of two viruses discovered during this study, namely Phasmaviridae_L.h (primer sequences: TGGCTGGGTTATTGGCACA & AGTTTGGGTGCTCTGTGAGA) and Iflaviridae_L.h (primer sequences: GGACGGGGTGCATTTGAATT, & ATCTGGACGATGCTGTACCC). The total RNA of single adult females was extracted using the RNeasy Mini Kit (QIAGEN) and DNAse treated following manufacturer’s instructions (Invitrogen, DNase I, AM 1906). After reverse transcription using the Kit SuperScript™ IV VILO™ (Invitrogen), the cDNA were used in a PCR assay (using DreamTaq from ThermoScientific, with an hybridization temperature of 59°C and 60°C respectively for Phasmaviridae_L.h and Iflaviridae_L.h). The validity of the PCR products (expected size respectively 191bp and 153bp) was checked by Sanger sequencing.

#### 2.8.2 Deciphering the vertical and horizontal transmission of the virus

In order to test the mode of transmission of the virus, we crossed Iflaviridae_L.h-infected and Iflaviridae_L.h-uninfected individuals (reared under the same conditions as the other parasitoids, described above). These individuals derived from two isofemale lines (3:infected and 38:uninfected). Four individuals from each parental line and from each of the F1s (female infected x male uninfected or female uninfected x male infected) were then checked by rt-PCR to infer their infection status.

In addition, we tested whether the virus could be horizontally transferred between parasitoids sharing the same hosts, through superparasitism. To do so, we offered young *Drosophila melanogaster* larvae to uninfected fertilized females during 24h. These supposedly parasitized hosts were then offered to 3 infected and unfertilized females during 24 additional hours. Because of the haplo-diploid sex determination system, this setup ensures that the emerging females are offspring of the uninfected fertilized egg-laying mother. After development, the infection status of emerging female wasps was checked either individually (n=4) or on a pool of 20 individuals to increase the detection power.

#### 2.8.3 Effect of the virus on phenotypic traits

The effect of the virus was measured on four important phenotypic traits: the parasitoid induced mortality (PIM), the rate of successful development or pre-imaginal survival (PS), the sex-ratio (SR) and the superparasitism behaviour. To measure these traits, couples of *L. heterotoma* (both infected, both uninfected or one infected and the other uninfected) were isolated for 24 h with a group of first instar *Drosophila* larvae hatched from 100 eggs. Four replicates of each parental line (infectedxinfected or uninfectedxuninfected) were performed, together with 8 replicates of each F1 cross (female infected x male uninfected or female infected x male uninfected). In addition, twelve controls without wasps were done the same way, in order to measure the natural mortality of *Drosophila*. After development, emerging adult *Drosophila* and wasps were collected every day, sexed and counted to estimate 3 parameters in each replicate: (1) Parasitoid Induced Mortality (PIM) is the estimated proportion of *Drosophila* killed by wasps, estimated by the difference between mean numbers of flies emerged from controls (without wasps) and from the experimental replicates (with wasps); (2) The pre-imaginal survival (PS) is the ratio of emerging wasps to parasitized hosts as estimated by PIM*100; (3) The sex ratio (SR) measured as the proportion of females among adult offspring.

To measure superparasitism behaviour, single or groups of 3 females (4 or 6 days old) were placed during 48h with ten first instar *Drosophila* larvae on an agar-filled Petri dish spotted with a small amount of yeast (at 22.5°C). After 24–48 h, two to three *Drosophila* larvae from each Petri dish were dissected and the number of parasitic immatures (eggs and larvae) counted. Superparasitism behaviour of each female was estimated as the mean number of immatures per parasitized host.

## 3 Results

### 3.1 Molecular confirmation of species identification

To check that our morphological identification of the different insect species was satisfactory, a CO1 PCR product was generated and sequenced for each sample along with the putative viral nuceic acids. Insect samples were correctly identified except for a few samples: one sample was taken as *L. heterotoma*, whereas the CO1 data indicated that this was *L. boulardi* (probably a labelling issue because these two species are easily distinguishable), 2 samples attributed to *D. phalerata* were in fact *D. immigrans* for one of them and *D. kuntzei* for the other, and finally the specimens identified as *D. subobscura* were in fact a mix of *D. subobscura* and *D. obscura* for two localities and the parasitoid wasps identified as *Asobara tabiba* were in fact a mix of *A. tabida* and *A. rufescens* (see supplementary file S2). The mapping of CO1 reads confirmed the presence of a single species in each sample (the average number of reads per sample was 697.7), with the exceptions of *D. subobscura*/*D.obscura* and *Asobara* species (*tabida*/*rufescens*). In total we obtained samples from 9 species of Drosophilidae and 5 species of Hymenoptera parasitoids, whose phylogenetic relationships is given in fig. 1. As far as possible 30 isofemale line of each species were established and maintained for two generations in the lab for each collection date (2 years and two seasons) and for each location in order to enrich for vertically transmitted viruses.

**Figure 1.**
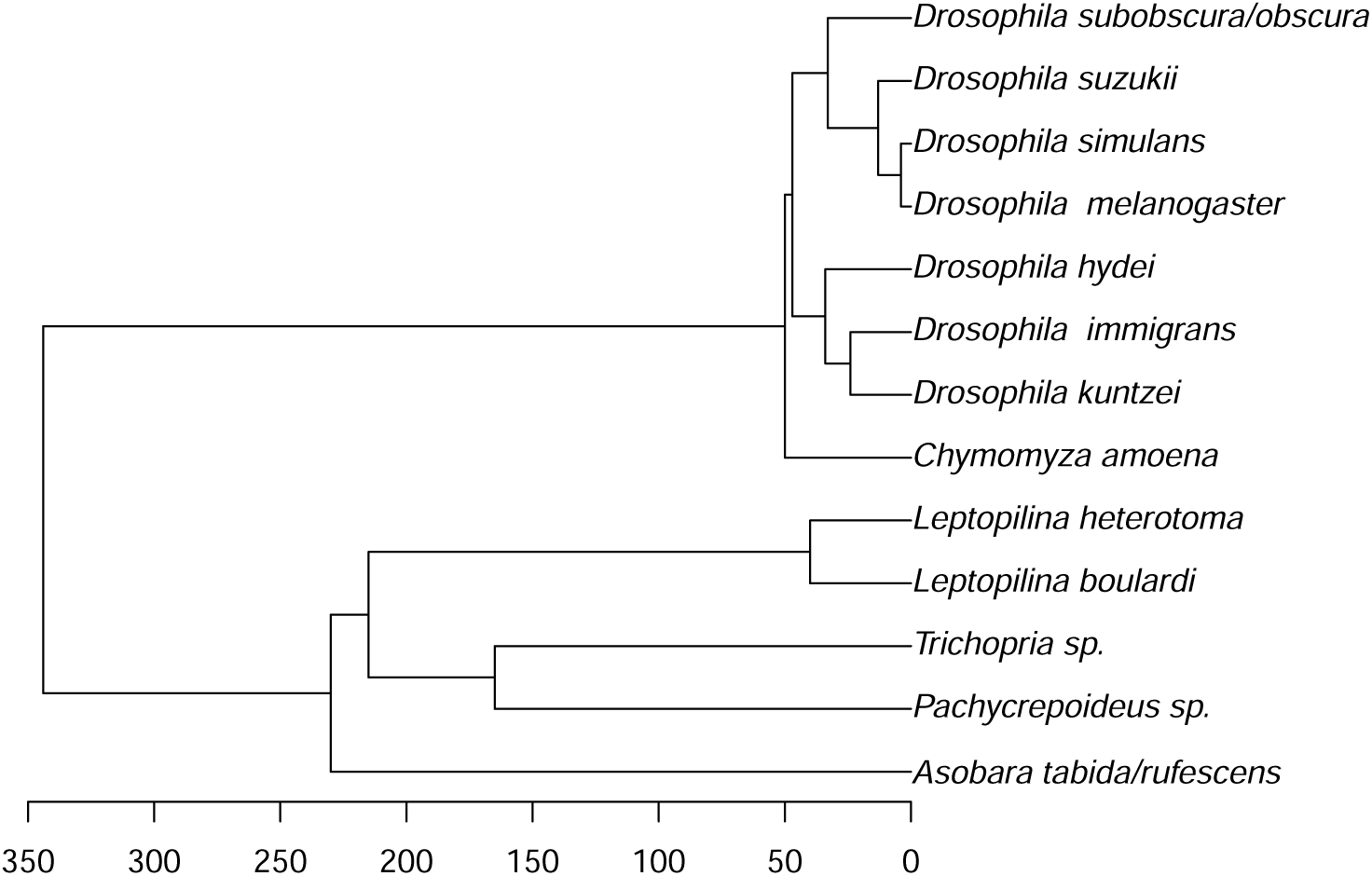
Phylogenetic relationships between wasps and Drosophilidae used in this study. The x-axis indicates the divergence time estimates in millions of years from present. The divergence times among Drosophilidae were obtained from [43] complemented by data from the database timetree for *D. kuntzei*[44]. For parasitoids, the divergence times were obtained from [45] complemented by data from [46] to estimate divergence between *Leptopilina* species. The divergence between the hosts (Diptera) and the parasitoids (Hymenoptera) has been estimated to be 344 million years, based on timetree estimates.

### 3.2 Overview of the viral diversity

From each sample (individuals belonging to a single species collected on a single date and a single location) and after two lab generations, insects were pooled and the putative viral nucleic acids were purified using a combination of filtration and enzymatic treatments. Purified nucleic acids were then split in two subsets in order to detect either DNA or RNA viruses. In total, we identified 47 and 192 contigs presumably belonging to DNA or RNA viruses respectively (Fig S2 & S3). Based on their co-occurrence in the same samples and on phylogenetic proximity, these contigs were then grouped respectively into 7 and 46 putative viruses with DNA or RNA genomes (Fig 2 & 3). Note that although the pipeline is designed to enrich for viral sequences, we cannot exclude that some of the contigs reported here derive from endogenized versions of the viruses. However, if this happens, we would expect the presence of “eukaryotic-like” sequences flanking viral sequences. To test this, we blasted (blastn) each of the DNA contigs against nt. None of the contigs revealed convincing evidence for eukaryotic sequence, with the exception of two contigs that may derive from endogenized versions (contig_9139 assigned to Reoviridae1_D.sub_obs and contig_22788 assigned to Vesantovirus_D.mel). The overall picture is thus that the contigs reported here are exogenous rather than endogenous sequences. A detailed description of the viruses detected in this study is given in Sections 3.6.1 and 3.6.2.

**Figure 2.**
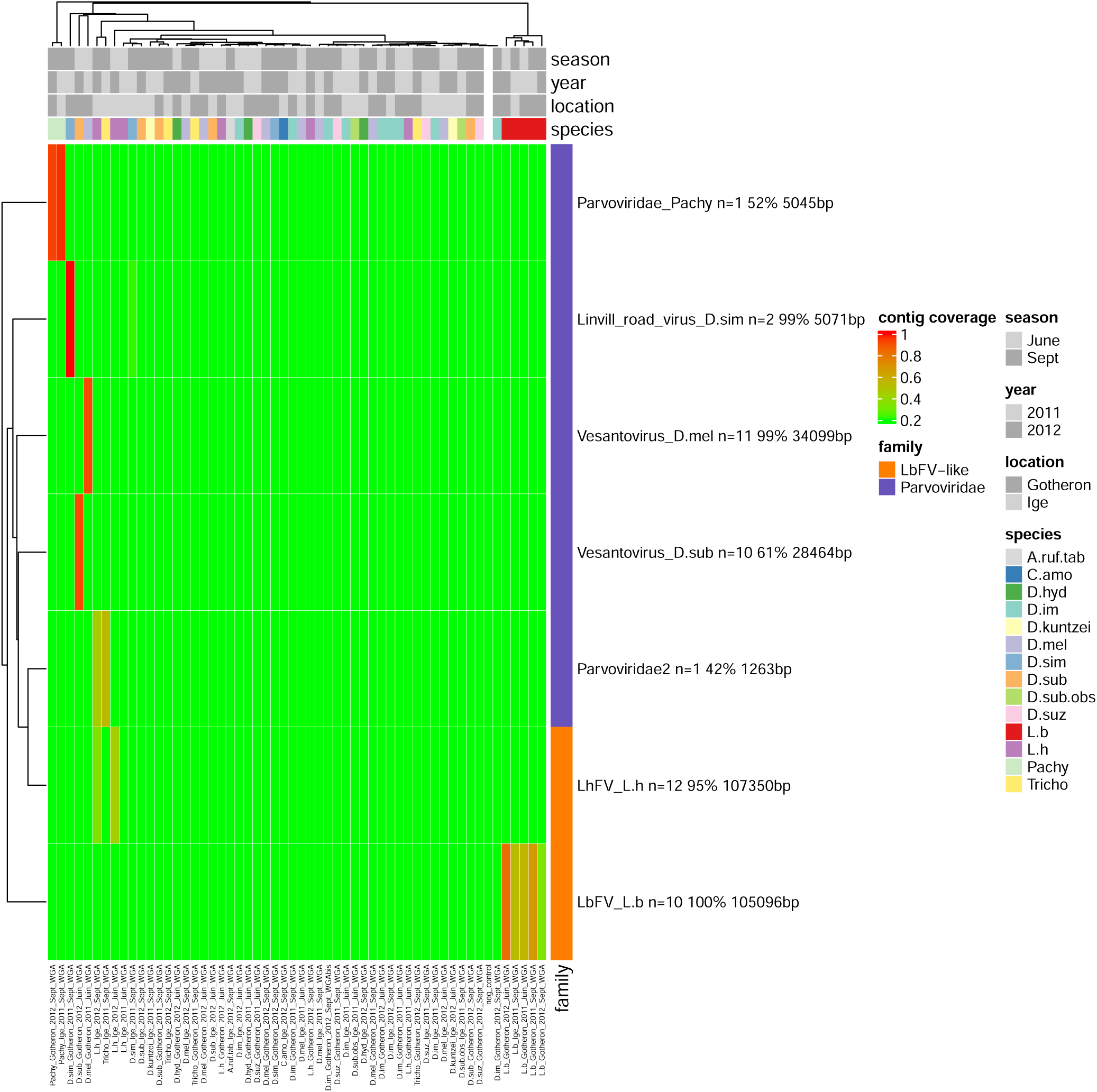
DNA viruses found in the *Drosophila*/parasitoid community. Rows correspond to the viruses, while the columns correspond to the 55 samples of insects from which viruses were detected. Both rows and columns were clustered using the Heatmap function default parameters (method “complete” based on euclidian distances, R package ComplexHeatmap) on both rows (viruses) and columns (insect species). Numbers indicated next to each virus name (n) indicates the number of contigs assigned to the virus. The percentage indicates the average percentage of identity with the closest proteins available in public databases. Total contig size is indicated next to % identity. Negative control is a water sample.

**Figure 3.**
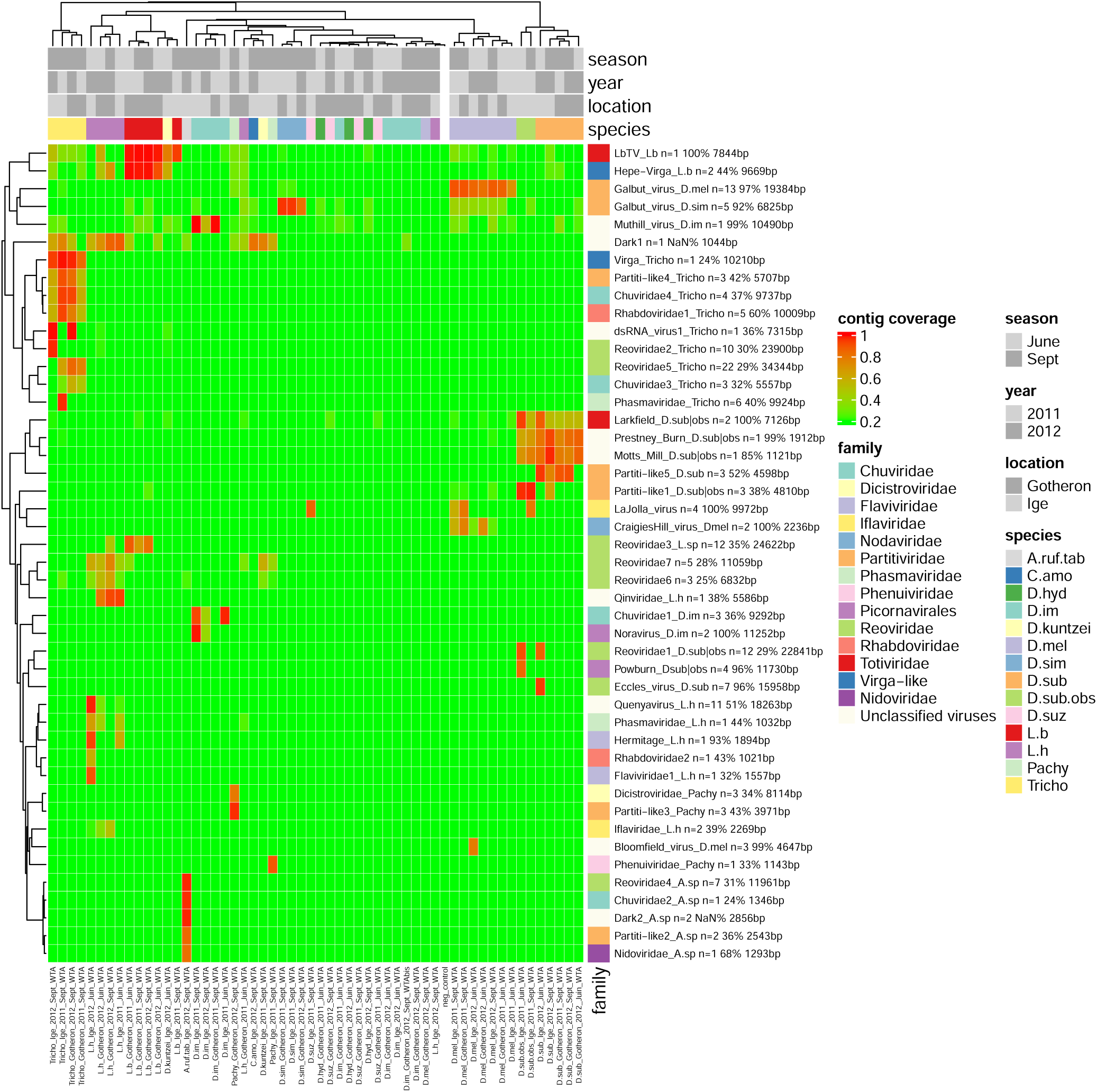
RNA viruses found in the *Drosophila*/parasitoid community. Rows correspond to the viruses, while the columns correspond to the 55 samples of insects from which viruses were detected. Both rows and columns were clustered using the Heatmap function default parameters (method “complete” based on euclidian distances, R package ComplexHeatmap) on both rows (viruses) and columns (insect species). Numbers indicated next to each virus name (n) indicates the number of contigs assigned to the virus. The percentage indicates the average percentage of identity with the closest proteins available in public databases. Total contig size is indicated next to % identity. Negative control is a water sample.

### 3.3 Factors structuring the viral community

First, we quantified the contribution of species, year, season, and location to the structuring of the viral communities. A sample was considered as being infected by a given virus if the mean coverage of the viral contigs was above 30%. Based on that 0/1 matrix of dimensions 55 (samples) x 53 (viruses), we then quantified the relative contribution of the different factors to the overall virus structure using variance partitioning analysis. The full model was highly significant (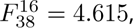 P<0.001) indicating that some of the factors included in our analysis structured part of the virus community. We then tested individual effects of each factor by performing permutation tests under reduced model. All factors were insignificant (all p-values > 0.33) at the exception of the host species which was highly significant (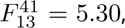 P<0.001) and explained 52.5% of the variance in virus distribution (Fig. 4).

**Figure 4.**
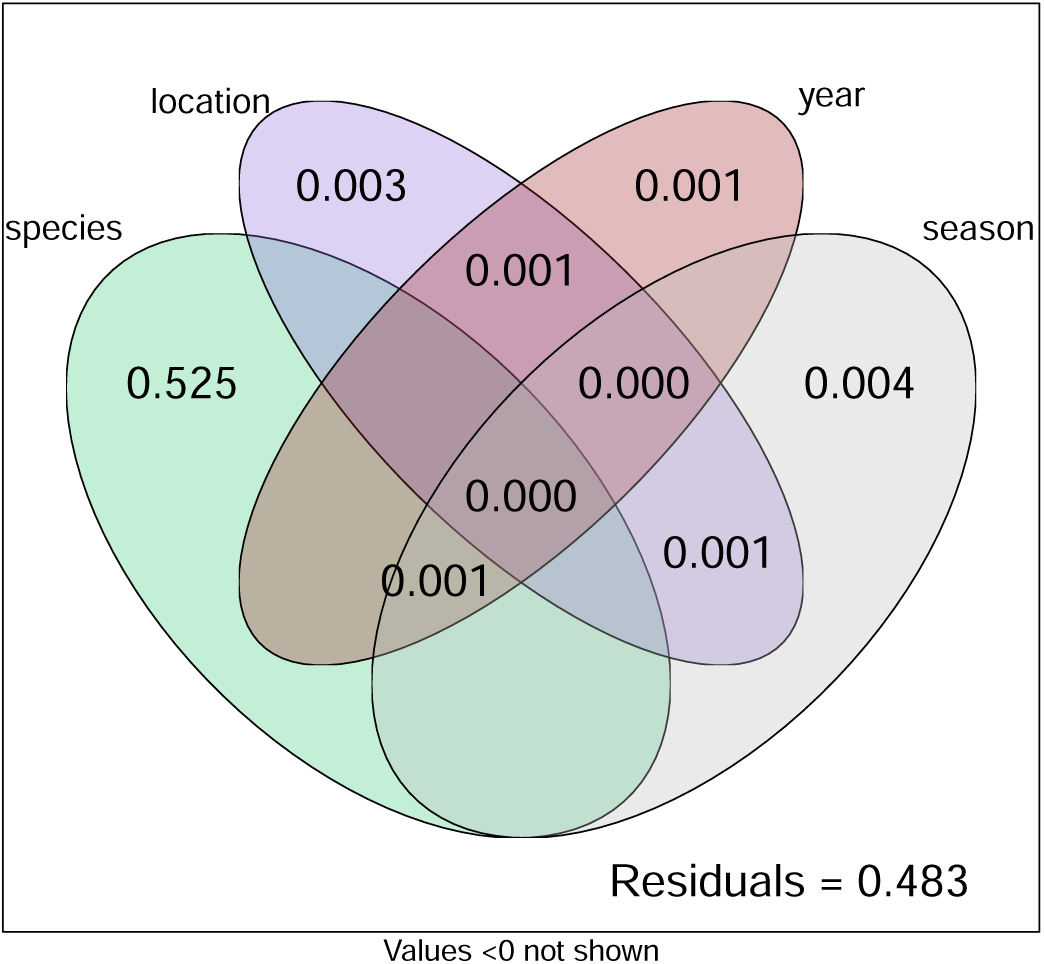
Results of the variance partitioning analysis showing the major contribution of host species on virus distribution. The numbers indicate the proportion of variance explained by each factor (based on the adjusted R square statistic which accounts for the difference in degree of freedom between factors[47]). When the adjusted R square method produced negative values, they were rounded to zero.

### 3.4 Insect relatedness does not correlate with virome composition

We then tested whether more closely related species had more similar virus communities than distantly related species. This was tested by correlating the genetic distance between all pairs of species (measured by time divergence) with the euclidian distances between the viral communities they harbour (insect species were considered as host for a given virus as soon as one of the sample was positive, i.e. coverage >30%). This correlation was performed with a Mantel test (using pearson, spearman and kendall correlation methods). None of the tests performed were significant (all p-values>0.20) indicating no significant effect of insect phylogeny on virome composition. Similar results were obtained by splitting the dataset between parasitoids and Drosophilidae and/or by using other distance for measuring viral communities dissimilarity (i.e. Bray-Curtis).

### 3.5 Parasitoids harbour more heritable viruses than their hosts

Since insect species was the main driver of virus presence, we associated viruses to one or several insect species based on virus contig coverage: an insect species was considered as a valid host as soon as the reads obtained from at least one pooled sample led to a viral contig(s) coverage above 30% (see Fig. S4). We then simply calculated the number of viruses infecting each insect species and tested whether host and parasitoids had the same propensity to host viruses (Fig. 5). We found that parasitoid species were hosts to more viruses compared to their hosts (Wilcoxon test, W = 7, p-value = 0.04479). The same conclusion holds if the samples *D.subobscura* and *D.subobscura*+*D.obscura* are merged together (W = 5.5, p-value = 0.03987). Because the number of samples or number of isofemale lines sampled differed among species and may mechanically impact the number of viruses discovered, we controlled for this by decomposing the variance in virus per insect species according to (i) the number of pooled samples or isofemale lines screened and (ii) the lifestyle (free-living/parasitoid). This was done both in a regular anova framework and in a glm framework with a poisson error distribution. All four analysis led to the same conclusion that parasitoids are infected by a higher number of viruses compared to their hosts (controlling for number of pooled samples: anova 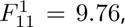 p-value=0.0097, glm: Estimate=0.95379, z-value=4.41, p-value=1.03e-05; controling for number of isofemales sampled: anova 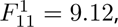 p-value=0.01164, glm: Estimate=0.936, z-value=4.41, p-value=1.44e-05).

**Figure 5.**
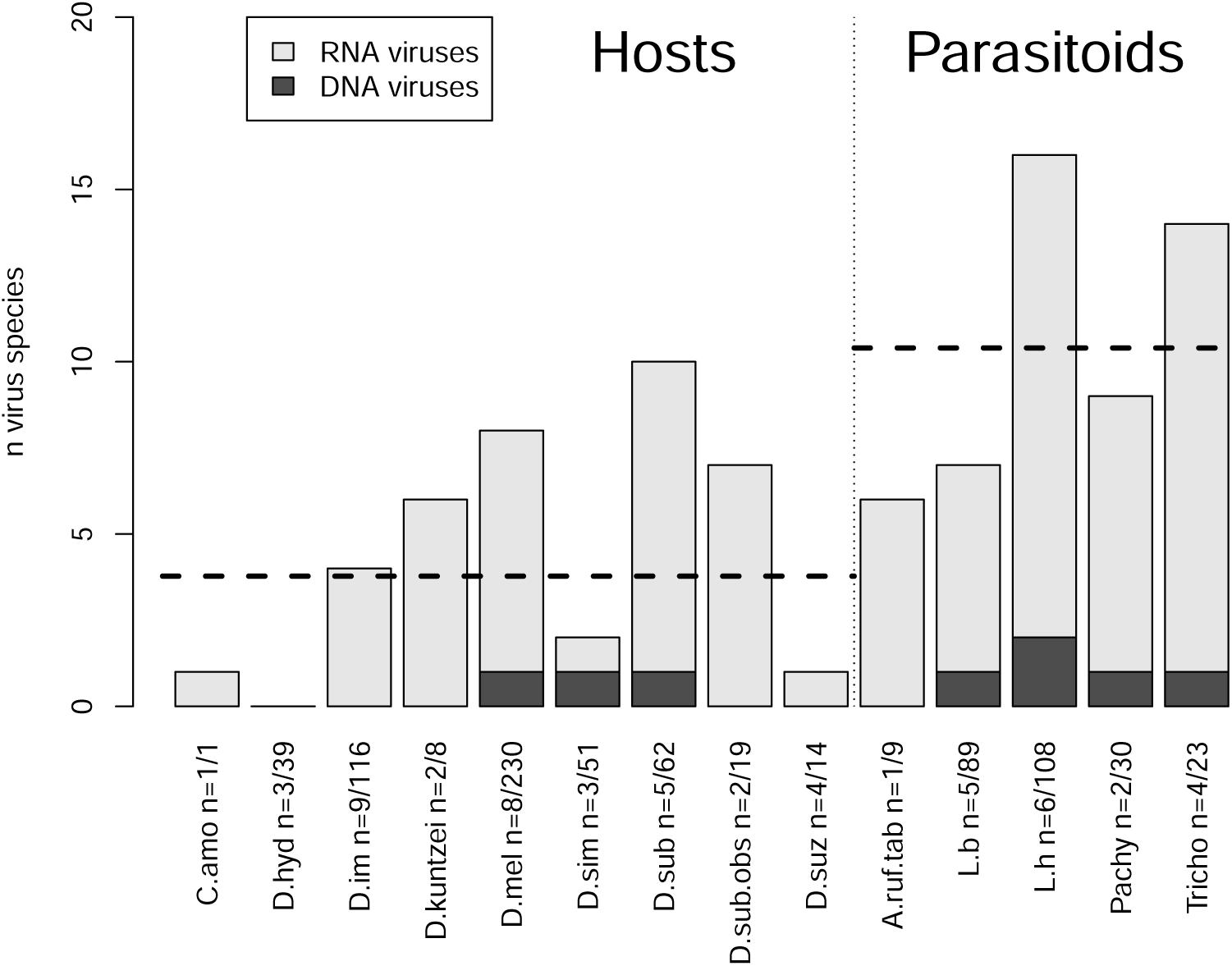
Parasitoids had a higher load of virus species compared to their hosts. The total number of virus detected was plotted for each insect species (dashed line represents the mean for hosts and parasitoids). The number of pooled samples (x1) and of isofemale lines (x2) analyzed is indicated next to each insect species names (n=x1/x2). Note that the virome is dominated by RNA viruses.

### 3.6 Description of the viruses

To identify the insect species contributing the most to viral dynamics, we performed a clustering analysis based on the number of reads obtained from each insect species (averaged per isofemale line sampled). This clustering led to the identification of a main single driver species for most viruses, in line with the results presented above (fig. 6). The results were overall highly consistent with the previous analysis relying on coverage clustering (fig. S4) but the picture was clearer using the number of reads. We thus used this vizualisation to organize the presentation of the different viruses identified in the community. This was especially useful for the discussion of the numerous RNA viruses. Readers who are primarily interested in general patterns, rather than the detailed composition of the viral community, may wish to skip to section 3.7.

**Figure 6.**
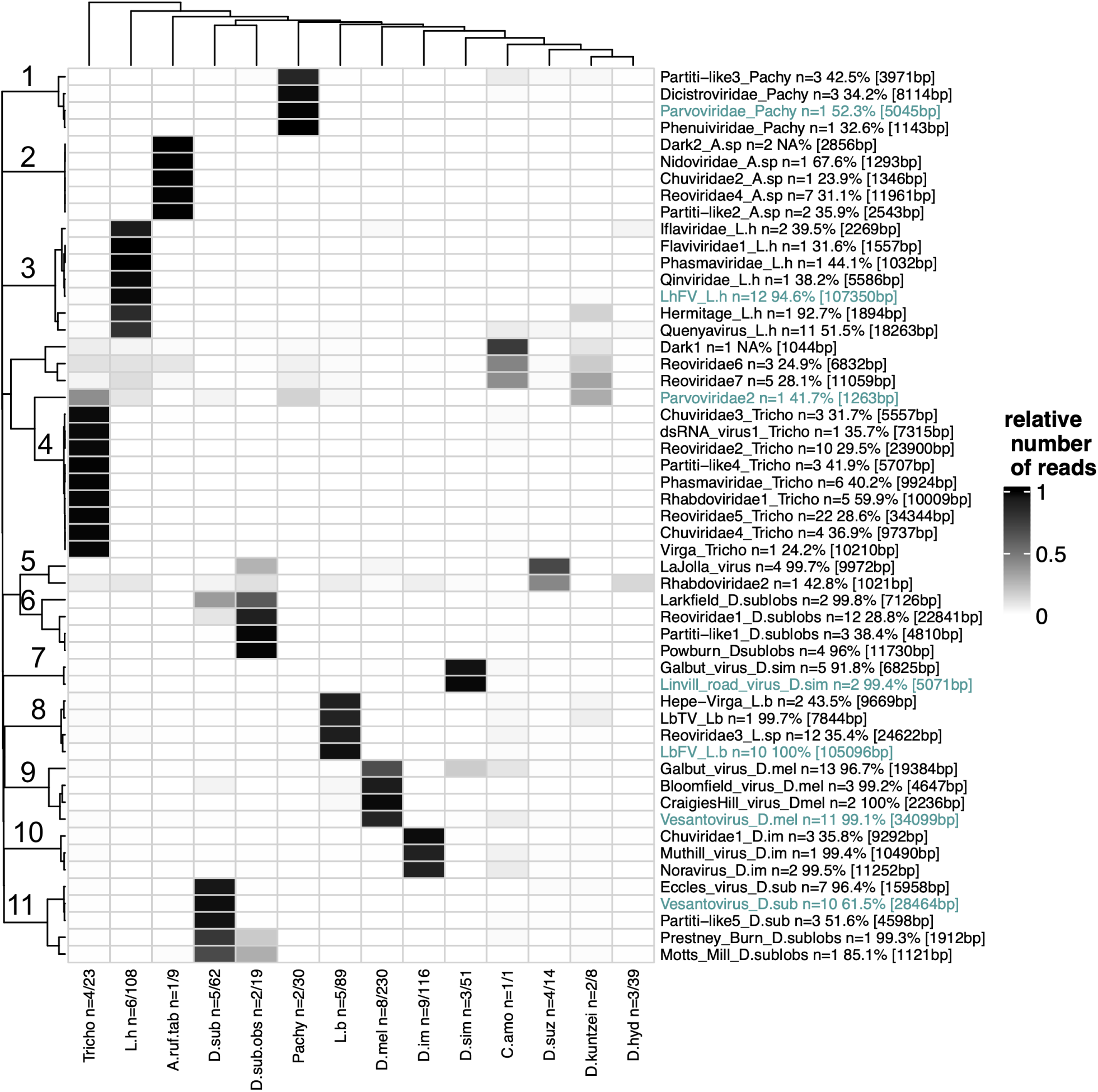
Virus distribution in the *Drosophila*/parasitoid community, grouped by host species, as measured by the average number of viral reads produced per isofemale line sampled, normalized by rows. The number of pooled samples (x1) and of isofemale lines (x2) analyzed is indicated next to each insect species names (n=x1/x2). Next to each virus name are indicated the number of contigs assigned to the virus, the percentage of identity with the first hit (amino-acids) and total length of the contigs in square brackets. RNA viruses are indicated in black, whereas DNA viruses are in green. Clustering method was performed using the Heatmap function default parameters (method "complete" based on euclidian distances, R package ComplexHeatmap) on both rows (viruses) and columns (insect species).

#### 3.6.1 DNA viruses

Seven DNA viruses were identified in this host-parasitoid community (Fig. 6). From our previous work, we expected the presence of the maternally inherited DNA virus LbFV in the wasp *Leptopilina boulardi*. This virus induces a behaviour manipulation on infected females by increasing their propensity to superparasitize, i.e. lay eggs in already parasitized larvae. Since superparasitism conditions permits the horizontal transmission of the virus between parasitoid lineages [19], the virus reaches high prevalence in wasp populations[31]. As expected, we found LbFV in all five *L. boulardi* samples encompassing the two sampling years, the two locations and the two seasons (fig. S2). Because our dataset is only composed of Illumina short-read sequences, the virus was split in 10 contigs due to the presence of repeated regions (hrs) in the genome of LbFV [42]. The total length of the LbFV-contigs obtained from this study was 105096bp which is very close to the expected size of the genome (111.5Kb). The percent identity with LbFV proteins was, as expected, close to 100%. This virus was exclusively found in *L. boulardi*, thus confirming its specificity [22].

Interestingly, in two samples of the related parasitoid *L. heterotoma*, we found a set of 12 contigs related to LbFV (25 to 48% identity at the protein level, see fig. S2). Surprisingly, most of these contigs also had high nucleotidic similarity (97.98%) with a virus recently discovered from a sample of pool-sequenced wild *D. melanogaster*, sampled in Spain (so-called Drosophila melanogaster Filamentous Virus, [48]). Our dataset revealed 1444 reads that mapped uniquely to these 12 contigs. Among them 1431 (99.1%) derived from only two *L. heterotoma* samples; the other 12 reads were found in 12 samples with a maximum of only 1 read mapped on a single sample. These residuals mappings were thus considered as spurious and are not considered further. In conclusion, our result rather suggests that the major host of this virus is actually the parasitoid *L. heterotoma* rather than *Drosophila melanogaster*. One hypothesis explaining the result obtained by Wallace and colleagues is either that this virus may occasionally infect the host of *L. heterotoma* without being able to spread efficiently, or that their sample was somehow contaminated by the *L. heterotoma* virus. Explanation for this possible contamination may be that one of the field-collected adult flies used in the analysis of Wallace and colleagues had been attacked by an infected *L. heterotoma* but survived infestation, as it is sometimes observed in the field [49]. In this later case, the virus would have been injected into the *Drosophila* during oviposition and survived until adulthood. This situation appears to be rather rare since DmFV was only found in a single pool-seq library out of 167 pool-seq samples (each with 30 flies), collected over 3 years from forty-seven different locations across Europe [48].

In order to obtain the full genome sequence of this virus, and because Whole Genome Amplification (which was used in our initial protocol) may induce coverage biases, we constructed an Illumina library from one of the two *L. heterotoma* positive samples (sample 43) without applying this WGA step. The sequencing data confirmed and extended the sequences obtained initially with the WGA step. With this additional dataset, we obtained 12 contigs. Sequencing depth for the 9 large contigs was homogeneous and high (mean=1181x, min=1148x, max=1302x), whereas the three small contigs ("contig_19696" "contig_19153" "contig_21206") had lower sequencing depth (mean=143.32x, min=45.35x, max=230.27x). In total the 12 contigs contained 107350bp, which is very similar to the genome size of its closest relative LbFV (111.5Kb). Our automatic pipeline predicted 105 ORFs (see suppl. file S1) which is also close to the number of proteins encoded in LbFV (n=108). Altogether, these data suggest that the 12 contigs contained the whole genome sequence of a virus infecting *L. heterotoma* that we propose to name LhFV instead of DmFV. A detailed description of this virus and other LbFV-related viruses will be given elsewhere ([34]).

We found two related viruses belonging to the Parvoviridae family in two *Drosophila* species. One of them, here denominated Vesantovirus_D.mel, was found in a *D. melanogaster* sample and was almost identical to the Vesanto virus discovered by [48] in the same species (Fig. S2). It was composed of 11 segments, 10 of which had high sequence homology with the Vesanto virus previously detected by [48]. Additionally, contig_8677 which showed strict association with the 10 other segments was most likely an additional segment of this Vesanto virus but without homology in public databases. This 1.8kb segment had high coding density as is expected for a viral genome (Fig S6). Note that one of the contigs assigned to this virus (contig_22788), may derive from an endogenized version of the virus as its 1020 first bp showed 99% identity with a retrotransposon gag protein from *Drosophila*. Alternatively, the *Drosophila* retrotransposon may have jumped into the viral DNA.

A second Vesantovirus (here denominated Vesantovirus_D.sub) was detected in *D. subobscura*. This new Vesantovirus was composed of ten segments. Although two of them (contigs 7654 and 17519) had no sequence homology with any known virus, they did show strict association with the other contigs and had high coding density (Fig S5), suggesting that they belong to the same virus. The phylogeny based on NS1 protein (ORF 940-2433, contig_2659) revealed that the two Vesanto viruses found in *D. melanogaster* and *D. subobscura* form a well supported monophyletic clade (Fig S7).

Another Parvoviridae, referred to here as Parvoviridae_Pachy, was also detected in two samples of *Pachycrepoideux dubius* (Fig. S2). The corresponding contig is 5kb long which fits well with the expected genome size of related densovirus [50]. In addition, we found the typical inverted terminal repeat (ITR) at the 3’end (but not at the 5’ end) suggesting that the genome is almost complete (). The genome is predicted to encode seven proteins (Suppl. file S1), and the phylogenetic analysis based on NS1 protein (positions 1853-2266) suggests that it is related to Diaphorina citri densovirus [50] and Bombus cryptarum densovirus [51] (Fig S8). Of note, Diaphorina citri densovirus is vertically transmitted along generations of *Diaphorina citri* through strict maternal transmission. Similar transmission mode may be at play in *Pachycrepoideus sp.*, possibly explaining its maintenance during two generations in our experimental setup.

Another new Parvoviridae (here denominated Parvoviridae2) was detected in *Trichopria sp.* and *D. kuntzei* but also in *L. heterotoma*, *Pachycrepoideus sp.* and *D. subobscura*. This 1263bp contig encoded a single orf (positions 2-1219) and is most likely a fraction of a larger virus genome, since ambidensoviruses are expected to have a 6kb genome. The phylogeny built using the encoded NS1 protein (Fig S8) showed a monophyletic clade containing this protein together with Ambidensoviruses detected in the feces of Cameroonian fruit bats [52]. Interestingly, the phylogeny built from NS1 protein of Parvoviridae (Fig S8) revealed that four sequences from ticks were nested within the viral diversity, suggesting that an horizontal transfer occurred from Parvoviridae to Ticks.

We found two contigs sharing high sequence similarity (*>* 99%) with the previously described Linvill road virus (Fig. S2). This virus was previously described in *D. melanogaster* by [48]. The authors noticed that the abundance of reads mapping to Linvill road virus was positively correlated with the level of *D. simulans* contamination, suggesting that infection by this virus may in fact better correspond to a spillover from *D. simulans*. Consistently with this hypothesis, we found that Linvill road virus was found almost exclusively in *D. simulans* with 99% of the reads mapped to *D. simulans* samples (n=11215 reads), with only marginal signatures of infection in *D. melanogaster* (with a maximum of 5 reads per sample, see fig. 6). This result suggests that the major host for this virus is rather *D. simulans*.

#### 3.6.2 RNA viruses

Because the dataset for RNA viruses was large, we choose to describe them by modules of viruses showing similar distribution among insect species. These modules were defined by the unsupervised clustering analysis and are indicated by a number in fig. 6. Eleven such modules were identified this way. The dynamics of a virus included in a module was mainly driven by a single insect species (that produced the great majority of reads), although other species may sill present signs of infection (fig. S4). Drosophilid viruses are discussed first, followed by parasitoid viruses. The viruses not included in any cluster are presented in the last part.

###### Drosophilidae modules

**Module 9: viruses mainly driven by *D. melanogaster***

Three RNA viruses appeared to be mainly driven by *D. melanogaster*. All three had been previously described from this same species. Our study thus confirms their abundance in *D. melanogaster*. We found 13 contigs corresponding to the previously reported Galbut virus (99% identity, Galbut_virus_D.mel in our notation). This virus was described as infecting *Drosophila melanogaster* and benefits from efficient vertical transmission by both males and females [53]. In addition, our data suggests that Galbut virus is able to infect the pupal parasitoid *Pachycrepoideus sp.* (fig. S4), although the great majority of reads was found in *D. melanogaster* samples (see fig. 6). Two previously described *D. melanogaster* viruses, namely Bloomfield and Craigie’s Hill viruses, were detected [48, 54].

**Module 7: viruses mainly driven by *D. simulans***

A small sized-module composed of two viruses was detected in *D. simulans*. This module contained one DNA virus (Linvill_road_virus, described above) and five contigs assigned to the Galbut virus first discovered in *D. simulans* (>92% identity) [54]. In our sampling, this virus, named Galbut_virus_D.sim, was detected in *D. simulans* but also in *D. melanogaster* and the parasitoid *L. heterotoma* (fig. S4). However, the great majority of reads detected for this virus were found in *D. simulans* suggesting that *D. simulans* is the main host for this virus (fig. 6).

**Module 6 and 11: viruses mainly driven by *D. subobscura* and/or *D. obscura***

These two modules were composed of 8 RNA viruses.

We found two contigs (5915bp and 1211bp) having 100% protein similarity with the Totivirus called Larkfield virus that was initally detected in *D. suzukii*[25]. Our data revealed no infection in the four *D. suzukii* samples (corresponding to 14 isofemale lines) but infection was detected in *D. subobscura* and/or *D. obscura* suggesting that the presence of the virus in *D. suzukii* could be the consequence of a spillover from *D. subobscura* or *D. obscura*.

A set of 12 contigs were assigned to a reovirus (Reoviridae1_D.sub). The RdRp encoded by contig_2830 aligns from positions 25 to 1402 with RdRp of Rice gall dwarf virus (ABF67520.1) which is 1458aa long, suggesting that this segment is complete or almost complete (fig S18). Because reoviruses have typically 9- to 12-segment dsRNA genomes, it is likely that this set of 12 contigs represents the full genome of a new reovirus specific to *D. subobscur*a and possibly to *D. obscura*. Note that the first 88bp of contig_9139 showed 98% identity with some uncharacterised *Drosophila* genomic sequences.

Three contigs presumably belonging to a Partitiviridae (Partiti-like1_D.sub) were co-occurring within *D. subobscura* samples (fig. S3). Two of them have sequence similarity with Vera virus (from 31 to 48% at the protein level) including one encoding an RdRp (fig S24). The third contig has sequence similarity with Chaq virus from *D. melanogaster*. It is unclear whether this contig belongs to the same virus or not.

As described by [55], we found evidence of infection by Powburn and Prestney Burn viruses in *D. subobscura* samples (96% and 99% protein identity).

Seven contigs with almost 100% identity with the reoviridae Eccles virus were detected in a single sample of *D. subobscura* (fig. S3). Eccles virus was first reported by [25] in *D. suzukii*. Our result thus indicates that this virus may also infect *D. subobscura*.

A virus displaying homology with Partitiviridae was detected in *D. subobscura*. This virus appears to be specific to this species, both in terms of contig coverage (fig. S4) and read numbers (fig. 6) and was designated Partitilike5_D.sub (see phylogeny in fig. S24).

Additionally, we found sequences related to the *Drosophila melanogaster* Motts Mill virus [54] in *D. subobscura* samples (85% protein identity, see phylogeny in fig. S17).

**Module 10: viruses mainly driven by *D. immigrans***

This module contained three RNA viruses, two of them were already identified as infecting *D. immigrans*, i.e. Nora virus, as previously found by [56] and Muthill virus (99% identity) by [55].

In addition, we found three contigs totalizing 9292bp with approx. 35% protein identity with Hubei chuvirus-like virus 3. The phylogenetic reconstruction built on RdRp indicated that this virus was part of the Chuviridae family (see Fig. S25) and was thus called Chuviridae1.

**Module 5: viruses mainly driven by *D. suzukii***

Two RNA viruses composed this module. La Jolla virus was already described by [54] and [25]. It is very prevalent in *D. melanogaster*, also found in *D. simulans* and *D. suzukii* [25, 54]. In our dataset, signs of infection were evident from *D. subobscura* or *D. obscura* and *D. melanogaster* (fig. 6 & S4).

In addition, a 1021bp contig showed protein sequence similarity (43%) with the Hubei dimarhabdovirus virus 2 (here called Rhabdoviridae2). The phylogenetic reconstruction based on the nucleoprotein confirms this relationship within the Rhabdoviridae family (fig. S14) and shows a proximity with sigma viruses detected in *D. immigrans* and *D. obscura*. Although most of the reads were detected from *D. suzukii* samples, we also detected signs of infection from several *Drosophila* species (*D. hydei*, *D. immigrans*, *D. subobscura/obscura*) and a few parasitoids (*L. heterotoma*, *L. boulardi* and *Trichopria sp.*, fig. 6 & S4)

###### Parasitoid modules

**Module 8: viruses mainly driven by *L. boulardi***

This module contained three RNA viruses.

We obtained two contigs (8247bp and 1422bp) of a virus distantly related (appr. 40% amino acid identity) to the Saiwaicho virus, that we refer to as Hepe-Virga_L.b. Saiwaicho virus was detected in wild collected *D. suzukii* [25]. Its expected size is around 10kb, suggesting that we obtained the complete or almost complete sequence, despite being split in two parts. Similarly to what they found, the polyprotein encoded by contig_22560 contains the following domains: methyltransferase, Ftsj-like methyl transferase, helicase and RdRp. The second ORF from this contig is homologous to the hypothetical protein AWA82267.1 [Saiwaicho virus] and the largest ORF from contig_9042 shows homology to the hypothetical protein AWA82265.1 which contains a conserved domain (pfam16504: Putative virion membrane protein of plant and insect virus). We built a phylogeny based on the RdRp domain only (fig. S21). This virus was found in *D. kuntzei* and *C. amomiza*, and in the parasitoids *Trichopria sp. Pachycrepoideus sp.*, *L. heterotoma* and *L. boulardi* (fig. S4). The great majority of reads were produced from *L. boulardi* samples suggesting that this species is the main driver for this virus (see fig. 6).

We found a 7844bp contig corresponding to the previously described Totivirus (LbTV) reported from *L. boulardi* [57]. In our analysis, although the great majority of reads was indeed detected in *L. boulardi* confirming that *L. boulardi* is the main host for this virus (see fig. 6), other species appear to be positive for LbTV (*Trichopria sp*., *D. subobscura*, *Pachycrepoideus sp.*, *L. heterotoma*, *D. kuntzei* and *D. melanogaster*).

A reoviridae (here called Reoviridae3_L.sp) related to Cimodo virus was detected in *L. heterotoma*. Cimodo virus has a segmented genome (around 12) with segments ranging from around 500bp to 4kb. Here we found 11 segments, ranging in size from 1kb to 4kb, including one without homology to known sequences. This contig was included as it co-occurs with the other 10 segments (fig. S3) and contains a large ORF (fig. S9). The phylogeny built on RdRp (Fig. S18) suggests that this virus is related to Cimodovirus found in African mosquitoes [58]. It has been suggested that Cimodo virus defines an as-yet-unidentified genus within the subfamily Spinareovirinae within the Reoviridae family [58]. Although most of the reads originated from *L. boulardi* samples (fig. 6), the related *L. heterotoma* showed signs of infection (fig. S4).

**Module 3: viruses mainly driven by *L. heterotoma***

This module contained six RNA viruses. Two contigs (respectively 1127 and 1142bp long) showing similarity (approx. 40%) with Formica exsecta virus were detected (here called Iflaviridae_L.h). Formica exsecta virus has a 9160bp genome encoding a polyprotein [59]. The genome we got here is thus probably incomplete. In the absence of RdRp in our dataset, we built a phylogeny based on the capsid domain (Fig. S13) which further confirmed the phylogenetic proximity of this virus with Formica exsecta virus and other Iflaviridae, including Deformed wing virus and Varroa destructor virus 1.

We detected a 1557bp contig showing similarities with Takaungu virus (32%). This virus was first detected in *D. melanogaster* samples [55]. This virus is poorly known and belongs to unclassified Flaviviridae. No phylogeny was built since our pipeline detected homology with only one sequence.

We named Phasmaviridae_L.h a partial genomic sequence related to Ganda bee virus (protein identity of 44%). Ganda bee viruses are segmented viruses (typically 3 segments around 2.2kb, 6.7kb and 2.8kb). The phylogeny built on the nucleoprotein confirmed the proximity with Ganda bee virus and other Phasmaviridae detected in Hymenoptera (fig. S10). Most viruses detected in Hymenoptera including the virus we found in *L. heterotoma* did form a relatively well supported clade (0.91 aLRT support), suggesting a specialization of this clade of Phasmaviridae on Hymenoptera (fig. S10). Interestingly, several insect sequences were nested within the viral diversity, suggesting that this viral gene has been endogenized in several insect genomes. Three Hymenoptera sequences (*Dufourea novaeangliae*, *Osmia bicornis* and *Bombus terrestris*) were found within the clade of viruses infecting Hymenoptera. Of note, a sequence apparently endogenized in the parasitoid *Leptopilina heterotoma* was detected but this sequence was not in the clade containing Phasmaviridae_L.h and most viruses associated with Hymenoptera. In addition, the phylogeny included a large monophyletic clade of several *Drosophila* species suggesting that some horizontal transfer may have occurred between Phasmaviridae and some ancestral *Drosophila* hosts. Notably, the topology of *Drosophila* sequences within this clade mirrors the known phylogeny of *Drosophila* species, suggesting that a single or a few events of transfer followed by vertical transmission (and an ancestral duplication) may explain the pattern. Within this *Drosophila* clade, two subclades were identified. Interestingly within each subclade, the sequences of species clustered according to their expected group and subgroup. For instance, D. persimilis, D. pseudoobscura, D. miranda, D. subobscura, D. guanche and D. obscura which all belong to the obscura group do form a well supported clade in both subclades of the tree. Additionnally, within this obscura clade, the sequences clustered according to the known subgroups (D. persimilis, D. pseudoobscura, D. miranda in the pseudoobscura subgroup and D. subobscura, D. guanche and D. obscura in the obscura subgroup). At a larger scale, we also observed the expected relationship among groups within each subclade: melanogaster group and obscura group did form a monophyletic clade (if we exclude *D. setifemur* sequence which is not well resolved) sister to a clade composed by (((repleta+virilis)+grimshawi)+immigrans). In addition, the basal species *Scaptodrosophila lebanoni* is detected in both subclades in basal position as could be expected. Most of this gene tree topology may be the explained by an ancestral endogenization that would have occurred before the diversification of these species, followed by a single duplication event and losses in lineages that do not nowadays encode this gene. This host integration event was previously identified by Ballinger et al. [60]. Their detailed analysis suggested that this gene has been coopted in *Drosophila* genomes. Our analysis, based on a more recent database further extends the diversity of species concerned by this event as it now includes the basal species *Scaptodrosophila lebanoni*. Clearly, this result deserves further investigation in order to clarify the evolutionary history of this virally-derived gene and test its potential phenotypic effect on *Drosophila sp.*.

A virus related to Wuhan insect virus 15 was part of this module (here called Qinviridae_L.h). A single 5.5kb contig was found, where we expect a bisegmented genome (one 1601bp segment encoding an hypothetical protein and the other is 5889 bp long and encodes the RdRp). Nevertheless, it contains a full length RdRp protein from which we constructed a phylogeny (fig. S11) indicating that this virus probably belongs to the family Qinviridae [11].

We found a 1894bp contig with 93% identify with the Hermitage virus (here called Hermitage_L.h). Hermitage virus was first described in Webster et al. 2016. No phylogeny was built since our pipeline detected a single homologous sequence in the database.

A putative Kwi virus, here named Quenyavirus_L.h, composed of 11 contigs was also detected. Among them, 3 contigs have no homology with known sequence in public database but, it was considered that they belong to this genome because of their co-occurrence with the other contigs (see fig. S3) and that they do contain an ORF covering most of the contig. Kwi virus has been described by [61] by analyzing “dark matter” obtained from a previous study. Their genome is expected to be composed of 5 segments approximately 2kb each. We think that several variants are present in our dataset, as some of these contigs appear to be homologous (contig_8520 and contig_9238; contig_9023 and contig_13838; contig_10017 and contig_19523). Both variants appear to be present at Ige in June 2012, whereas only one was present at the same location in 2011; the other variant is the unique we found at Gotheron in 2012. The phylogeny based on the putative RdRp confirms the proximity with the founder members of this new clade (Kwi and Nai viruses), referred to as Quenyaviruses, with which they form a well supported monophyletic clade (fig. S12).

**Module 2: viruses mainly driven by *Asobara sp*.**

Five viruses were detected in *Asobara* species, including a putative reovirus composed of seven contigs (called Reoviridae4_A.sp, fig. S18) and a partitivirus composed of two contigs as expected for partitiviruses (called Partiti-like2_A.sp, fig. S24). A partial genome of nidovirus referred to as Nidoviridae_A.sp was also detected. This virus is related to Fuefuki-like virus which has been detected in *D. suzukii* [25]. Nidoviridae have non segmented 16kb +ssRNA genomes. In our case, the contig is only 1kb, suggesting it is incomplete. No phylogeny was constructed, as no homology other than that of the Fuefuki virus could be detected. We also found a contig (contig_13728) encoding a partial RdRp (only 424 amino acids whereas the closest relative has is 2172 amino acid long) related to Chuviridae (called Chuviridae2_A.sp, see phylogeny in fig. S25). Finally, two contigs (contig_10171 and 16255) were named "dark2_virus", since they were associated together in the *Asobara* sample (fig. S3), each encoding a large protein (see suppl. file S1), but without homology with public databases.

**Module 4: viruses mainly driven by *Trichopria sp*.**

Nine RNA viruses were mostly found in *Trichopria sp.*.

Chuviridae3_Tricho was identified as a mivirus-like composed of three contigs (5374, 12656, 11296) totalizing 5557bp. Mivirus belong to the Chuviridae family and have either one or two segments encoding typically L and G protein, a N protein and a VP. Contig 11296 encodes an homolog of a nucleoprotein (N), Contig 12656 encodes an homolog of a Glycoprotein (G), Contig 5374 (287-1540) encodes an homolog of a nucleoprotein (N) and Contig 5374 (1884-2501) encodes an “hypothetical protein”. We were not able to find the RdRp from these three contigs.

A 7315bp contig (dsRNA_virus1_Tricho) revealed sequence similarity (36%) with Circulifer tenelus virus. Circulifer tenelus virus is known as a non segmented dsRNA virus with a 8086bp genome [62]. The phylogenetic analysis conducted on the RdRp protein revealed that the virus found in *Trichopria sp.* formed a monophyletic clade with other non segmented RNA viruses (Circulifer tenelus virus, Spissistilus festinus virus 1 and Persimmon latent virus) whose genome sizes are respectively 8086bp, 7951bp and 7475bp (fig. S15). This suggests that the contig assigned to Trichopria circulifer virus is complete or almost complete. Circulifer tenelus virus and Spissistilus festinus virus were isolated from threecornered alfalfa hopper (*Spissistilus festinus*) and beet leafhopper (*Circulifer tenellus*), two plant-feeding hemipteran insect pests. The taxonomic assignment of these viruses is unclear but they show proximity with Chrysoviridae and Totiviridae.

Ten contigs were considered as composing a new Reovirus genome totalizing 23900bp (named Reoviridae2 Tricho). Among them, seven showed clear signs of homology with reoviruses (30% protein identity) and four contigs without homology in our initial analysis did show a perfect association with them (see fig. S3). In a subsequent analysis using an updated nr databases, three of them revealed homology with reoviruses (blastx): contig_9440 shows very weak similarity with Zoersel tick virus (VP7, QYV43125.1, evalue 3e-7, 24% identity), contig_23185 shows similarity with VP5 from Zoersel tick virus (QYV43123.1, evalue 2e-24, 23 %identity). Contig_7972 does not display any similarity with known sequences but contains a single ORF covering the majority of the contig, as other segments do. Zoersel tock virus is an unclassified Reoviridae. contig_13079 has no sequence similarity with any public sequence. A phylogeny built on the RdRp revealed its proximity with Operophtera brumata reovirus and Eccles virus (Fig. S18).

We found 3 contigs (ranging in size from 1737bp to 2143bp) related to Wuhan insect virus 22 (48% identity on RdRp). Wuhan insect virus 22 has been classified in the "Partiti-Picobirna" clade by [12]. It is composed of two segments encoding respectively the RdRp (1869bp), and an hypothetical protein (1766bp). In our dataset, it is unclear whether the three contigs belong to the same virus or if only one of the two contigs encoding the “hypothetical protein” does. We built a phylogeny based on RdRP (fig. S24) that confirmed the proximity with Wuhan insect virus 22, Galbut virus and other Partitiviridae such as Vera virus. We refer to this virus as Partiti-like4_Tricho.

Six contigs were considered to be part of a single incomplete Phasmaviridae related to Ganda bee virus (Phasmaviridae_Tricho). Four of the contigs encode parts of an RdRp suggesting either that several viruses are present or that the assembly is fragmented. After inspection of the blast results, it was clear that the assembly is incomplete leading to a fragmented Ganda-like RdRp. We artificially fused the 4 parts of the RdRp (order is: contig_10992, contig_10108, contig_10707 and contig_17877) which covered most of the the related RdRp protein (from positions 57 to 1958, out of 2087 amino acids for YP_009666981.1) in order to build a phylogenetic tree (fig. S16). This tree confirmed the proximity with Ganda bee virus and other related Phasmaviridae. Ganda bee virus genome is typically composed of three segments: 6453bp for the RdRp coding segment, 2101bp for the glycoprotein precursor (GnGc) gene and 1906bp for the nucleoprotein (N) gene (see fig 3 in [63]). It seems that the genome is almost complete, apart from the fact that the RdRp is scattered among 4 contigs: contig 9260 putatively encodes the Glycoprotein (M segment), Contig 6541 encodes the nucleoprotein and other unannotated ORFs (S segment), and the other four contigs collectively encode the RdRp protein.

We found 5 contigs presumably belonging to a single virus (Rhabdoviridae1_Tricho) related to Hubei Dimarhabdovirus 2. Hubei Dimarhabdovirus 2 is a non segmented virus composed of a 11332bp genome. Our assembly is thus fragmented since we obtained 5 contigs. However it seems that they cover at least most of the 2119 amino acids that compose the RdRp (YP_009337071.1). Contig 11939 encodes a protein that aligns with YP_009337071.1 from 11 to 497 (contains a RdRp domain); Contig 10949 encodes a protein that aligns with YP_009337071.1 from 531 to 1035 (contains a RdRp domain); contig_3788_54_3266_+ encodes a protein that aligns with YP_009337071.1 from 1048 to 2119 (contains a mRNA capping region and a viral-capping methyltransferase). The phylogenetic reconstruction based on RdRP confirms its assignation to Rhabdoviridae (fig. S20).

22 contigs were assigned to a possible reovirus in *Trichopria sp.* that we refer to as Reoviridae5_Tricho. Among these contigs, nine had homology with reoviruses, in particular with Rice gall dwarf virus (RGDV), which has a 12-segmented dsRNA genome. The remaining 15 contigs had no homology with known sequences but are likely part of this reovirus genome based on their association within samples (fig. S3). Three reovirus-like contigs spanned the major part of the RdRp from RGDV (YP_001111373.1, 1458aa long): contig_12726 covers amino acids from position 50 to 433, contig_18151 covers from position 407 to 781 and contig_8245 covers amino acids from position 870 to 1386. We merged these three parts of the RdRp to build a phylogeny which further suggests this is a reovirus related to Rice Gall Dwarf virus (fig. S18).

A chuvirus, composed of 4 contigs, was identified (Chuviridae4_Tricho). The majority of the RdRp of the closest relative (Hubei chuvirus-like virus 1, 2172 aa long, YP_009337904.1) is covered, but is split between two contigs: contig_22765 covers protein from 201 to 1698 and contig_13828_62_1273_-covers from 1791 to 2158. Hubei chuvirus-like virus 1 is composed of two segments 6873bp and 3958bp. The P-protein seems to be lacking in our dataset, indicating that this is a partial genome. However, we were able to build a phylogeny based on RdRP domain (fig.S25) that confirmed the positioning of this virus within the Chuviridae family.

We found a 10kb contig showing homology to Virga-like viruses in *Trichopria sp.* (24% identity, 10210bp). The assembly is likely complete or almost complete since Virga-like viruses are non segmented +ssRNA viruses, with genomes up to 10kb [64]. The virus, referred to as Virga_Tricho, is composed of a single Contig_923_1627_10098 encoding the expected domains (as in fig. 2 of [64]). The phylogeny given in fig. S19 confirms the proximity with virga-like viruses such as Hubei virga-like virus 1 and 2.

**Module 1: viruses mainly driven by *Pachycrepoideus sp*.**

Four viruses were strictly associated with *Pachycrepoideus sp.*, including a DNA virus discussed above (a Parvoviridae) and three RNA viruses.

We found a contig showing similarity (33%) with glycoprotein G of Cumuto Goukovirus. Goukoviruses are negative sense ssRNA viruses belonging to the Phenuiviridae. They infect insects and have a genome composed of 3 segments (1.1kb, 6.4kb and 3.2kb). In our case, the protein is most likely incomplete (327 aa versus 1000 aa for the related sequences). The phylogeny confirms the proximity with Cumuto Goukovirus (fig. S23).

A set of three other contigs were found to be strictly associated in one of our *Pachycrepoideus sp.* samples. These three contigs showed homology with Black queen cell virus, which belongs to the family Dicistroviridae and is composed of a single Monopartite, linear ssRNA(+) genome of around 9 kb. In our case, the three contigs sum up to 8114bp suggesting it is almost complete, although fragmented. The phylogeny built on the RdRp confirmed the positioning of this virus in the Dicistroviridae family, in a clade containing Drosophila C viruses (fig. S22).

Three contigs were found to be strictly associated in one of our *Pachycrepoideus sp.* samples (here called Partiti-like3_Pachy). They showed sequence similarity with viruses described in [12] (Wuhan cricket virus 2 and Hubei tetragnatha maxillosa virus 8). These viruses are related to the partitiviridae thus we call them "partiti-like". In addition, one of the contig (contig_15227) encoding a large 392aa ORF did not show sequence homology with known sequences, but did show a perfect association with the other partiti-like contigs. Since partiti-like viruses are composed of 4 to 6 segments, it is likely that this contig correspond to an additional segment. The phylogeny built on RdRp shows that this virus together with Wuhan cricket virus 2, Wuhan millipede virus 4 and Wuhan tetragnatha maxillosa virus 8 do form a monophyletic clade, related to Partitiviridae (fig. S24).

###### Viruses not included in modules

Four putative viruses, three of them being RNA viruses, were not included in any module since they had a more diffuse distribution among insects.

A 1044bp contig (contig_20830) without homology with public databases in our initial analysis, was considered as a viral candidate (here denominated Dark1). This was motivated by the fact it encodes a large ORF (suppl. file S1). This interpretation was validated by a less stringent analysis revealing a low sequence similarity with an hypothetical protein from a *Diaphorina citri* cimodo-like virus (QXG83187.1 Length: 636 evalue 0.006). Most of the reads that mapped on this contig were generated from *C. amomyza* and *Leptopilina heterotoma* samples (see fig. 6), but other *Drosophila* and parasitoids also provided some reads mapping on this contig. Further studies are clearly needed to confirm the viral nature of this contig.

Three contigs were assigned to a Reoviridae (named Reoviridae_6) distantly related to other Reoviridae members with only 25% identity (fig. S18). The RdRp protein is split in two contigs (contig 6823 for the N-terminal part and 8100 for the C-terminal). Samples considered as positives were found in *L. heterotoma* and *D. kuntzei*, although the majority of reads were detected in *L. heterotoma* samples (see fig. 6).

We found a new Reoviridae related to Bloomfield virus (26-30% identity) which was first detected in *D. melanogaster* [54]. This putative virus, here denominated Reoviridae7, was composed of 5 contigs. The phylogeny built on RdRp confirmed the proximity with Bloomfield and Grange viruses (fig. S18) which were detected in *D. subobscura* [55]. These three viruses formed a highly supported monophyletic clade. This virus was detected in *Leptopilina sp*, *Pachycrepoideus sp*. and *D. kuntzei* (fig. S4), although the majority of reads were detected in *L. heterotoma* samples (see fig. 6).

### 3.7 Case study of two viruses detected in the parasitoid *L. heterotoma*

Six *L. heterotoma* isofemale lines originating from a different location (Lyon, Fance) were analyzed by rt-PCR for the presence of two viruses discovered during this study, namely Phasmaviridae_L.h and Iflaviridae_L.h. All six lines were positive for Phasmaviridae_L.h, while only 4 were positive for Iflaviridae_L.h. The presence of both infected and uninfected lines for the Iflaviridae_L.h, offered the opportunity to determine its mode of transmission and overall phenotypic effect. As expected, in crosses involving both infected females and males (line 3 x line 3), all four offspring tested were positive for infection confirming vertical transmission of the virus, while no infection was observed in the cross involving uninfected individuals (line 38 x line 38, fig. S26). When infected females were crossed with uninfected males (3x38), all offspring were also infected; while in the reciprocal cross (38x3), no infection was observed (fig. S26). From these results, we concluded that Iflaviridae_L.h was strictly maternally transmitted along generations. Additionally, this experiment did not reveal any significant differences in the three phenotypic traits measured between the four modalities (parasitoid induced mortality (PIM), pre-imaginal developmental success (PS), sexratio, Kruskal-Wallis test, df=3, all p-values >0.05, fig. S27).

This experiment revealed an unexpected result in the “no RT” control. Such a control contains all the reaction components except for the reverse transcriptase. It is used to test for contaminating DNA (such as genomic DNA in the case of a transcriptomic study). Reverse transcription should not occur in this control, so if PCR amplification is seen, it is most likely derived from DNA. In our assay, as expected, no amplification was observed in the "no RT" controls from individuals that do not show signs of infection (no amplification in the reactions with RT, fig. S26-B). However and unexpectedly, an amplification was observed in the "no RT" controls performed on individuals that show signs of infection (positive in the reaction with RT, fig. S26-B). The presence of amplification in the no-RT controls could, in principle, be the consequence of an integration of the viral locus in the wasp genome. However, if this was the case, amplification should also be observed in the offspring of infected individuals (since they should inherit at least one copy of the endogenized locus). However, this was not the case: the profile obtained for individual 13, offspring of an uninfected female crossed with an infected male, was negative with or without RT (fig. S26-B). This ruled out the possibility of a genomic integration of the viral locus targeted by our rt-PCR assay. Hence, from this experiment, we conclude that infected individuals do produce or inherit from their infected mother a viral DNA copy of the viral RNA (vDNA). Similar cases of maternal transmission of vDNA have been described in the fly *D. melanogaster* and the tiger mosquito *Aedes aegypty* challenged with several positive-sense single stranded RNA viruses [65].

We then tested whether the virus could also benefit from horizontal transmission when infected and uninfected wasps shared the same *Drosophila* host, as observed in the behaviour-manipulating virus LbFV infecting the related wasp *L. boulardi* [20]. To test this possibility, *Drosophila* larvae were sequentially offered to unfertilized infected wasps and then to fertilized uninfected wasps. Because of the haplo-diploid sex determination in Hymenoptera, this setup ensured that emerging females were offspring of the uninfected female. In case of horizontal transmission, we expected at least part of these emerging females to be positive for infection. In this assay, four emerging females were rt-PCR tested individually, as well as a pooled sample of 20 emerging females to increase power in the event of low horizontal transmission rate. A control test indicated that our rt-PCR assay was capable of detecting a single infected female in a group of 20 (fig. S26-C). None of the four emerging females tested individually was positive while the group of 20 emerging females was positive (fig. S26-C). These results show that, although quite infrequent, horizontal transmission of the virus occurs when infected and uninfected females share the same hosts (between 1/20=0.05 and 1/4=0.25). Note that because we did not directly measure the occurrence of superparasitism, the frequency of HT reported here must be seen as a lower bound value, because uninfected wasps that by chance developed alone were not exposed to the virus.

Finally, since some horizontal transfer through host sharing (most likely occurring when *Drosophila* larvae are super-parasitized by both infected and uninfected females), we tested whether the Iflaviridae_L.h could manipulate the superparasitism behaviour of the wasp, as observed for the virus LbFV infecting *L. boulardi*. We measured the superparasitism intensity under two densities: either one female or a group of 3 females was placed on a batch of ten *Drosophila* larvae. No superparasitism was observed for any larvae when a single female laid eggs, whatever their genotype or infection status. Moderate superparasitism was observed when three females were foraging, but this was independant of Iflaviridae_L.h infection (fig. S28, Kruskal-Wallis test=3.668, df=3, p=0.3). This experiment thus did not provide evidence of behaviour alteration induced by the Iflaviridae_L.h.

## 4 Discussion

Using a protocol designed to identify both RNA and DNA viruses, our analysis uncovered a rich community of heritable viruses in this interacting community of insects. As most studies generally focus on (or enrich for) viruses of one of the two types (RNA or DNA), neglecting the other, we believe this dataset is quite unique, giving an unbiased view of the virome in this insect community. Of the 53 viruses detected, the vast majority (45) had a genome composed of RNA. This result is in line with the diversity deposited in the public databases [16, 66], reinforcing the conventional wisdom that RNA viruses dominate the eukaryotic virus community [66].

Current knowledge about what factors shape viral communities in animals is still very limited, despite its importance in the context of sanitary vigilance required by certain systems. For instance, it is only very recently that factors such as season or host age structure shaping the viruses infecting the common vampire bat (*Desmodus rotundus*) or three common European rodent species have been identified [27, 67], despite the possible zoonotic risks associated with these species. In addition, these studies focused on single or very few species (3 in this case), reducing the power for detecting structuring factors. Here, we tested three main factors that were suspected to possibly shape the viral communities in this multi-host-multi-parasitoid system. The factors tested were namely geographic location, timing (season/year) and host species. Among them, the species factor was by far the most structuring one, explaining 52.5% of the total variance, while the other factors were not significantly associated with viral structure. Since we are focusing on vertically transmitted viruses (because we maintained the iso-female lines during two generations in the lab before viral purification), this suggests that vertical transmission necessitates specific adaptations in the virus genomes that comes with related costs in other species (trade-offs). This phenomenon then translates into specificity. We also tested whether insect phylogeny explains virome composition. This was not the case. This contrasts with results obtained on bacterial communities, where host phylogeny is the main driver of bacterial communities[6, 7]. However, we must stress that our sampling scheme here was limited compared to the datasets used to identify such an effect in insect bacterial communities, which may have reduced our power to detect such an effect.

In addition, we found that parasitoids had more viruses than their hosts. Because in this system, all parasitoids belong to the Hymenoptera order while all hosts belong to Diptera (which diverged more than 300 Mya), it is impossible to disentangle the effects of phylogeny and that of the lifestyle (free-living versus parasitoid). Additionally, for practical reasons, we had to use different temperatures for rearing the hosts (21°C) and the parasitoids (25°C). Thus we cannot rule out the possibility that the effect observed is driven by temperature, although to date, the available results obtained on a few viruses of the Drosophila/parasitoid community indicates either no effect of temperature on viral transmission ([31]) or an effect going in the opposite direction ([68]). Keeping in mind these limitations, we can discuss the reasons that may lead to such a pattern. As mentioned in the introduction, parasitoids have established a special relationship with viruses. This can be seen from free-living infecting viruses, as well as through the abundance of endogenized version of viral genes (in particular deriving from dsDNA viruses) specifically in the genomes of parasitoid wasps (but not free-living hymenoptera, [16]). Thus, this result further suggests that the reproductive tract of parasitic wasps is an ideal location for viruses, allowing them to efficiently transfer along generations either vertically or horizontally in case of superparasitism. Accordingly, our experimental assay on the Iflaviridae discovered in *L. heterotoma* revealed that the vertical transmission of the virus was achieved thanks to maternal transmission (but not paternal transmission), and that low but detectable rate of horizontal transmission (around 1/20=5%) was at play in this system. Both transmission features are expected if the virus is injected together with the egg into the host (typically produced in the venom gland as observed for other viruses[20][35]). Confinement within the host’s body certainly favours transmission to developing offspring. Drosophilidae, on the other hand, lay their eggs in a much less confined environment (typically rotting fruit), possibly explaining the difference. Although we can not rule out that phylogenetic inertia or rearing temperature explain the pattern, this difference in heritable virus abundance between parasitoids and their hosts may well be explained by the particular biology of parasitoids.

Another beneficial effect of studying the viral community as a whole is that it gives us a better understanding of host specificity. For instance, before our work, a large scale work focusing on *Drosophila melanogaster* identified a new Filamentous virus (called DmFV for D. melanogaster Filamentous Virus) as a potential DNA virus of *Drosophila melanogaster* [48]. This work relied on pool-seq data obtained on wild-collected individuals. This virus was discovered in one pool of 30 individuals (originating from Spain) out of 167 pools collected throughout Europe. On the contrary, and despite all specimens collected here were sympatric, we found similar sequences in the parasitoid *L. heterotoma*, but none in the *Drosophila* hosts. This led us to suggest that this virus is in fact a parasitoid virus, that may be occasionally found in *Drosophila*, either because of a temporary spillover, or more likely, because *Drosophila* from the field may have been attacked by parasitoids during their larval life, but survived thanks to an encapsulation response (this scenario was also envisaged by Wallace and colleagues although they favoured the Drosophila virus hypothesis). In total, we screened 230 isofemale lines of *D. melanogaster* and none appeared to be positive for this virus, while two of our five samples of *L. heterotoma* were positive (totalizing 27 isofemale lines out of 108 in total). This suggests that the two generations of vertical transmission purged the possible presence of similar DmFV traces in field-collected flies, while allowing the transmission of the virus in the parasitoid *L. heterotoma*. This reinterpretation better matches the virus’ phylogenetic position, since its closest relative infects the related wasp *Leptopilina boulardi* [42]. We thus propose to call it LhFV for L. heterotoma Filamentous Virus instead of DmFV. This virus is a member of a new viral family that will be described elsewhere[34].

Finally, this global survey of Drosophila and their parasitoids, identified only one virus (or possibly two depending on the metrics used) in *D. suzukii*. This species, native to Southeast Asia, has been invasive in North America and Europe since 2008, and is associated with dramatic losses in fruit production due to the unusual behavior of females, who lay their eggs in unripe fruit using their sclerotized ovipositor. The factors underlying the invasive success of this species have been searched for by several means [69]. Here, our data show that *D. suzukii* is among the species with the lowest number of heritable viruses. This may originates from a lower overall abundance of heritable viruses in this species in general (including in native populations) or could be a specific feature of introduced populations. If this last interpretation is correct, and if heritable viruses are costly to *D. suzukii*, then this may suggest a role for this reduction in viral prevalence in invasive success. This idea, known as enemy release hypothesis [70], needs to be tested in *D. suzukii*.

In conclusion, our exploratory analysis has revealed a rich community of heritable viruses in these interacting insect communities. Because we purified viruses after two generations of lab-rearing, we probably enriched not only for vertically transmitted viruses, but also for viruses having low virulence (otherwise the insects may not have been collected in the wild, and selection may have eliminated them during the lab-rearing). The combination of vertical transmission, low virulence, and strong host structuring makes these viral partners potential sources of heritable variance in phenotypes [71], with the possibility of beneficial effects, similarly to what is observed for symbiotic bacteria. Measuring their overall contribution to phenotypic variance will require phenotypic assays such as the one we carried out on the Iflaviridae_Lh/*L. heterotoma* system, as well as estimates of viral prevalence in different environments.

## Acknowledgements

We are indebted to Ms Weistroffer (Igé) and the INRAE station of Gotheron for allowing us to collect the insects in their garden/orchards. The bioinformatic work was performed using the computing facilities of the CC LBBE/PRABI. The sequencing has been performed on the ProfileExpert platform of the University Lyon 1. We thank Stéphane Dray for helpful discussions on the variance partitioning analysis, Darren Obbard for its input on the Vesantovirus sequences and three anonymous PCI reviewers for helpful comments. This work is dedicated to Roland Allemand, who left us too soon.

## Supplementary information availability

Scripts, codes and additional figures are available online: https://github.com/jVaraldi/Viromics/ or https://zenodo.org/badge/latestdoi/672215335 (permanent doi).

## Annexes or Supplementary Information

Supplementary file S1: S1_All_orfs_predictions.pdf. This file contains all the ORFs predictions for the 53 putative viruses.

Supplementary file S2: S2_CO1_assignment.pdf. This file shows the mapping of CO1 reads on the BOLD database for each sample.

---

**Table S1.**
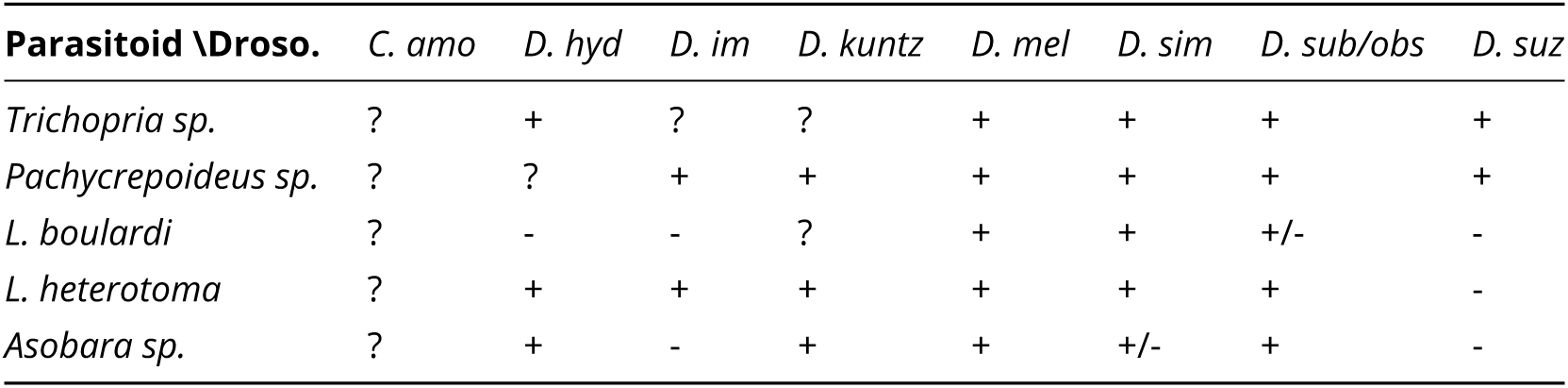
Host range of the five major parasitoid species attacking Drosophilidae. Based on [29, 72–75] and personal observations.

**Table S2.**
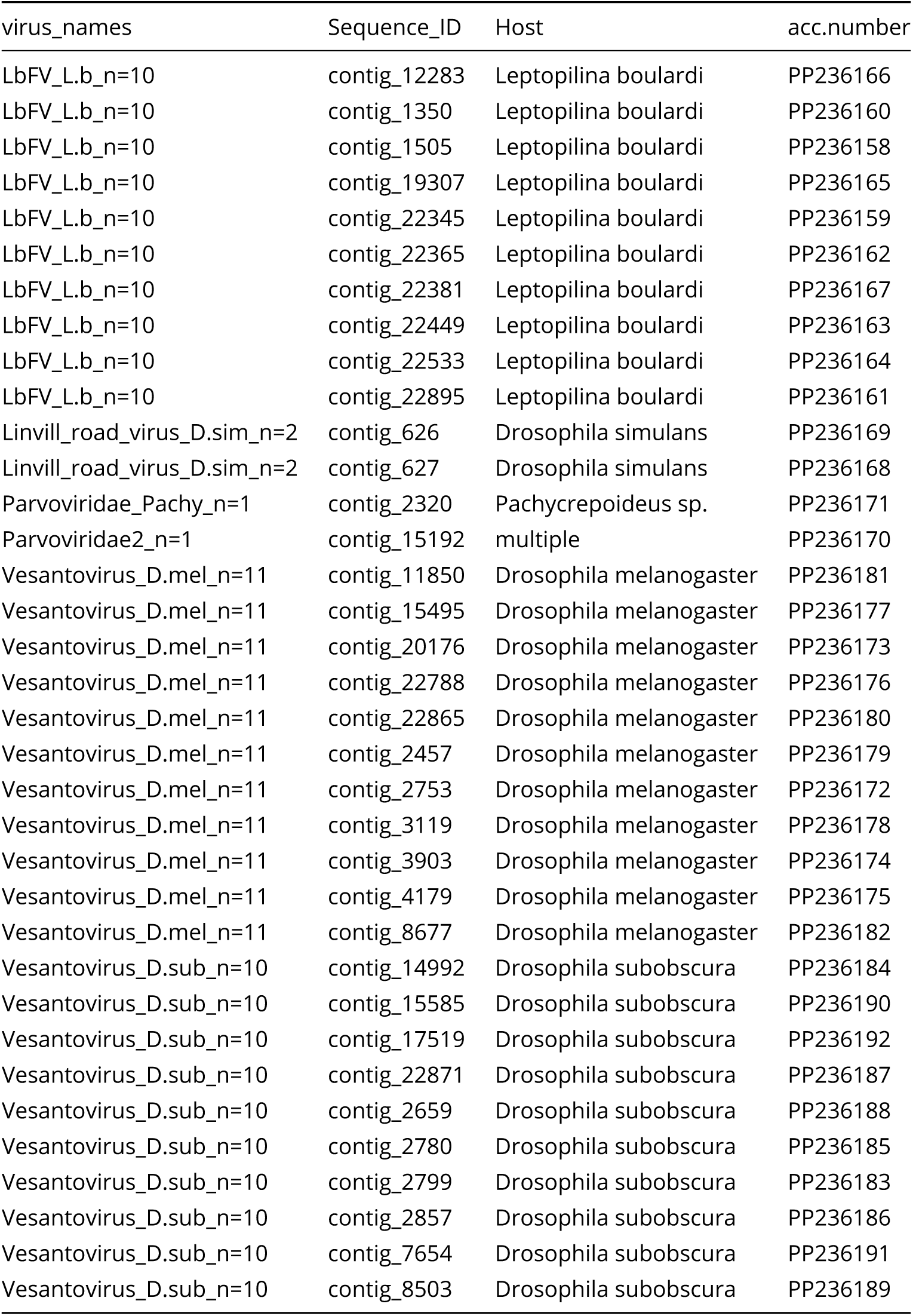
Informations related to DNA virus contigs

**Table S3.**
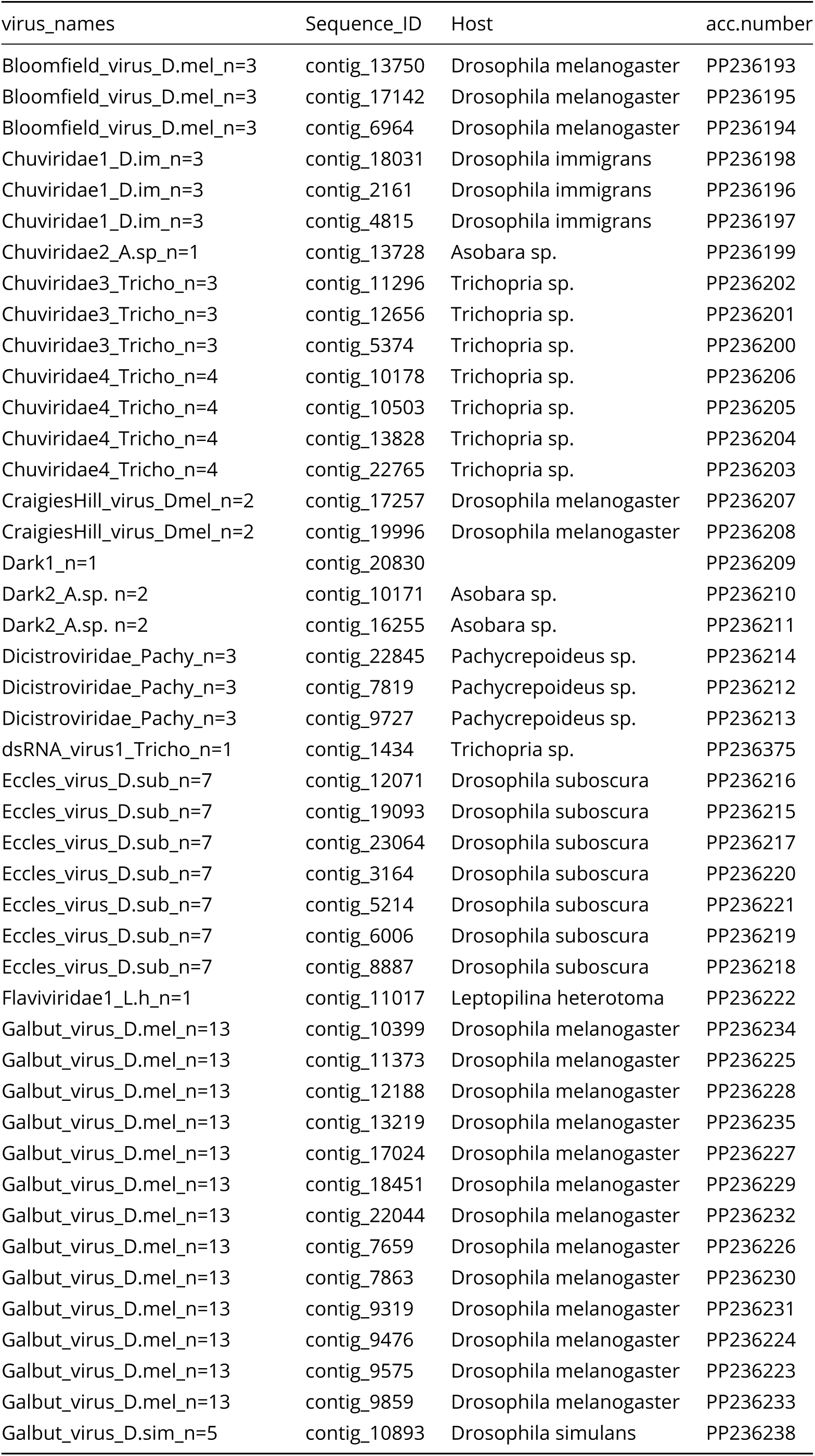
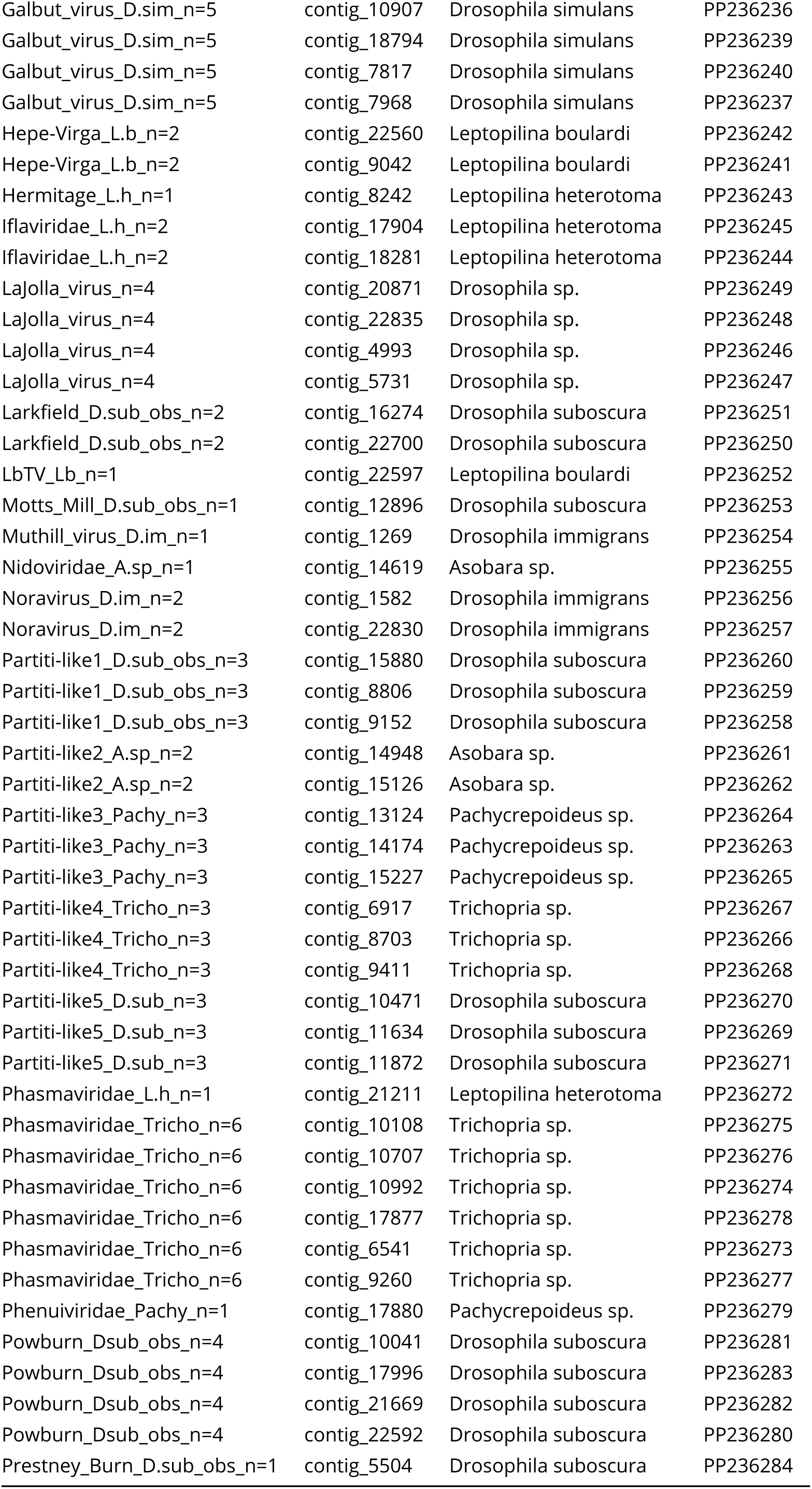
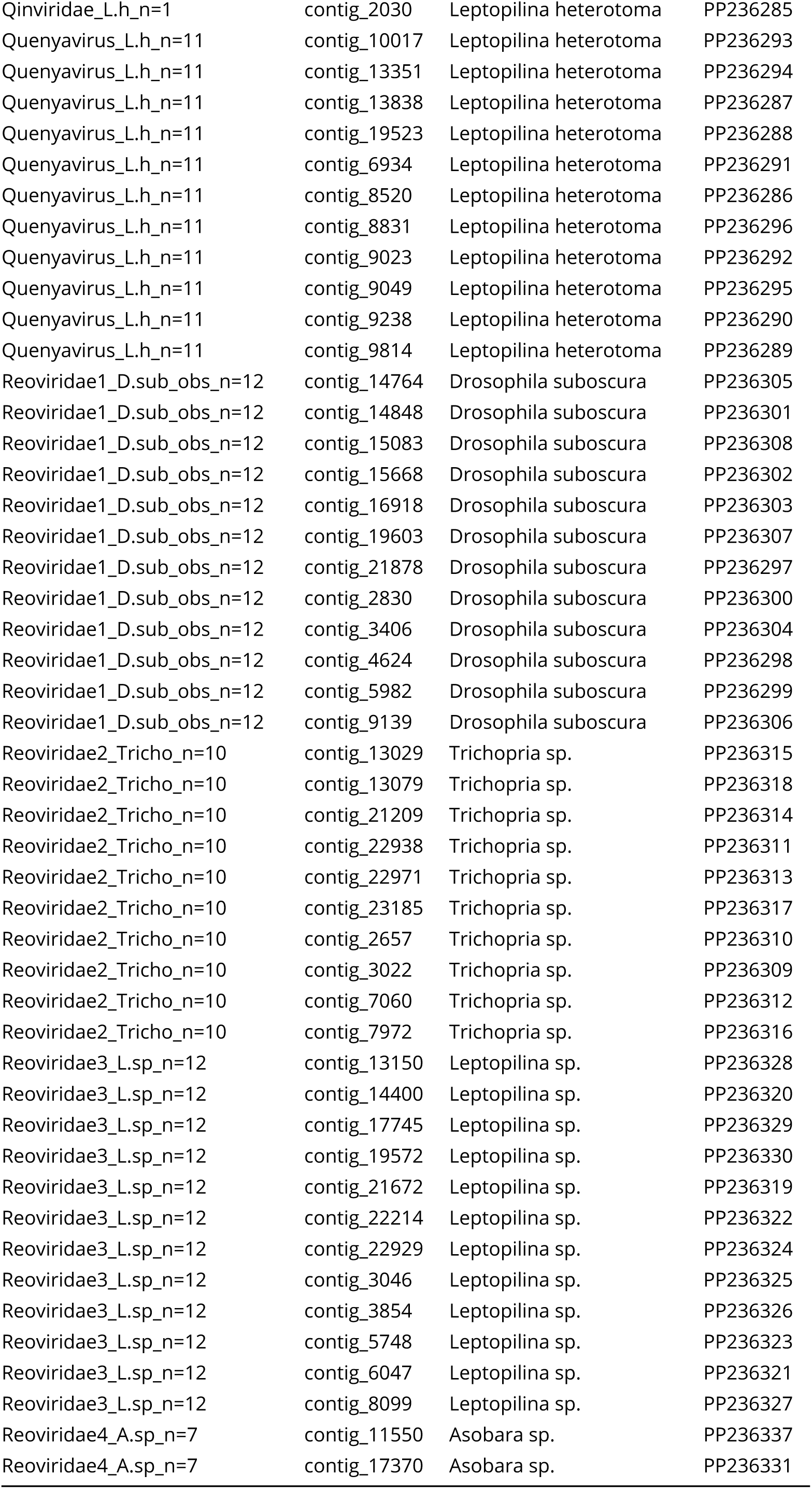
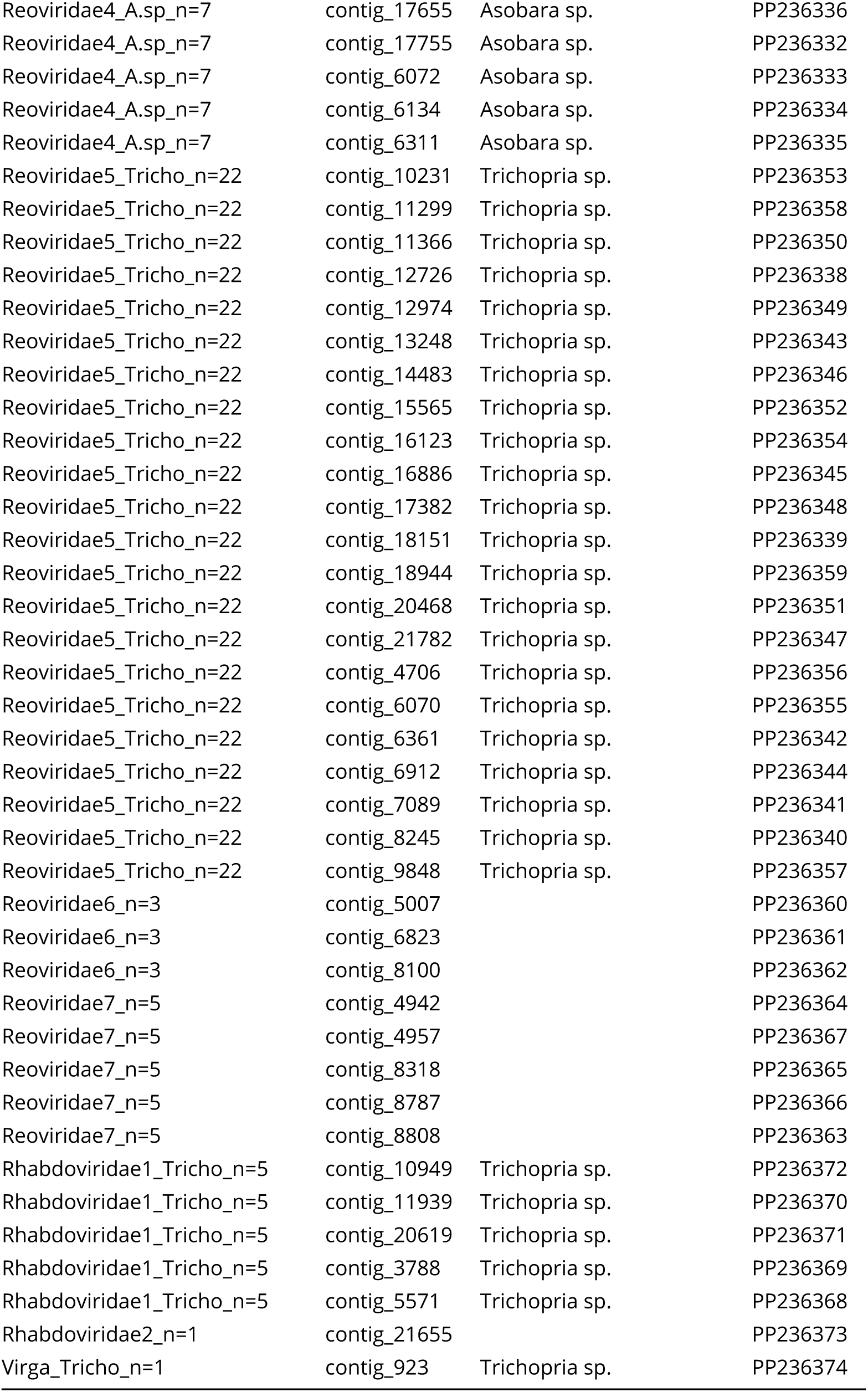
Informations related to DNA virus contigs

**Figure S1.**
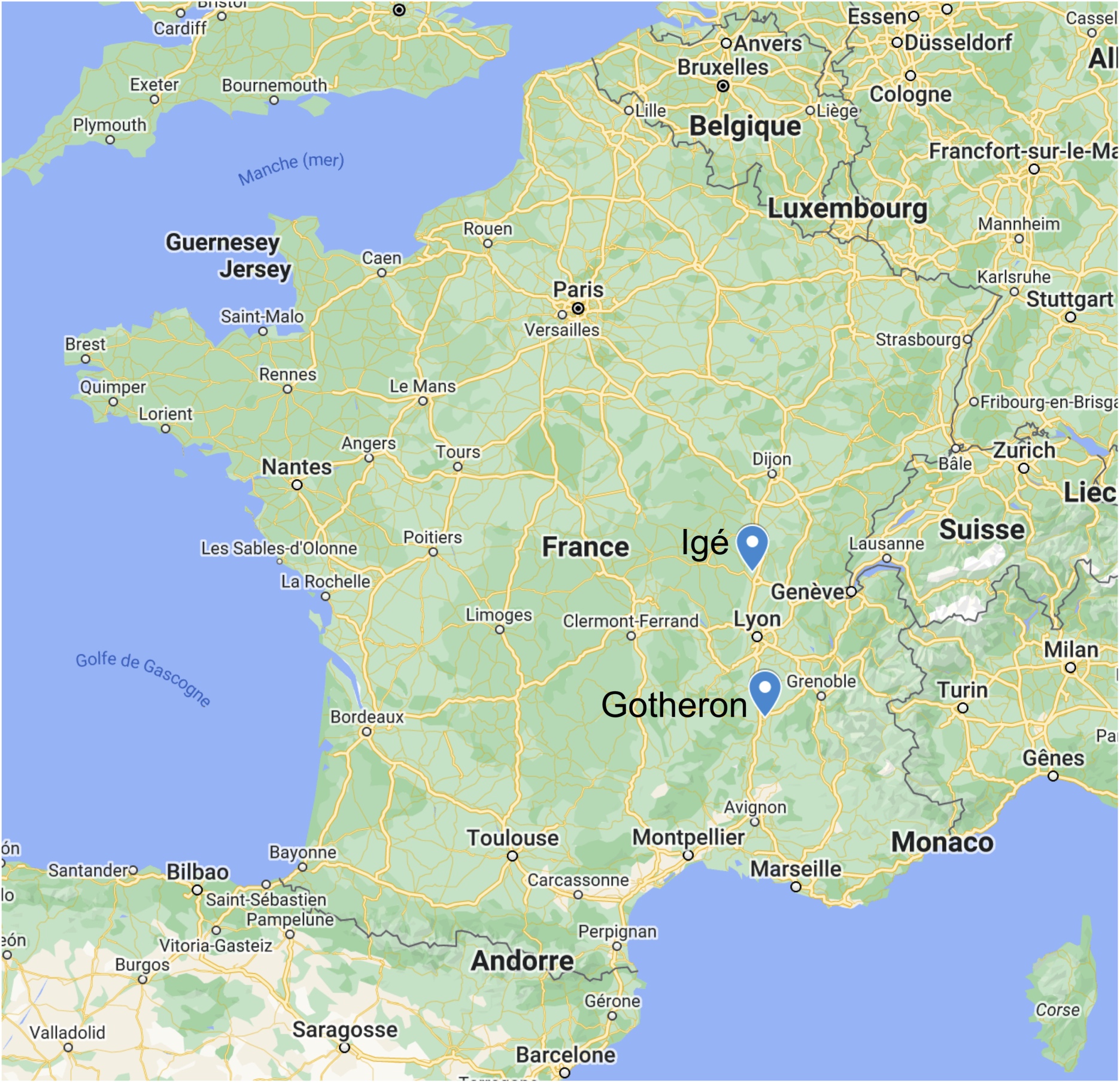
Sampling sites. Igé, Burgundy; GPS: 46.399N, 4.742E; Gotheron, near Valence, GPS: 44.978N, 4.930E.

**Figure S2.**
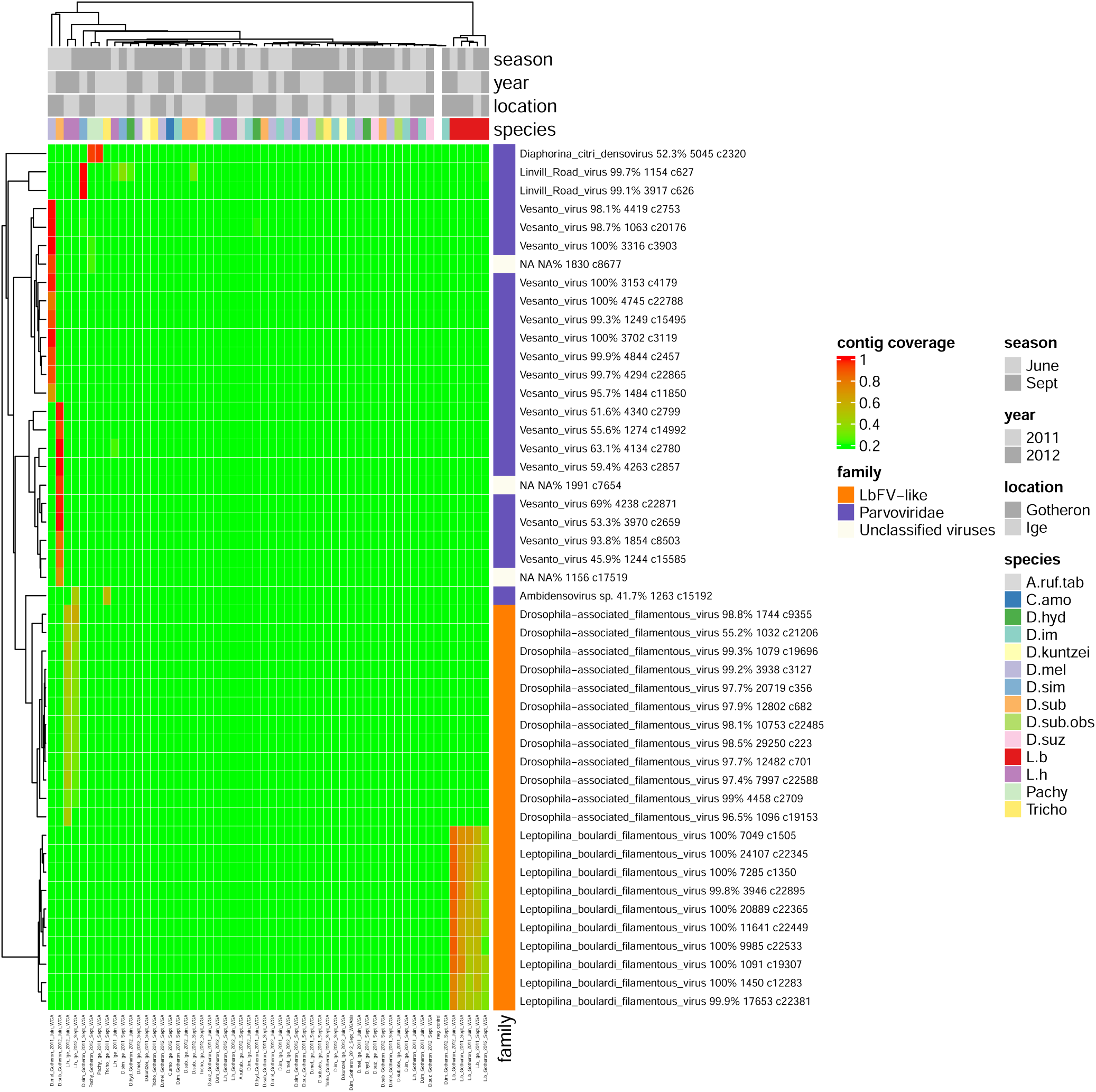
DNA virus contigs found in the Drosophila/parasitoid community

**Figure S3.**
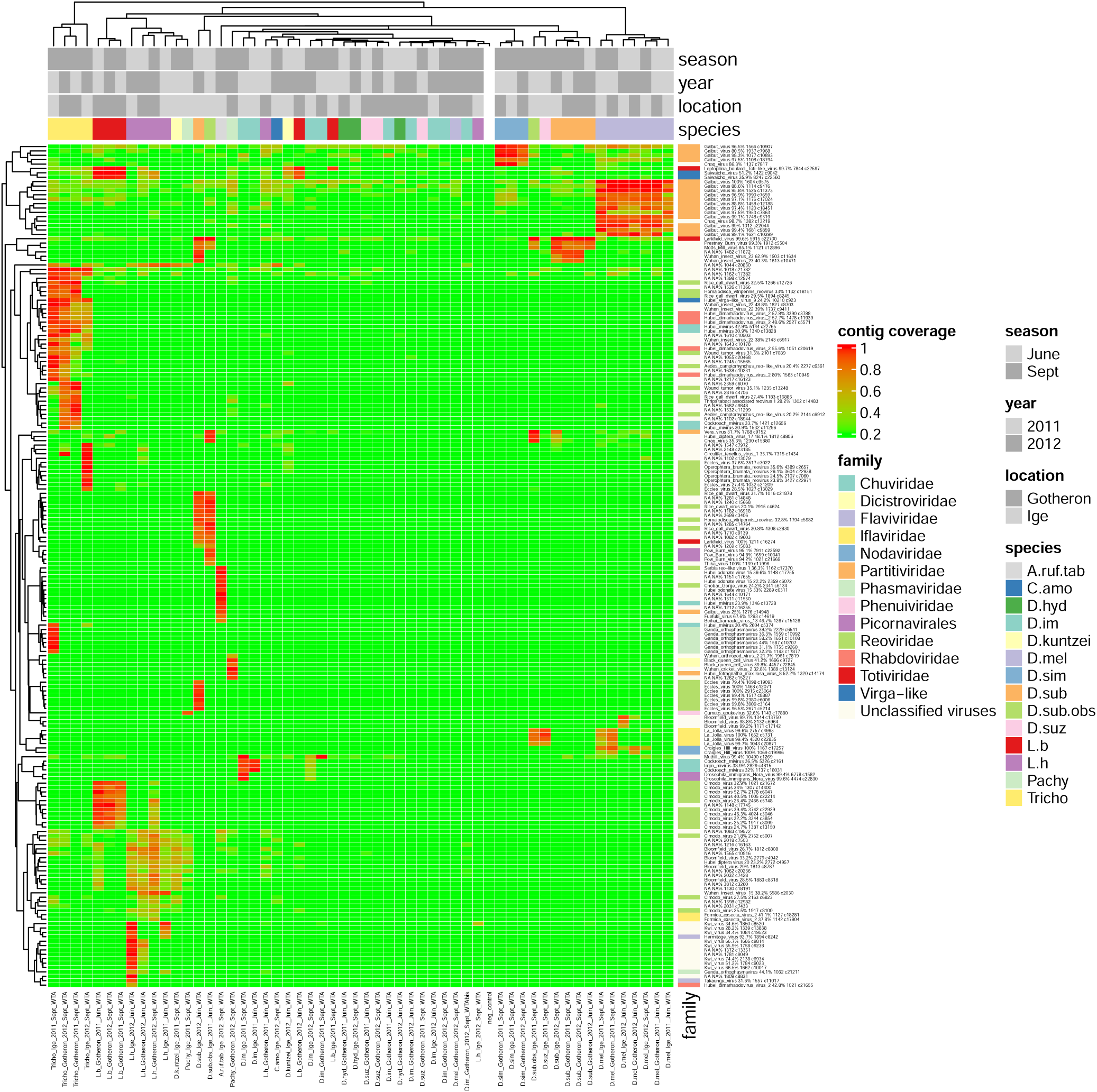
RNA virus contigs found in the Drosophila/parasitoid community

**Figure S4.**
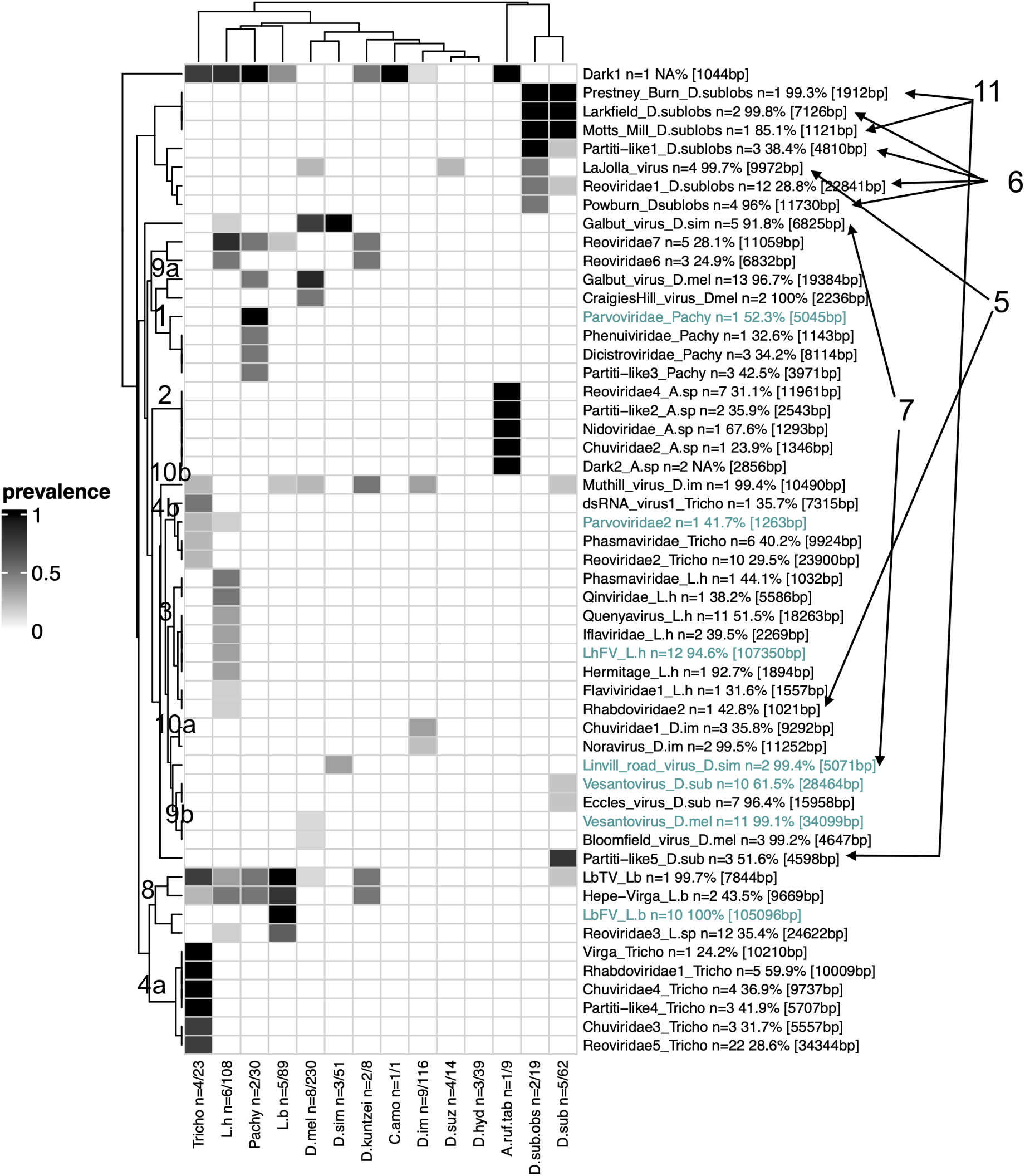
Virus distribution in the *Drosophila*/parasitoid community, grouped by host species. The "prevalence" is defined as the number of samples considered as positives (with a threshold for contig(s) coverage of 30%). The number of pooled samples (x1) and of isofemale lines (x2) analyzed is indicated next to each insect species names (n=x1/x2). Next to each virus name are indicated the number of contigs assigned to the virus, the percentage of identity with the first hit (amino-acids) and total length of the contigs in square brackets. RNA viruses are indicated in black, whereas DNA viruses are in green. Clustering method was performed using the Heatmap function default parameters (method "complete" based on euclidian distances, R package ComplexHeatmap) on both rows (viruses) and columns (insect species). The numbers displayed along the rows-clustering and in the right margin indicate the "modules" as defined in fig. 6 and discussed in the main text. Modules that are here split in two are noted a a and b. A few discrepancies were still remaining (i.e. Parvoviridae_2 which is not included in module 4, and Rhabdoviridae_2 which is not part of module 3 but 5.)

**Figure S5.**
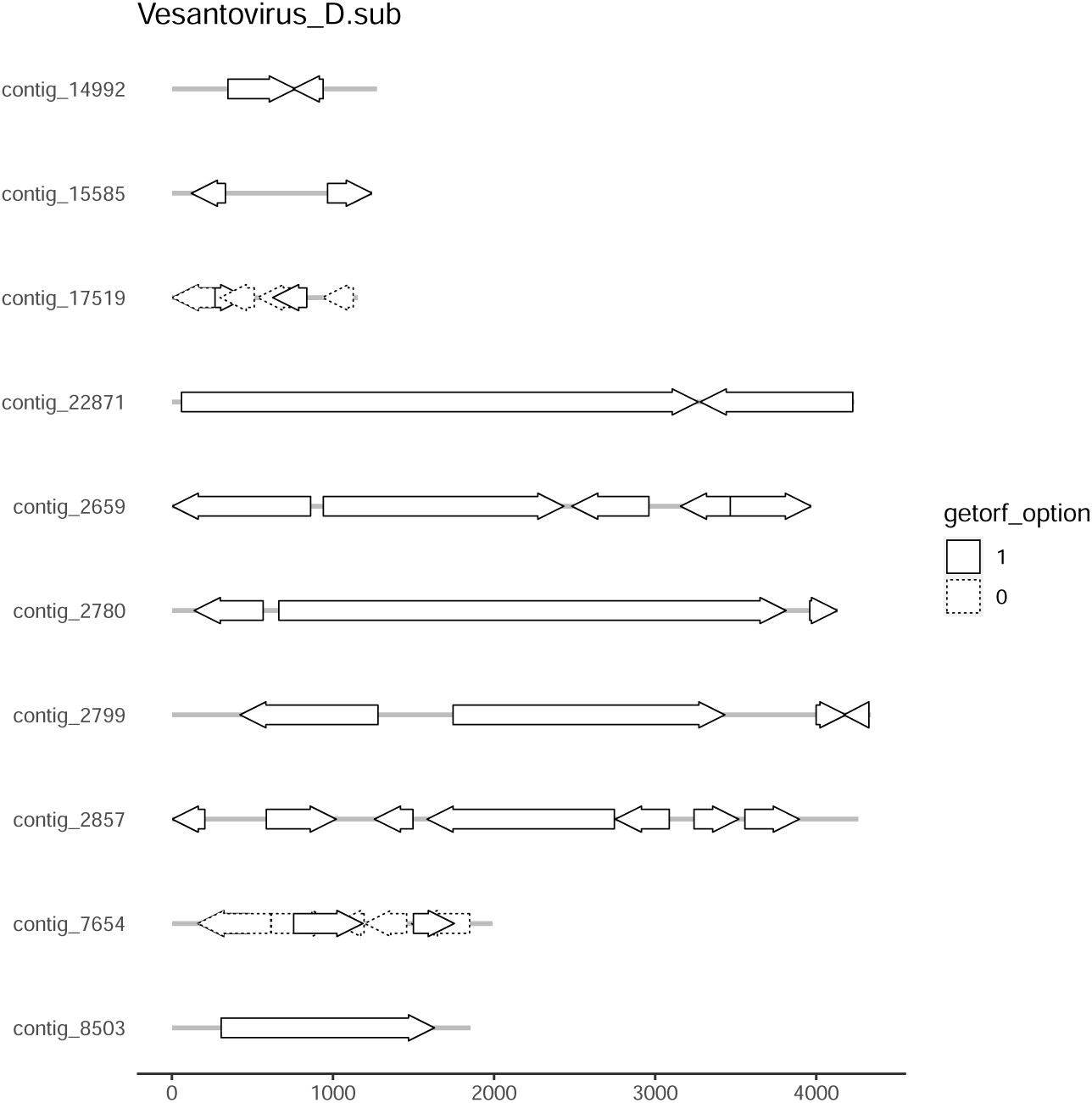
Annotation of a new Vesanto virus found in *D. subobscura*

**Figure S6.**
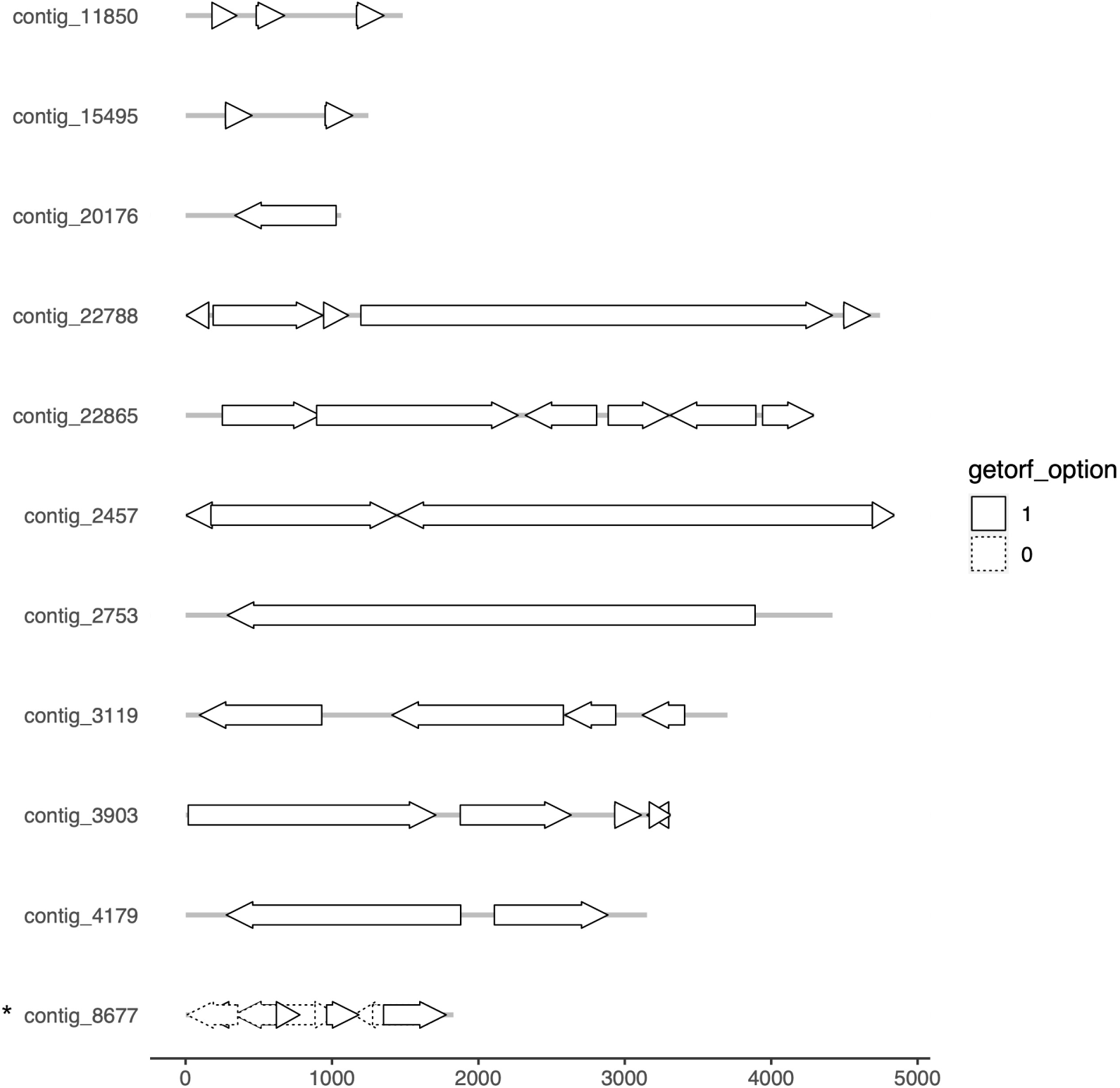
Annotation of the Vesanto virus found in *D. melanogaster*. The contig noted with a star is an additional candidate segment compared to what was reported in [48].

**Figure S7.**
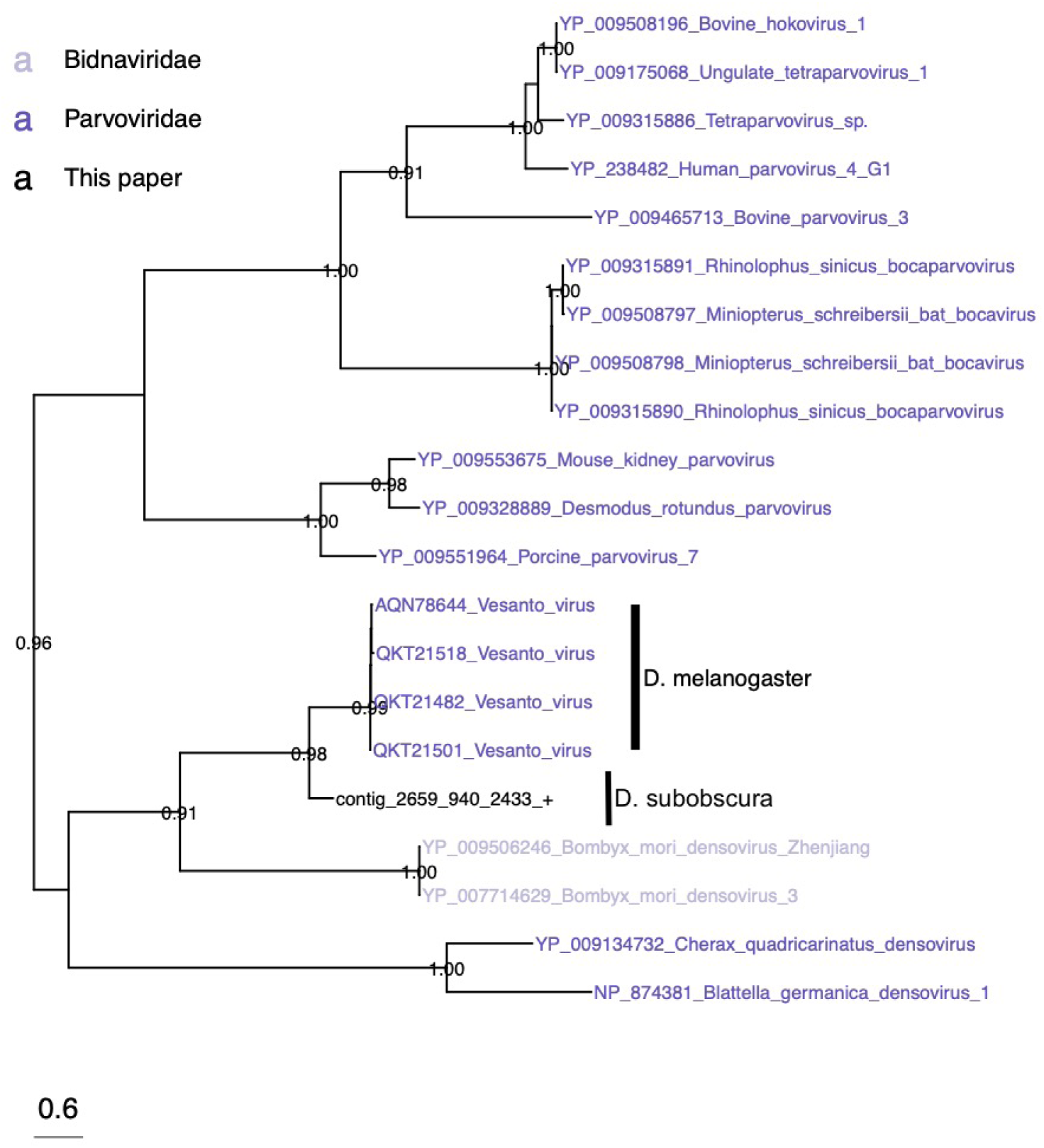
Phylogeny of the new Vesanto virus found in *Drosophila subobscura* based on NS1 protein

**Figure S8.**
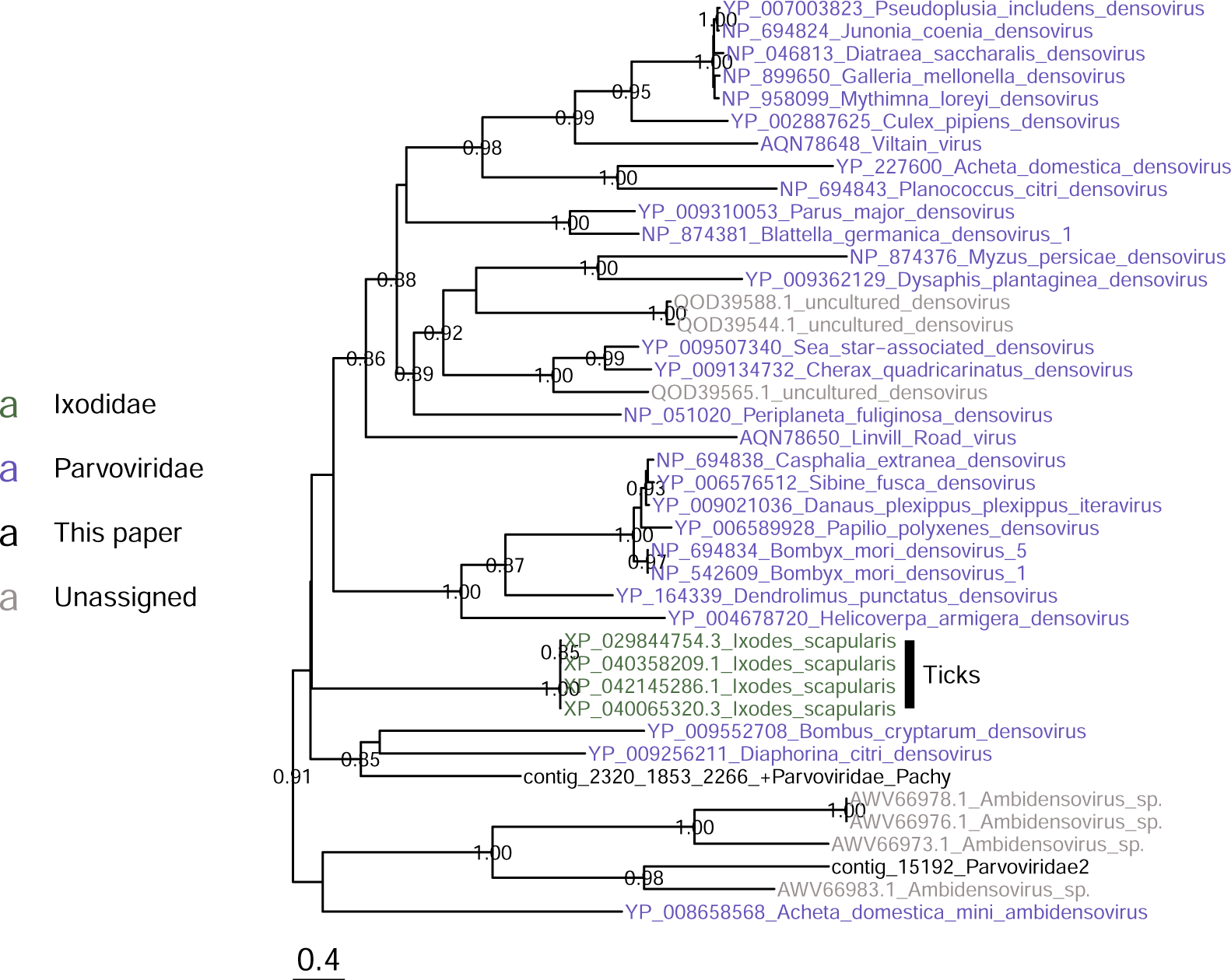
Phylogeny of Parvoviridae based on NS1 protein.

**Figure S9.**
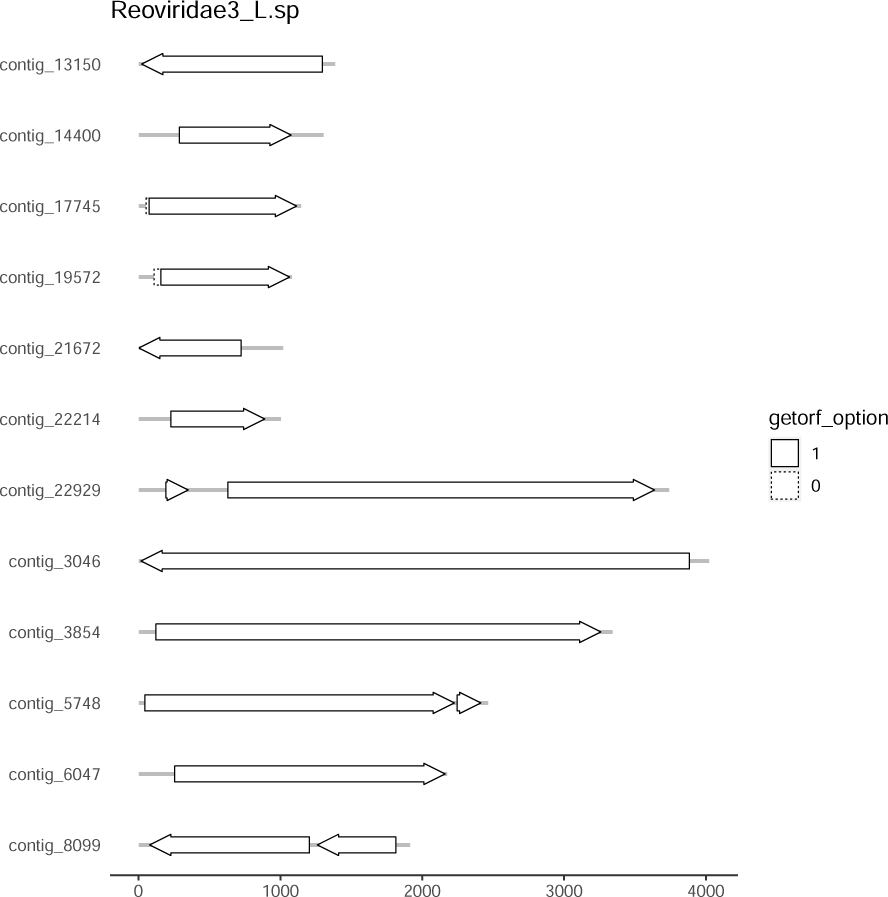
ORF prediction for Reoviridae3_L.sp. Contig_17745 did not show sequence homology with public databases, but was associated with the other contigs, suggesting it is part of this virus genome.

**Figure S10.**
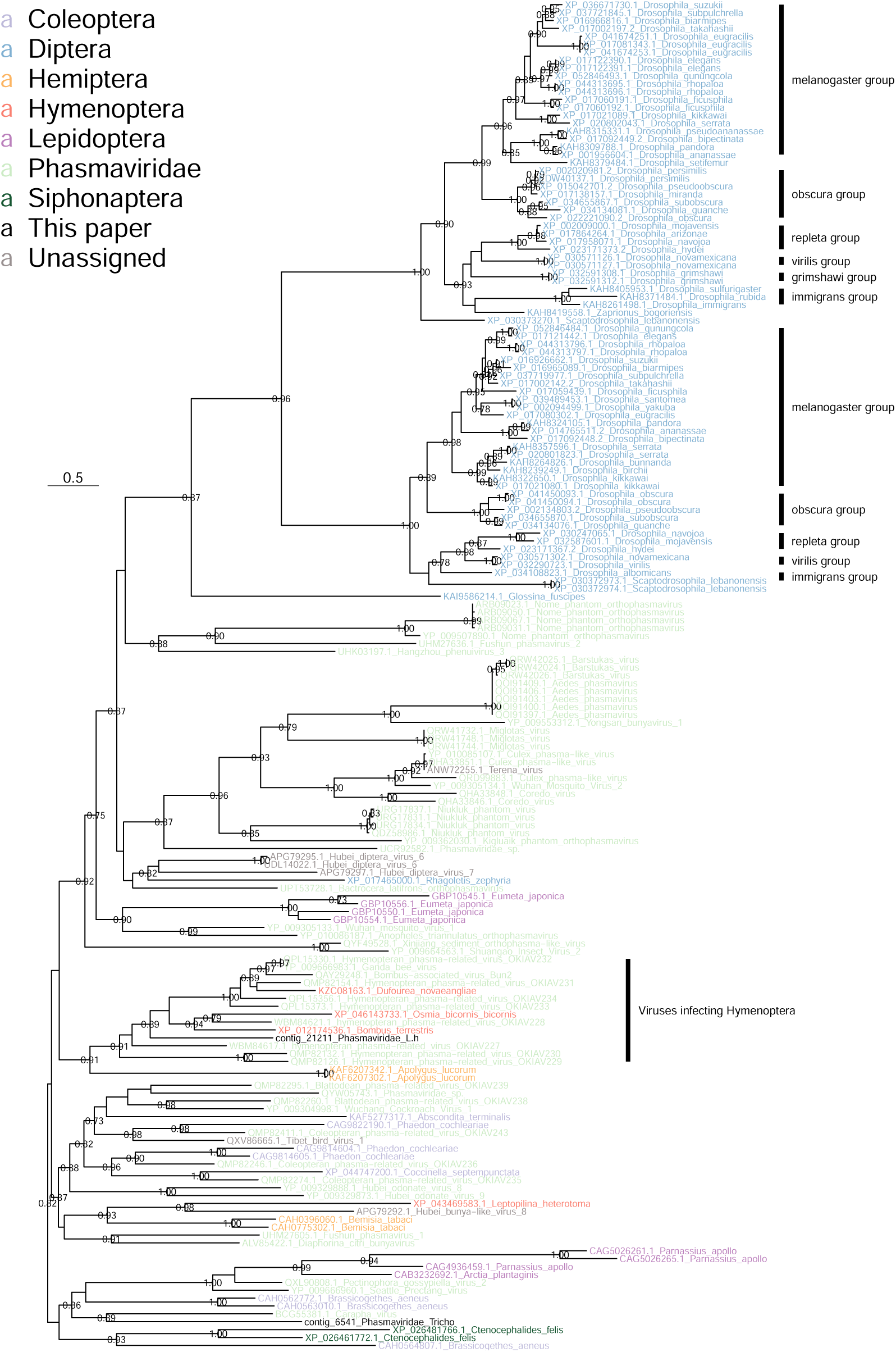
Phylogeny of two new Phasmaviridae detected in *L. heterotoma* and *Trichopria sp.* and evidence for horizontal transfers to various insects, including numerous *Drosophila* species. The nucleoprotein was used to build this phylogeny.

**Figure S11.**
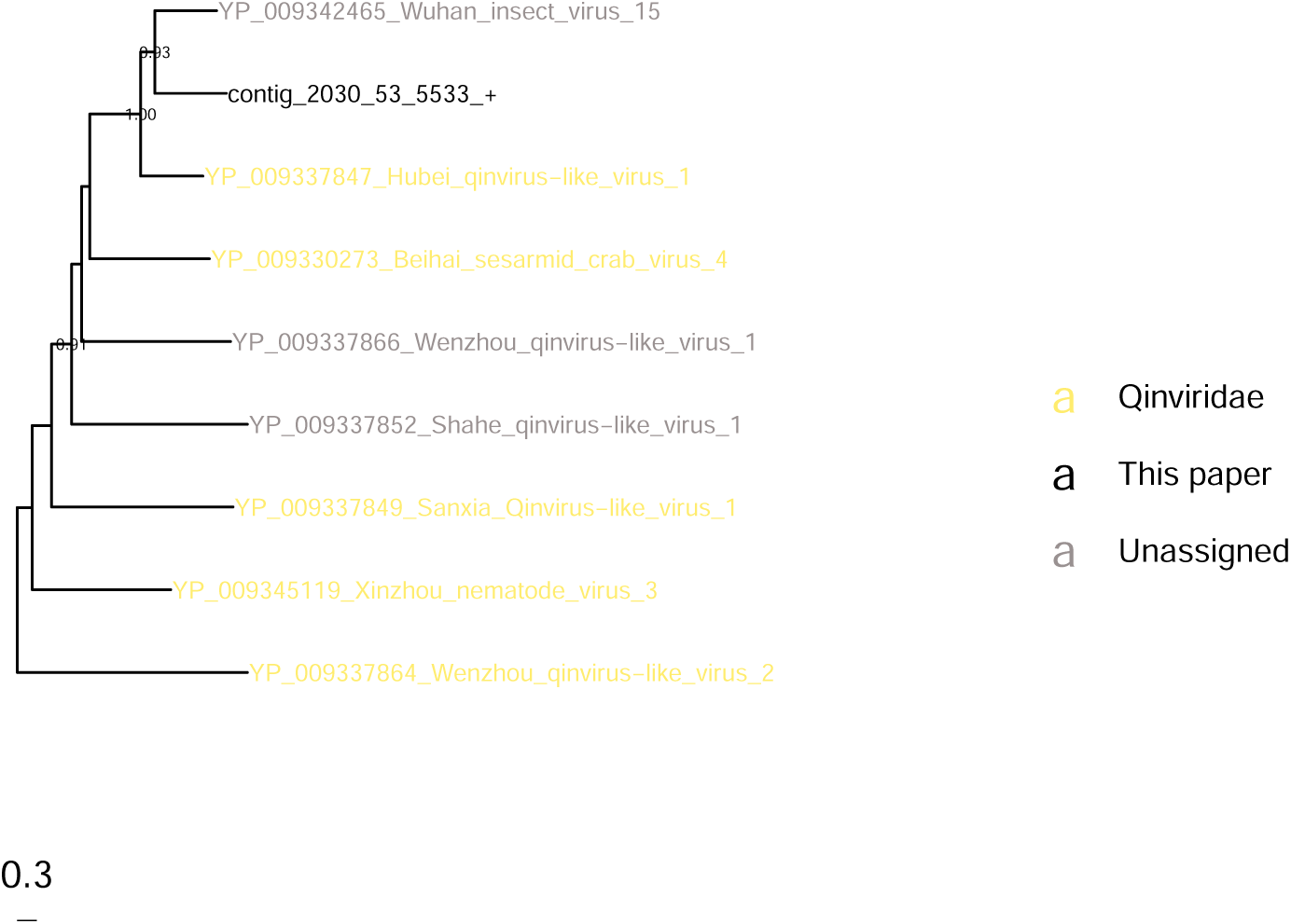
Phylogeny of a new Quinviridae (here called Quinviridae_L.h) detected in *L. heterotoma* based on RdRp.

**Figure S12.**
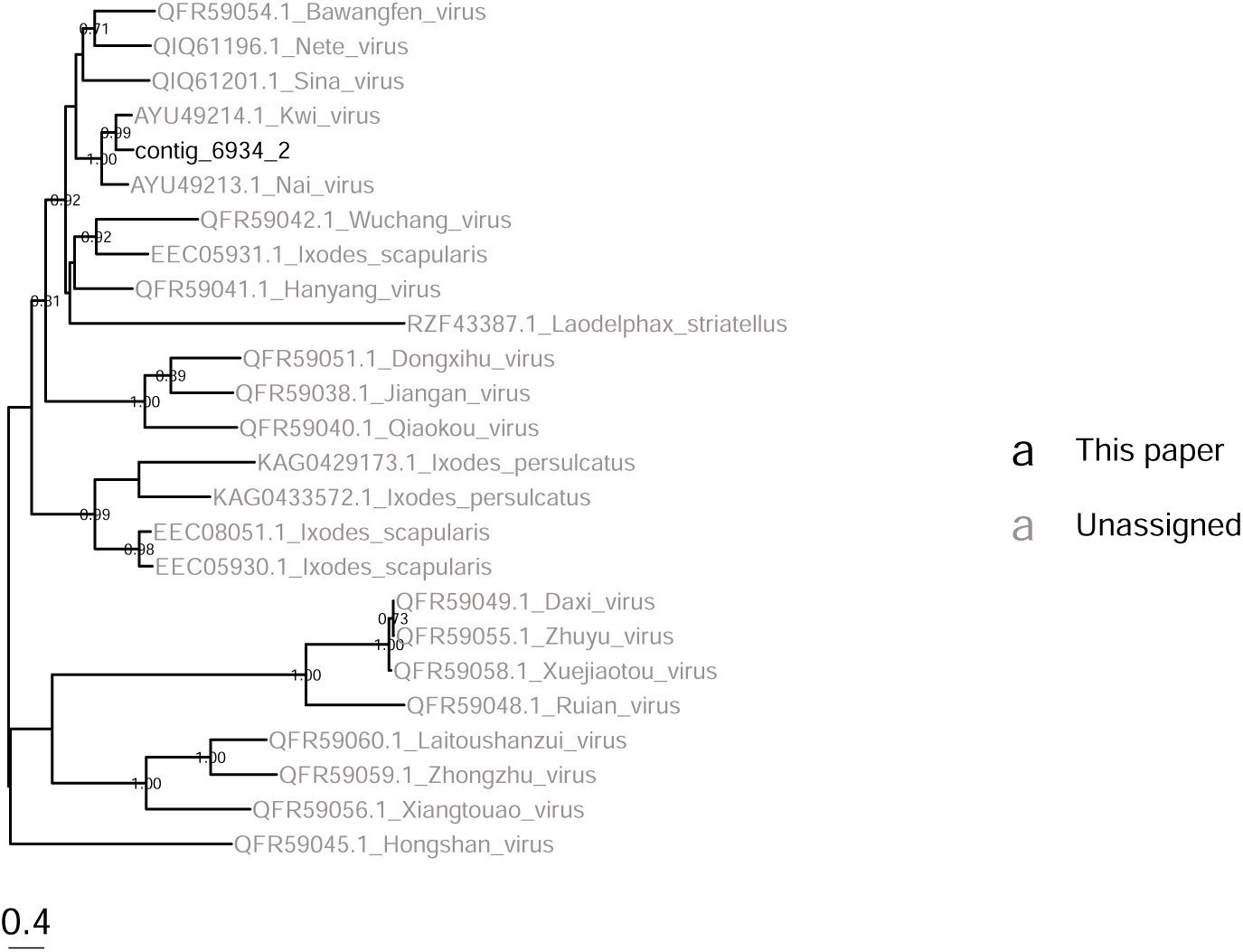
Phylogeny of a new Quenyavirus (denominated Quenyavirus_L.h) detected in *L. heterotoma* based on RdRp.

**Figure S13.**
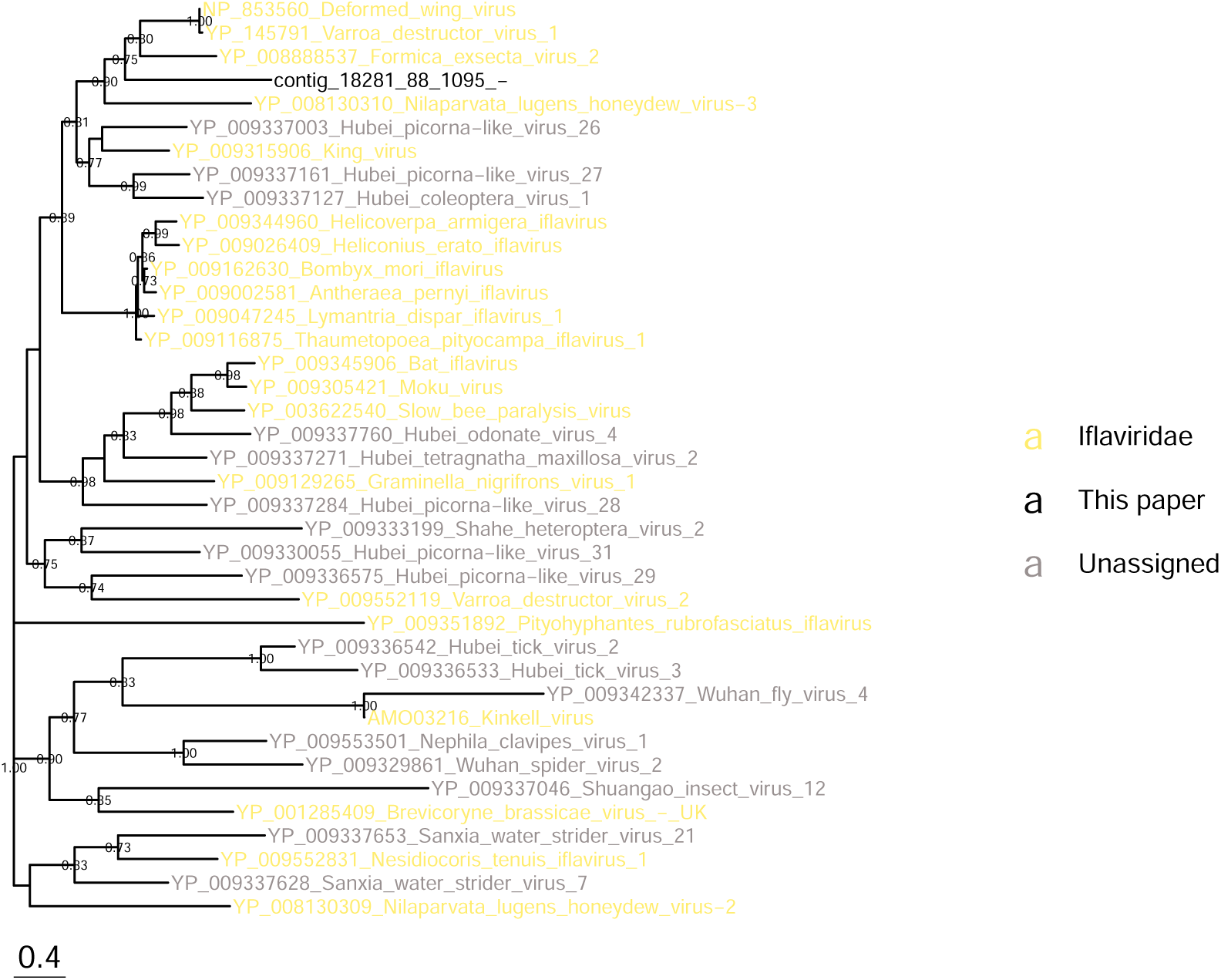
Phylogeny of a new Formicavirus-like (denominated Iflaviridae_L.h) detected in *L. heterotoma*, based on capsid protein.

**Figure S14.**
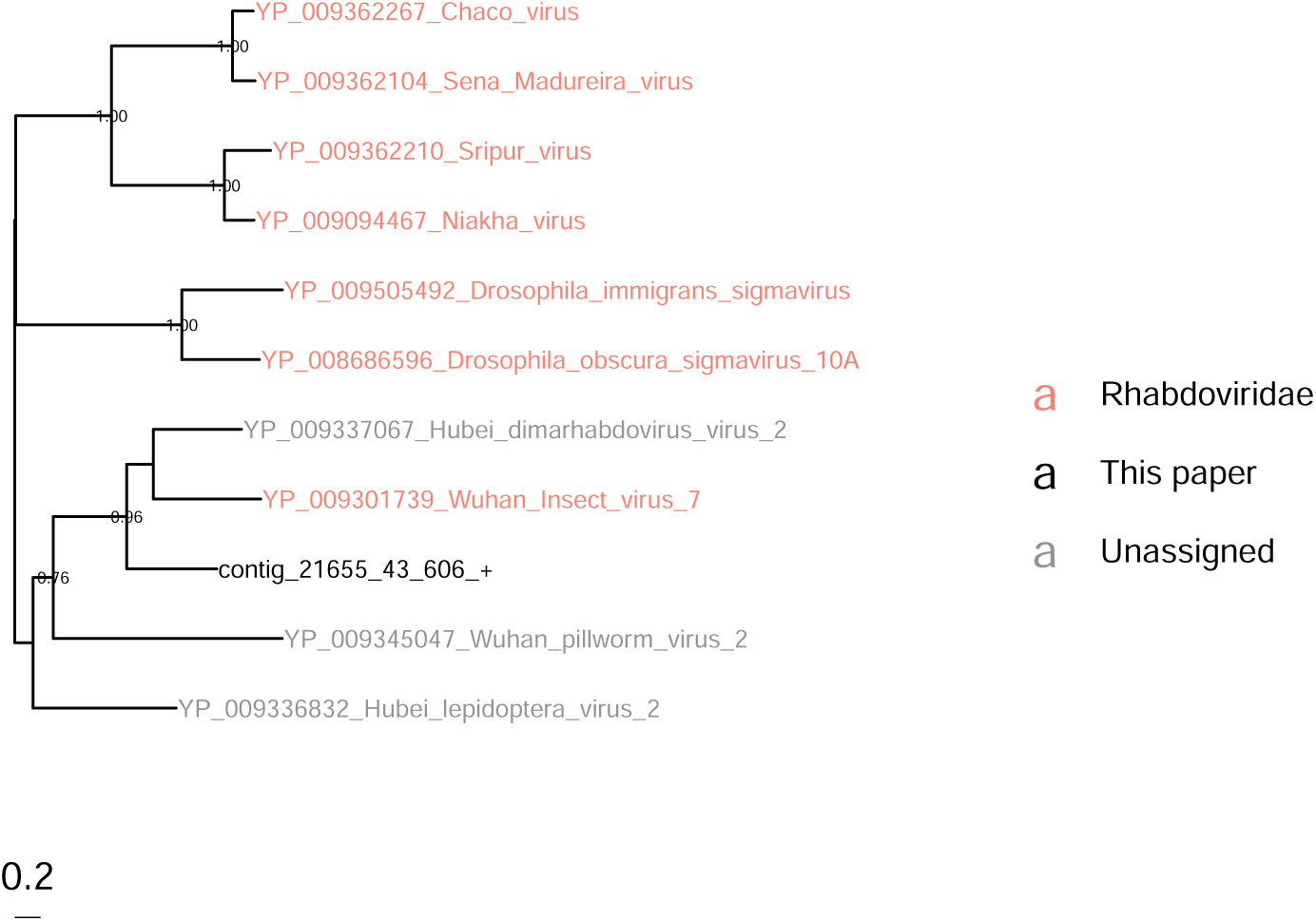
Phylogeny of a new dimarhabdovirus (Rhabdoviridae2) detected in *D. suzukii* and a few other samples based on RdRp.

**Figure S15.**
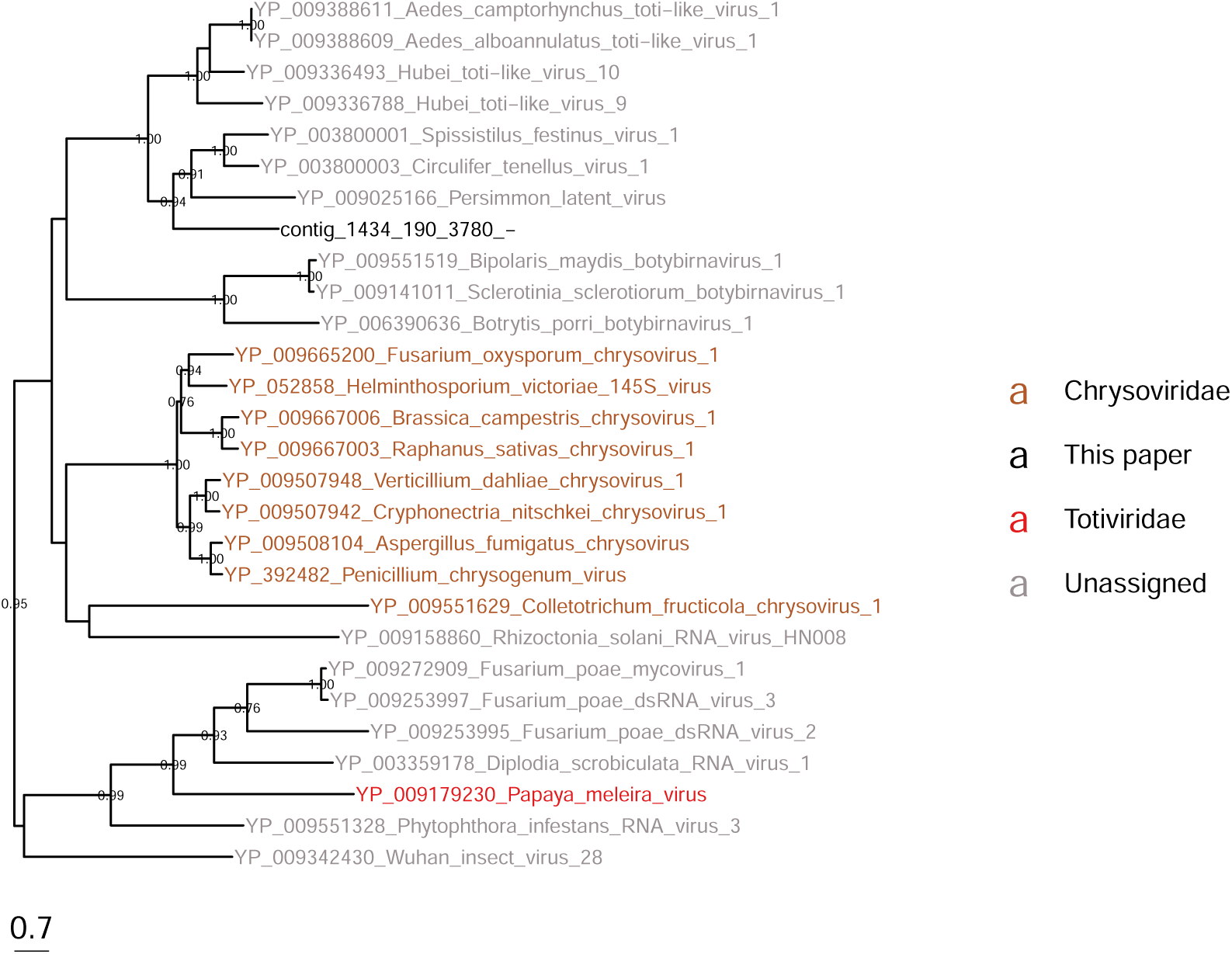
Phylogeny of a new Circulifer-like virus (dsRNA_virus1_Tricho) detected in *Trichopria sp.* based on RdRp.

**Figure S16.**
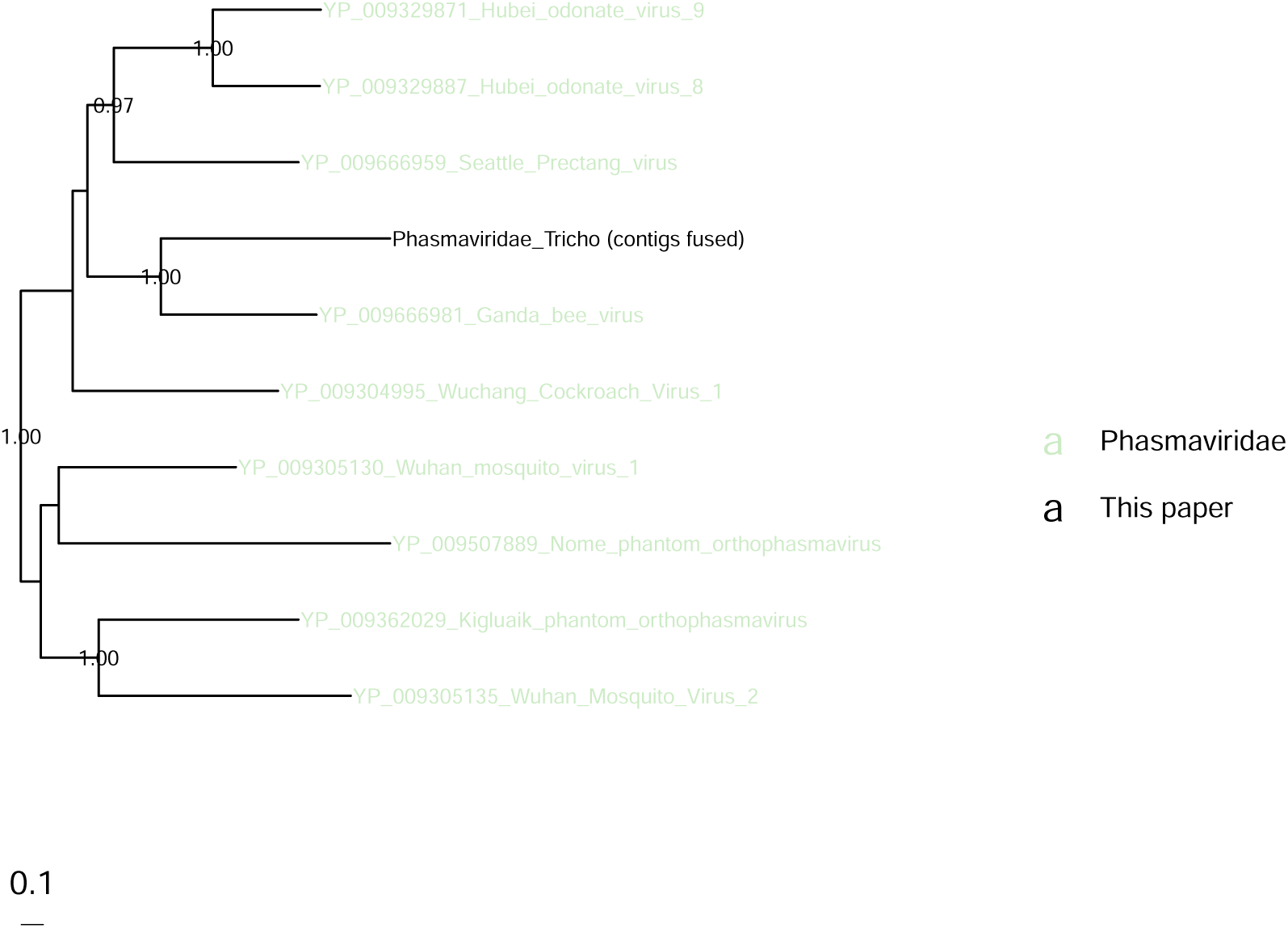
Phylogeny of a new Ganda-like virus (Phasmaviridae_Tricho) detected in *Trichopria sp.* based on RdRp.

**Figure S17.**
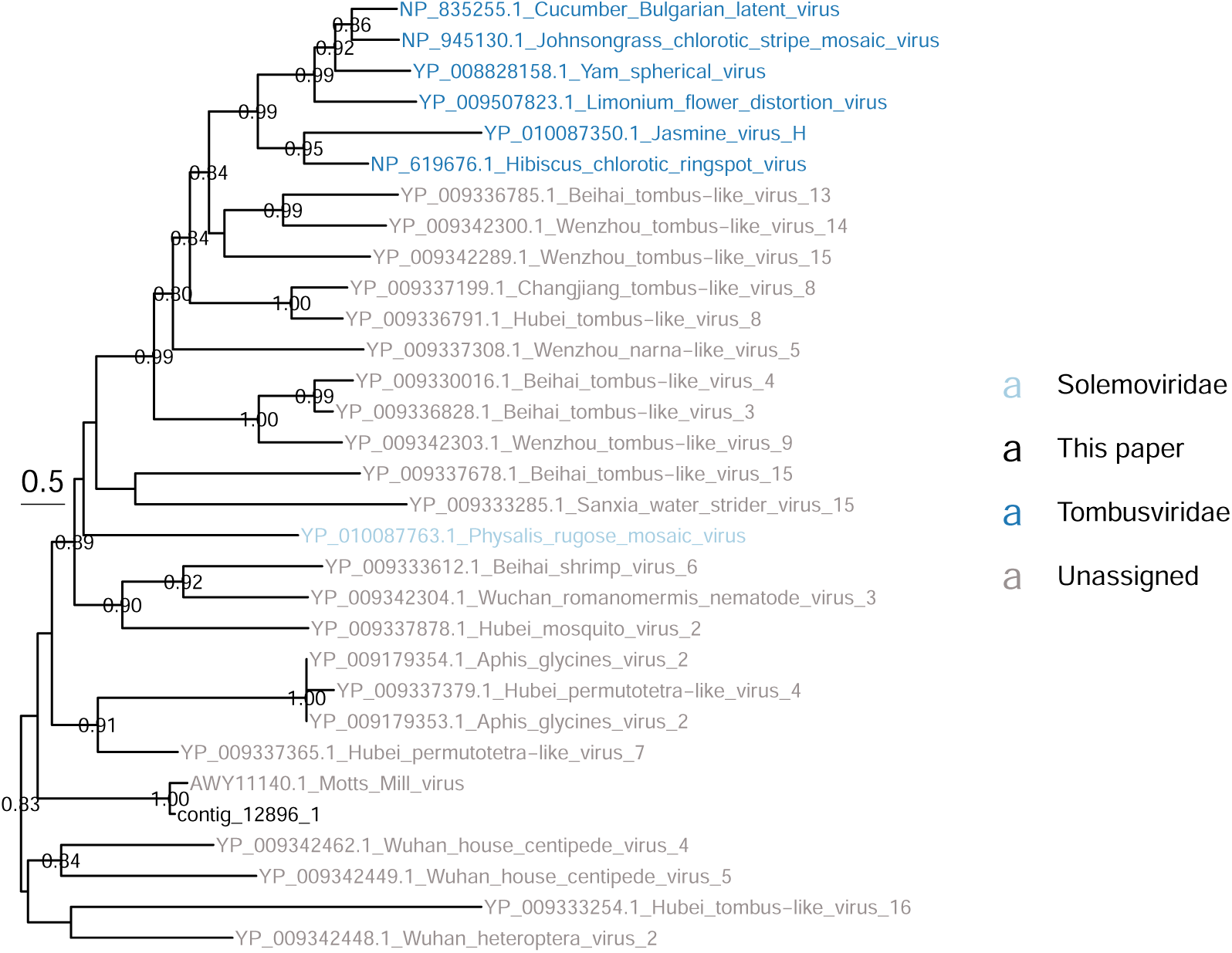
Phylogeny of virus closely related to Motts Mill virus (Motts_Mill_virus_D.sub) detected in *D. subobscura* and possibly *D. obscura* based on putative viral coat protein.

**Figure S18.**
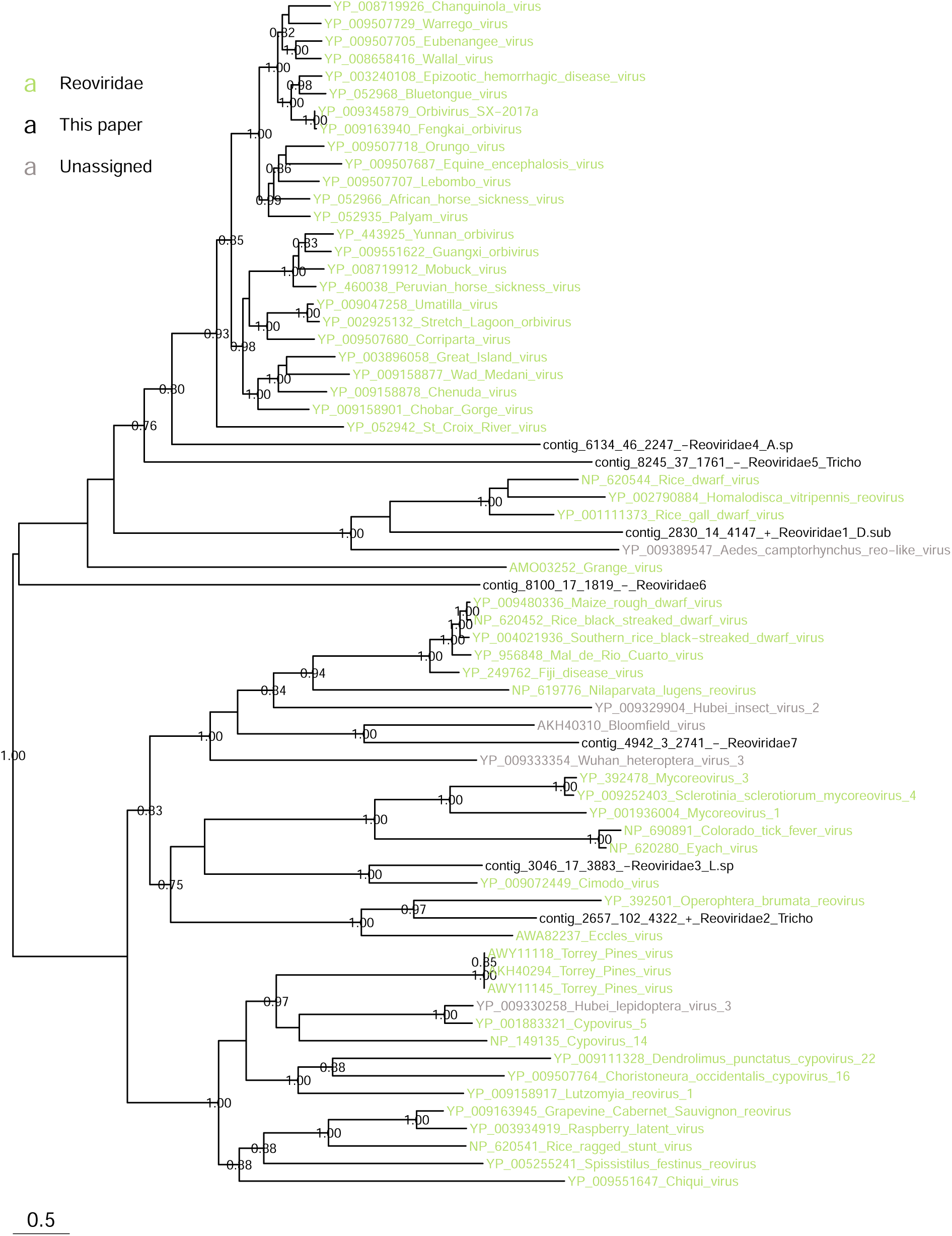
Phylogeny of the Reoviridae detected in this study based on RdRp.

**Figure S19.**
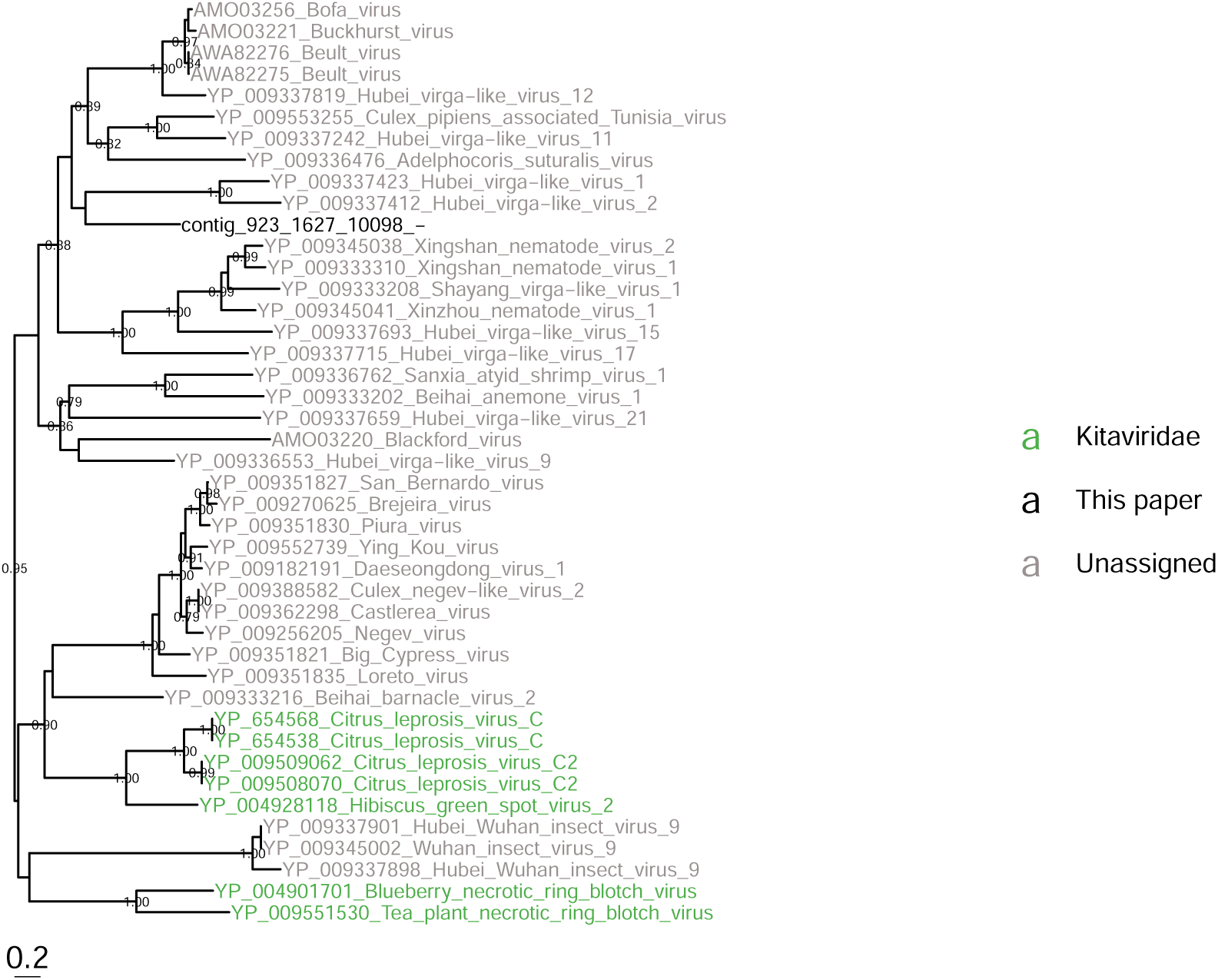
Phylogeny of a new virus (here called "Virga_Tricho") related to Virga viruses detected in *Trichopria sp.* based on RdRp.

**Figure S20.**
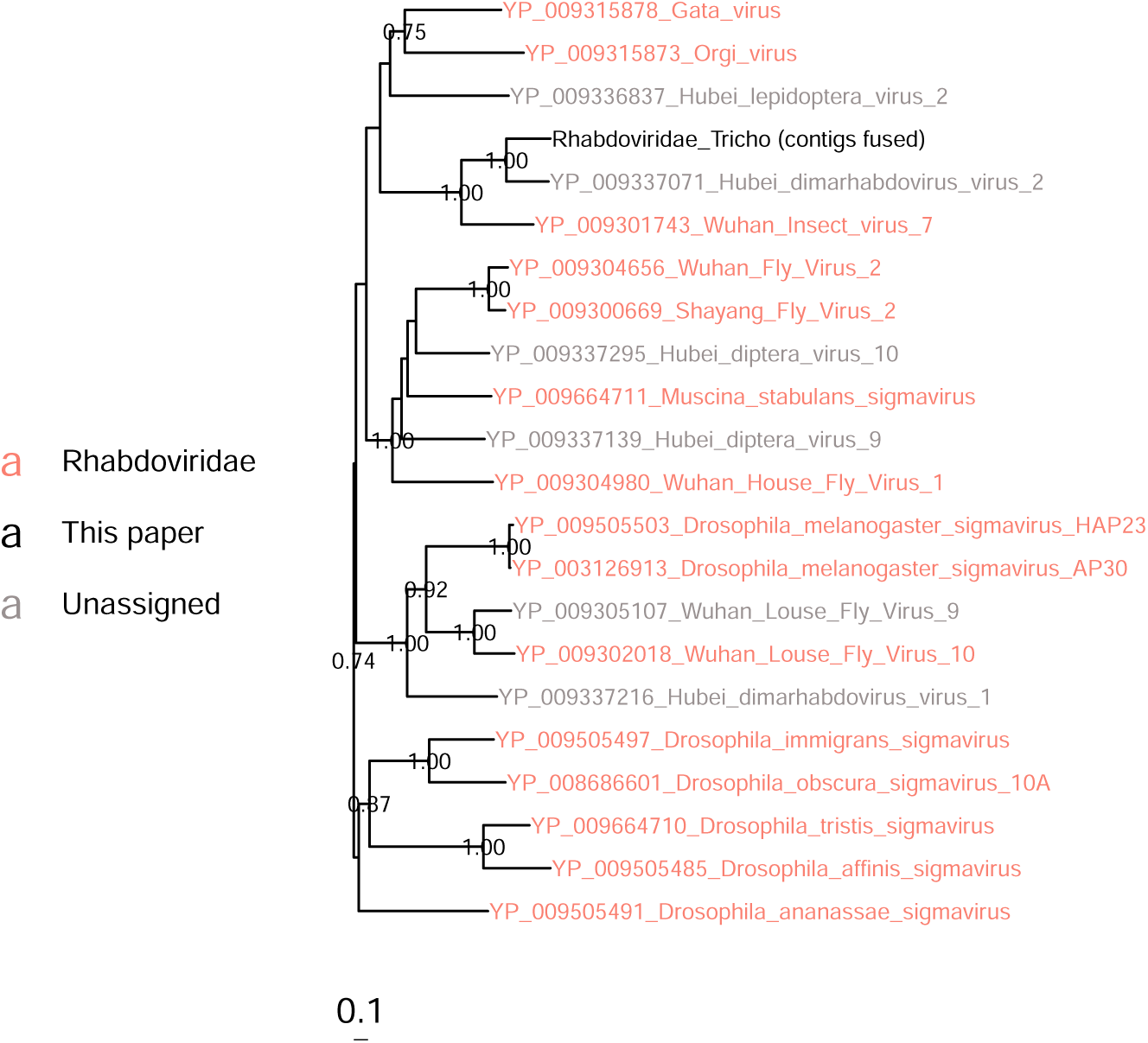
Phylogeny of a new Dimarhabdovirus (provisionnally called "Rhabodviridae_Tricho") detected in *Trichopria sp.* based on RdRp protein.

**Figure S21.**
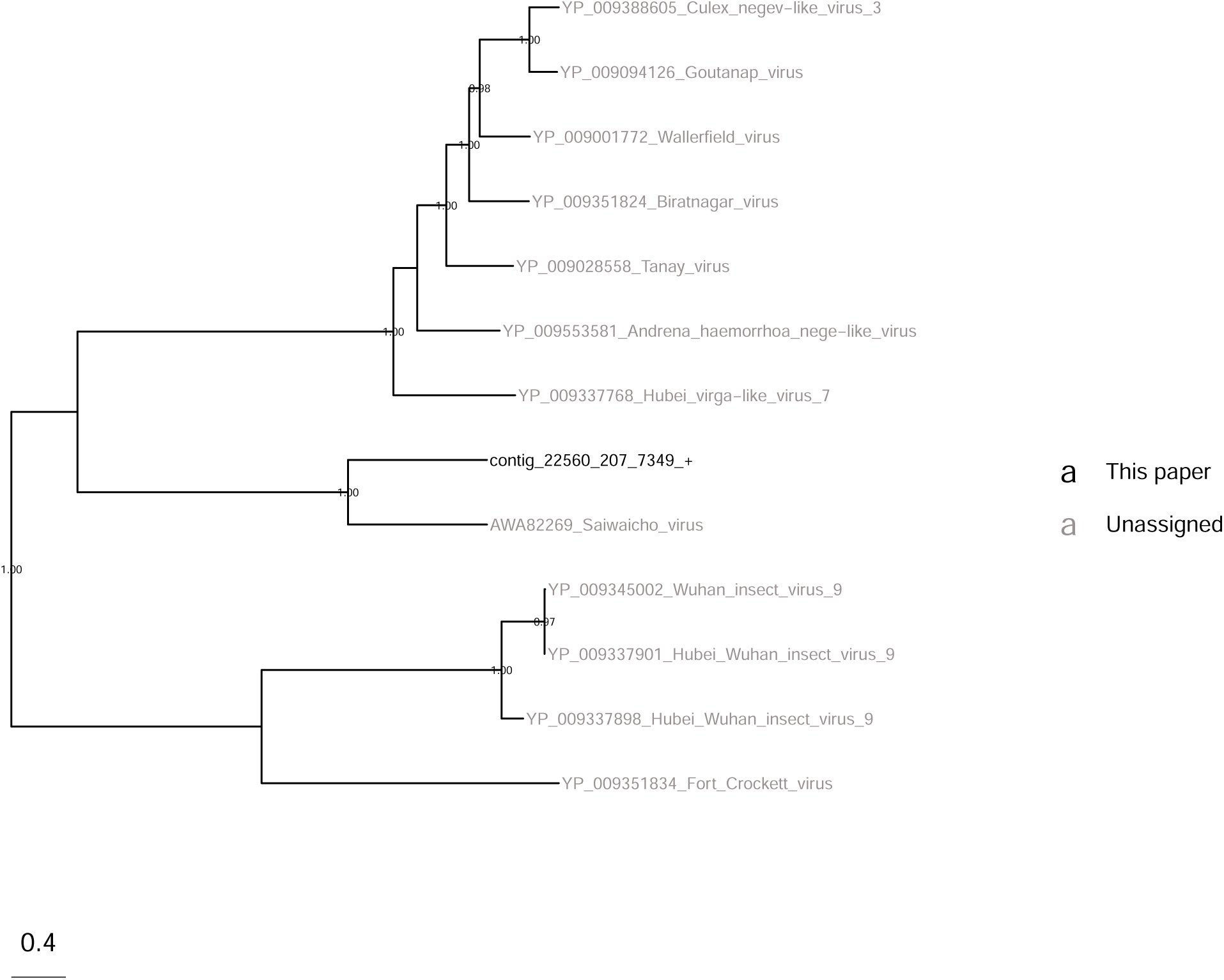
Phylogeny of a new Hepe-Virga virus (denominated Hepe-Virga_L.b in this paper) mostly detected in *L. boulardi* based on RdRp.

**Figure S22.**
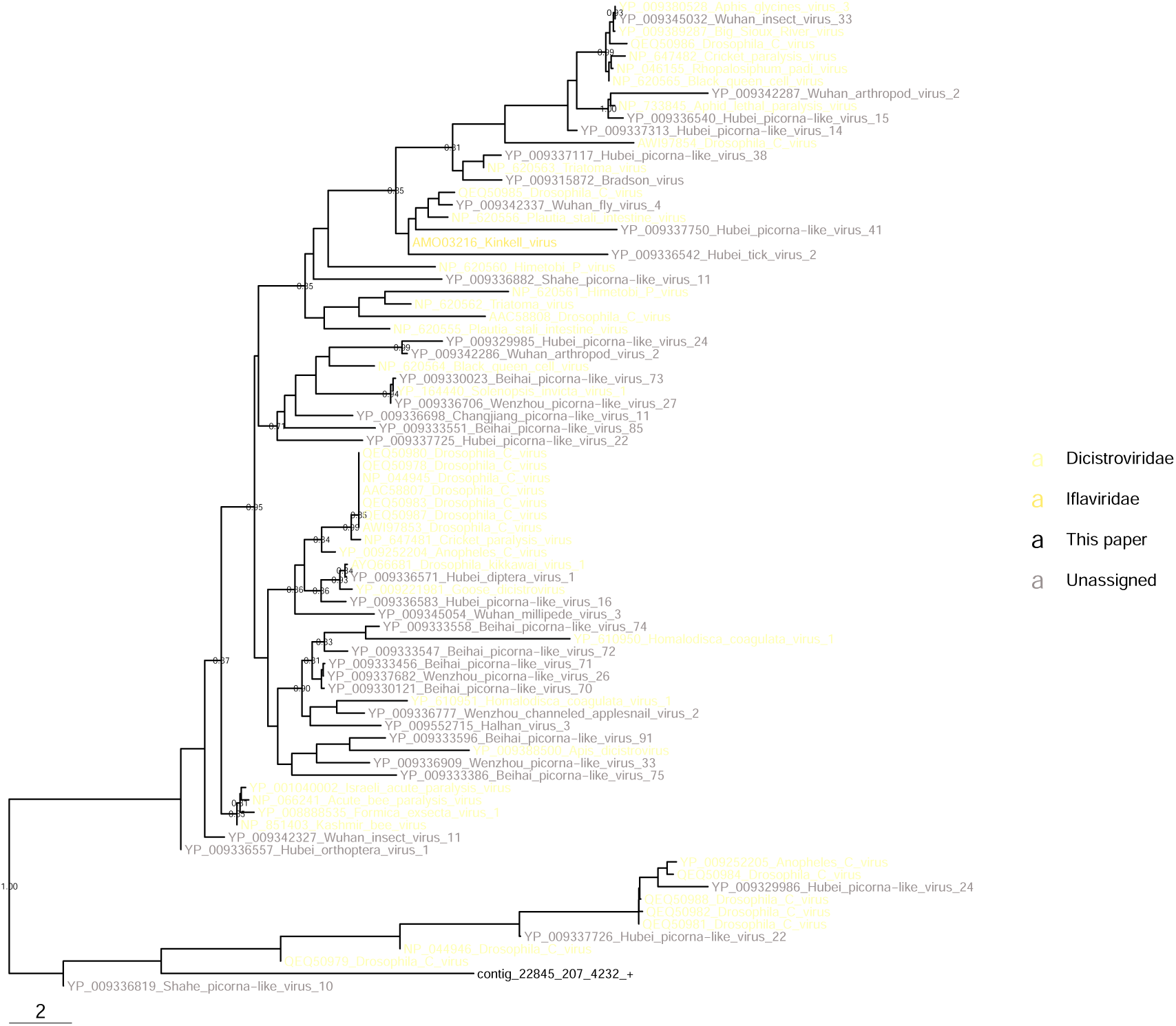
Phylogeny of a new Dicistroviridae (Dicistroviridae_Pachy) detected in *Pachycrepoideus sp* based on RdRp.

**Figure S23.**
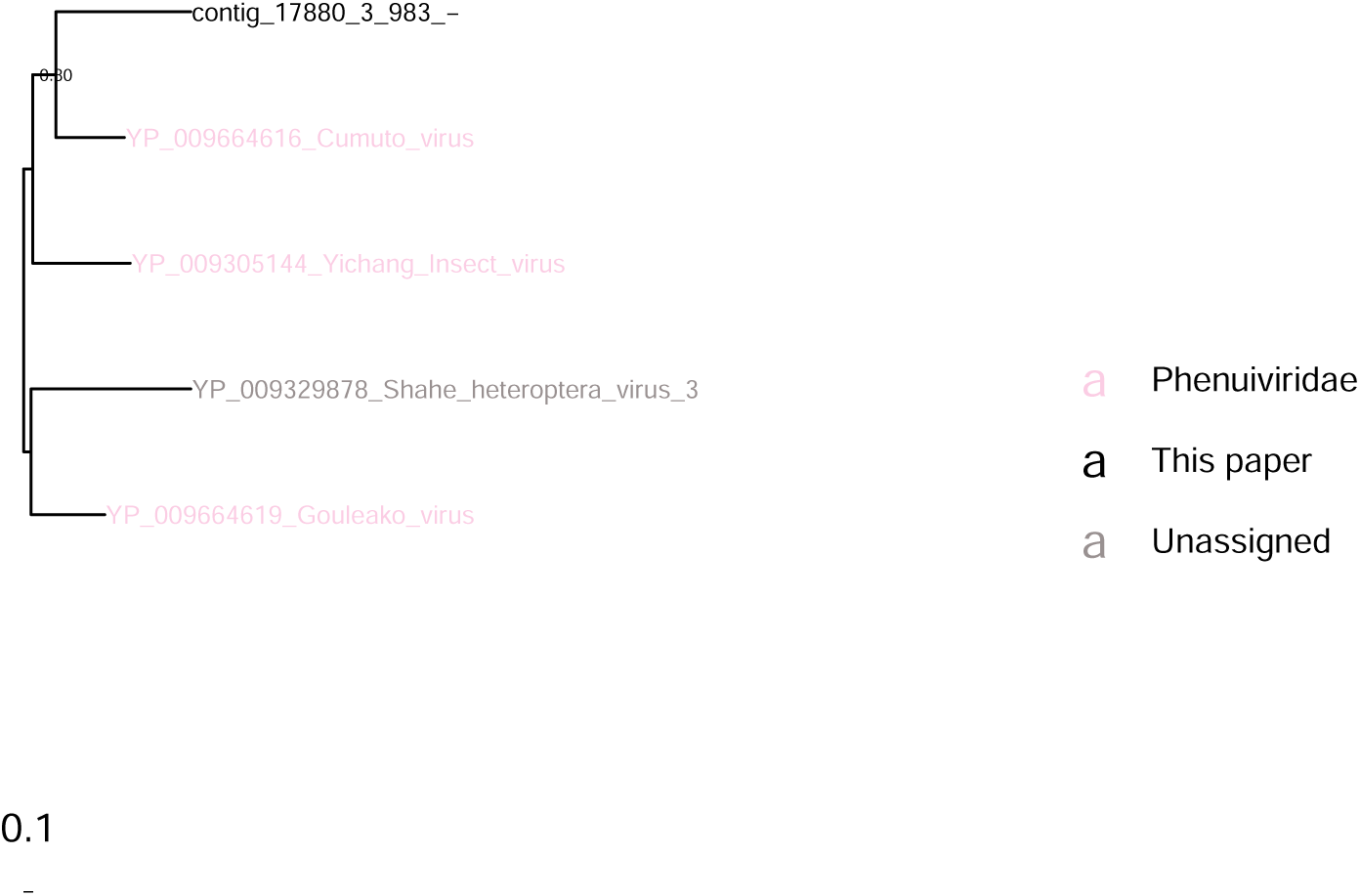
Phylogeny of a new Phenuiviridae (Phenuiviridae_Pachy) detected in *Pachycrepoideus sp* based on glycoprotein G.

**Figure S24.**
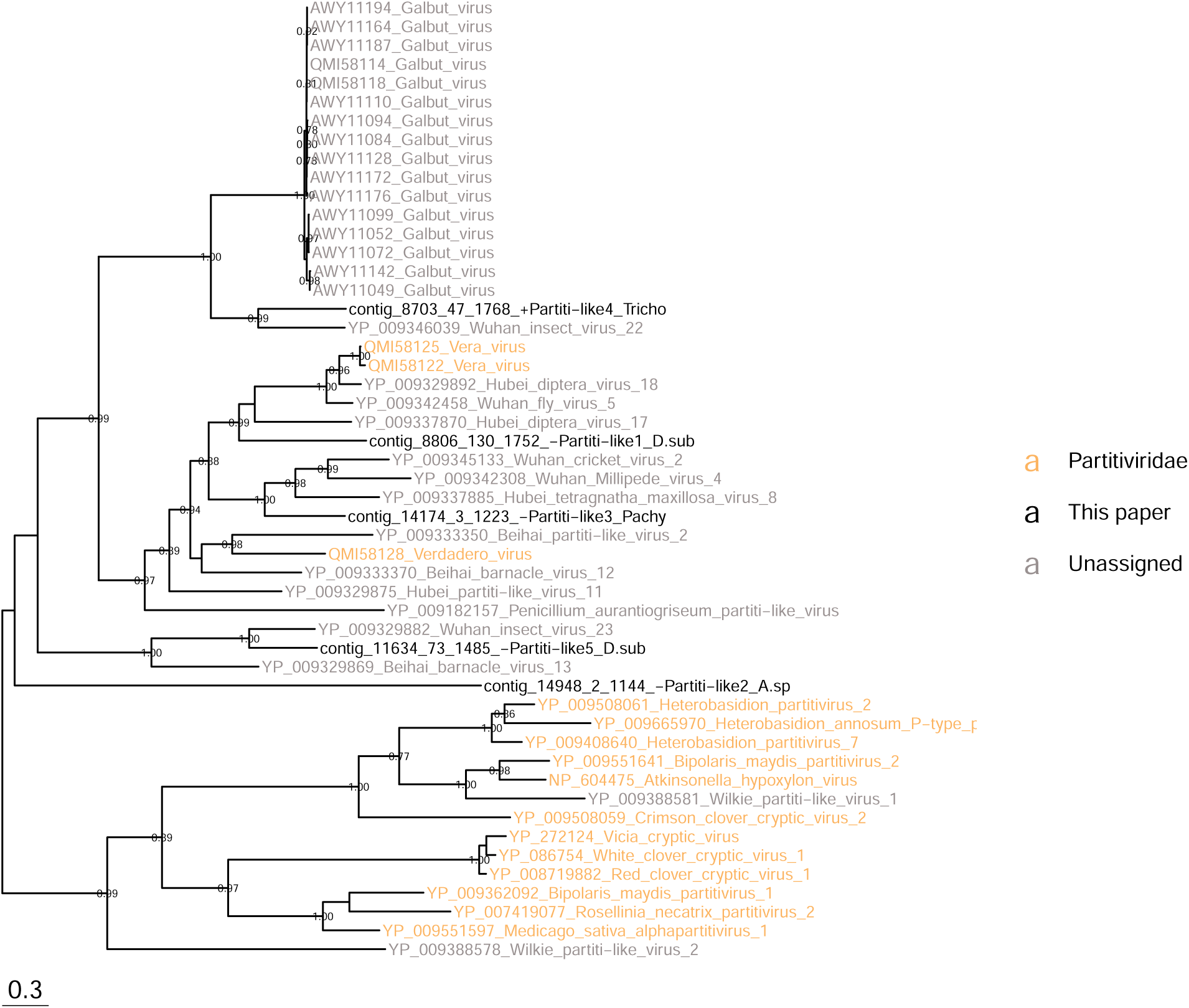
Phylogeny of Partitiviridae based on RdRp protein.

**Figure S25.**
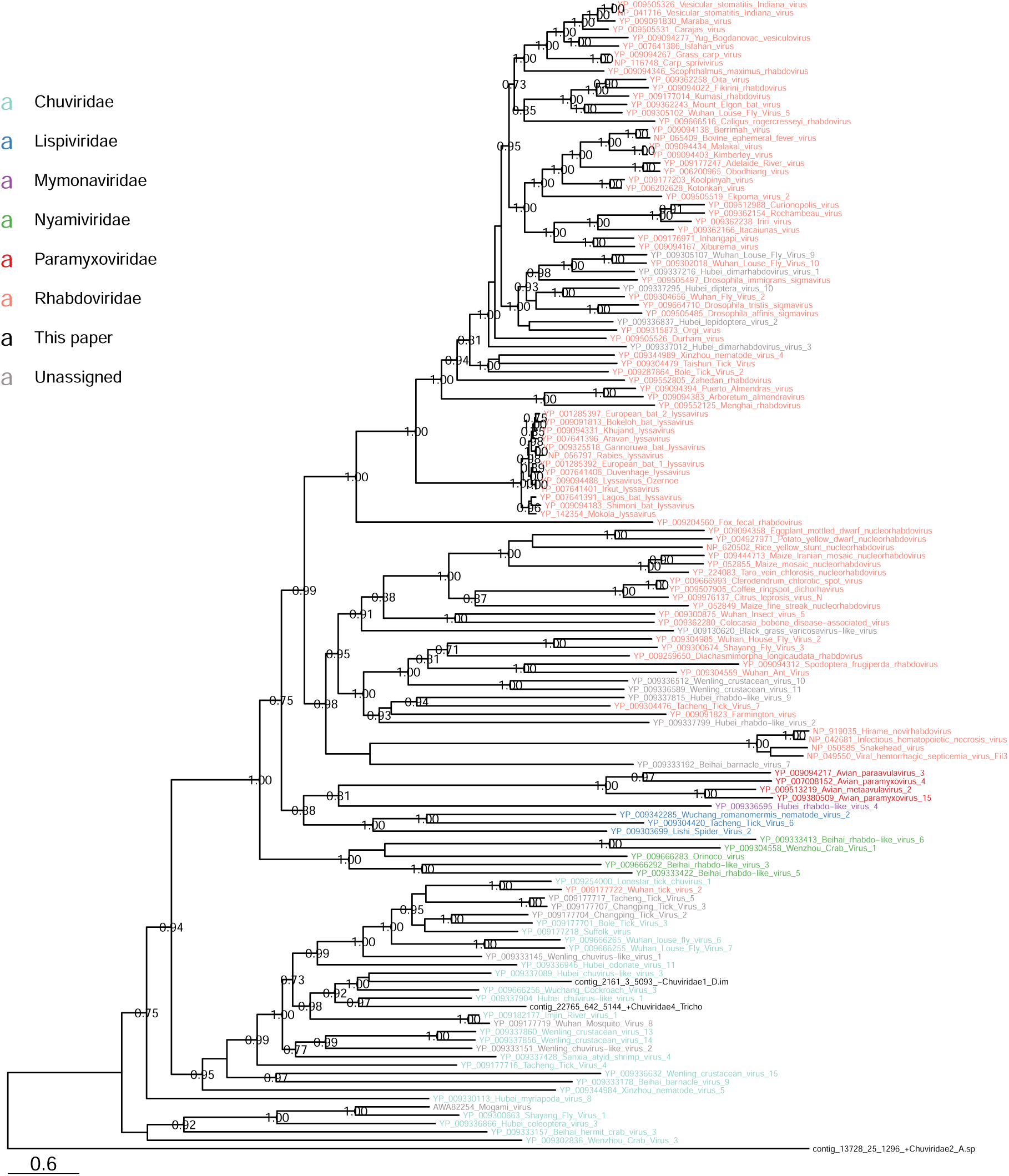
Phylogeny of Chuviridae based on RdRp.

**Figure S26.**
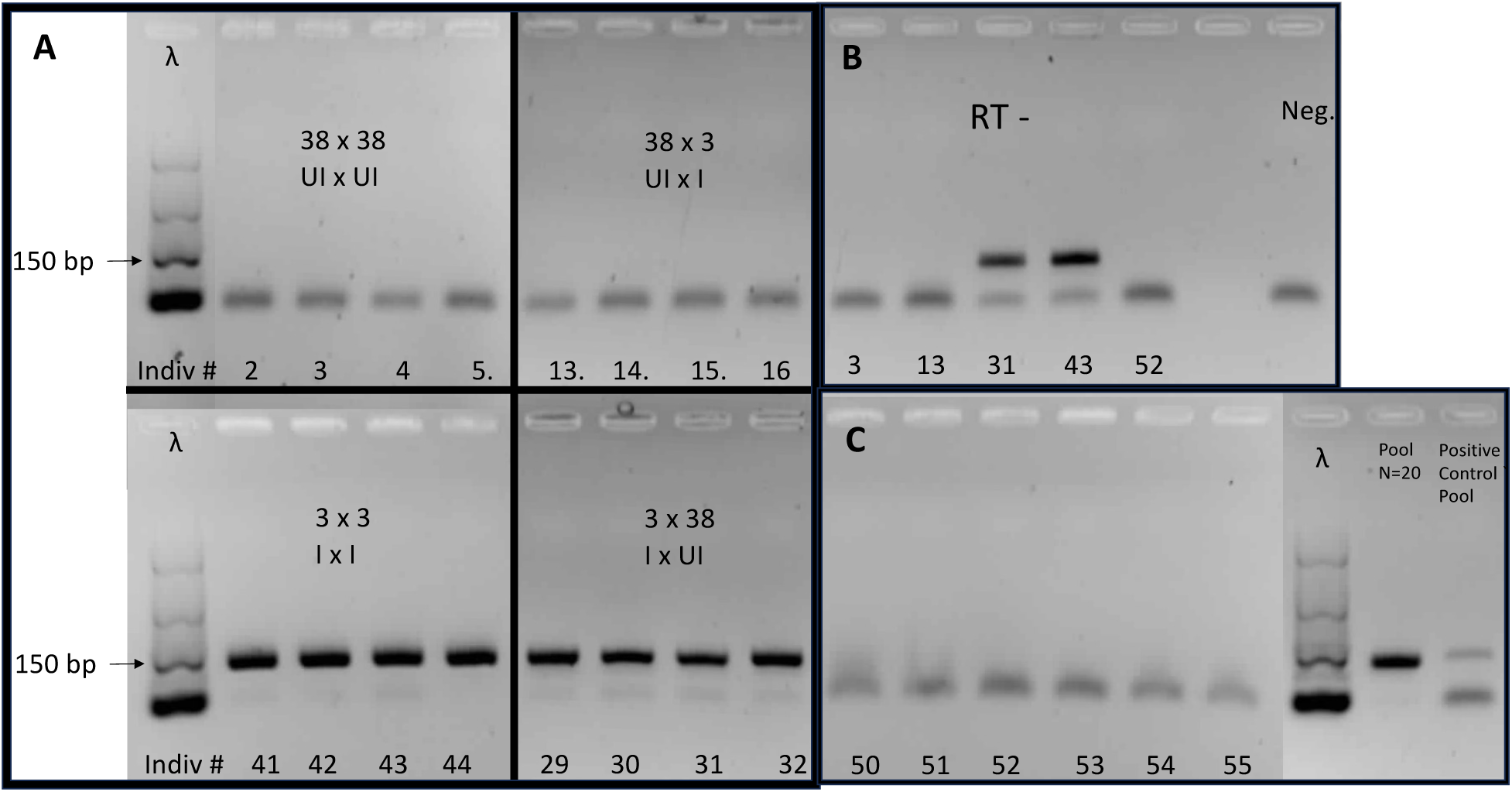
The vertical and horizontal transmission of the Iflaviridae/Lh was studied by rt-PCR using Line 38 (UI: uninfected) and line 3 (I: infected). These lines were collected in Lyon, La Doua. (A) Detection of Iflaviridae_Lh by rt-PCR in the four crosses of the parasitoid *L. heterotoma* (female x male) (B) Negative control (Neg.) and controls without reverse-transcription (RT-) from five individuals extracts (individual number is indicated by a number at the bottom of each lane). (C) Results of the rt-PCR obtained for six individuals (individuals 50 to 55) emerged from the horizontal transmission assay. A positive control pool of 1 infected individual mixed with 19 uninfected individuals (Positive Control pool) revealed rt-PCR amplification as expected. The "Pool N=20" lane corresponds to a pool of 20 test individuals emerged from the horizontal transmission assay. The ladder is the O’RangeRuler 50 bp from ThermoFisher (*λ* lane).

**Figure S27.**
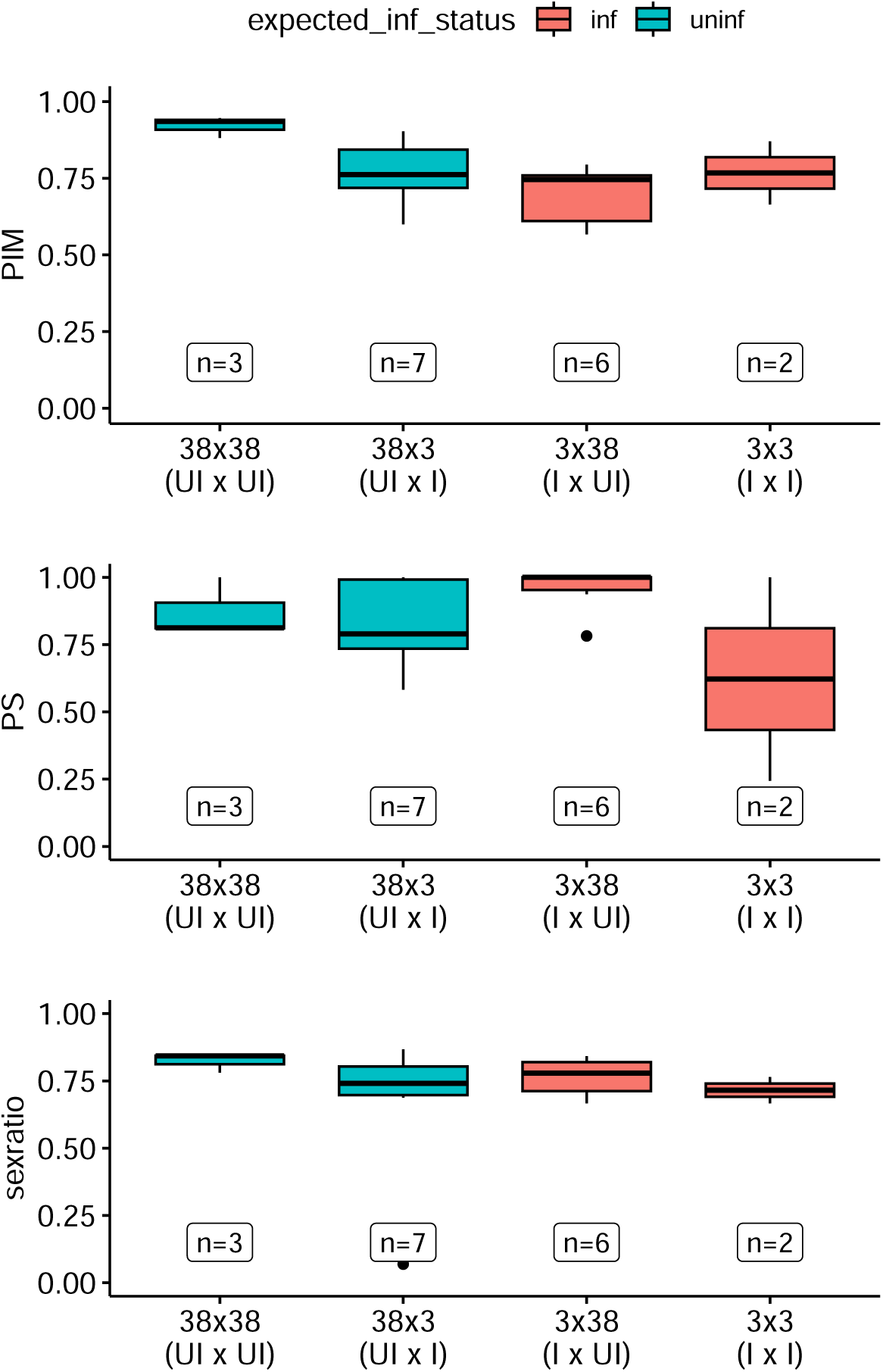
Phenotype assay in *L. heterotoma* infected or not by Iflaviridae_Lh. Crosses are indicated as female x male. UI stands for uninfected and I for infected. Sample sizes are indicated as n=x.

**Figure S28.**
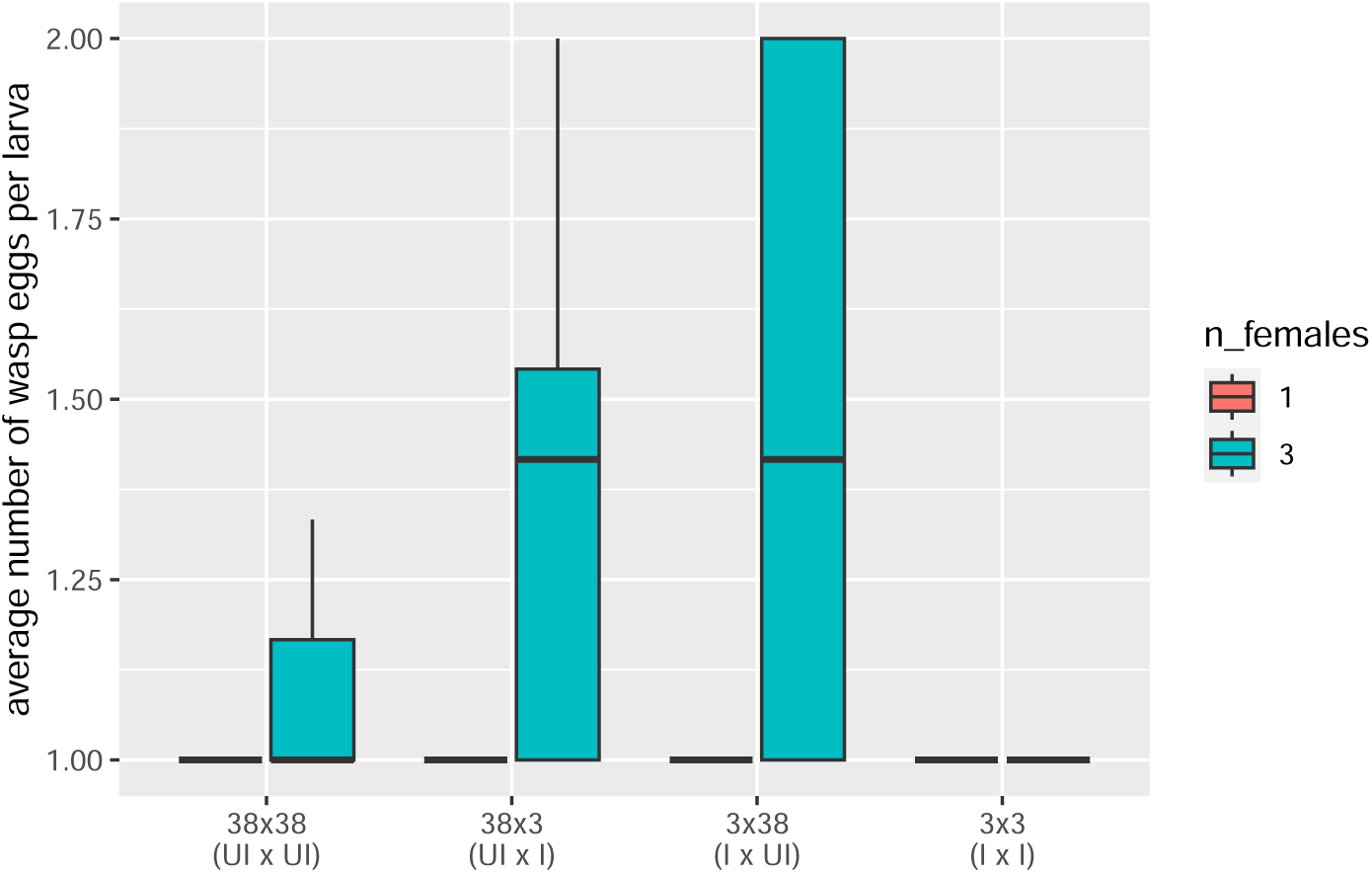
Superparasitism was low and independent of infection by Iflaviridae_L.h in *L. heterotoma*. One or three female wasps were provided with 10 *Drosophila* larvae for 48 hours. Crosses are indicated as female x male. UI stands for uninfected and I for infected. Sample sizes are the following from left to right: n=4, n=8, n=8, n=4 (density 1) and n=3, n=8, n=8, n=2 (density 3).

## References

1. Manzano-Marín A., Oceguera-Figueroa A., Latorre A., Jiménez-García L. F. & Moya A. Solving a Bloody Mess: B-Vitamin Independent Metabolic Convergence among Gammaproteobacterial Obligate Endosymbionts from Blood-Feeding Arthropods and the Leech *Haementeria Officinalis*. Genome Biology and Evolution 7, 2871–2884. ISSN: 1759-6653 (Oct. 2015).

2. Sudakaran S., Kost C. & Kaltenpoth M. Symbiont Acquisition and Replacement as a Source of Ecological Innovation. Trends in Microbiology 25, 375–390. ISSN: 0966842X (May 2017).

3. Engelstädter J., Hammerstein P. & Hurst G. D. D. The Evolution of Endosymbiont Density in Doubly Infected Host Species. Journal of Evolutionary Biology 20, 685–695. ISSN: 1010-061X, 1420-9101 (Mar. 2007).

4. Douglas A. E. Symbiotic Microorganisms: Untapped Resources for Insect Pest Control. Trends in Biotechnology 25, 338–342. ISSN: 01677799 (Aug. 2007).

5. Moran N. A., McCutcheon J. P. & Nakabachi A. Genomics and Evolution of Heritable Bacterial Symbionts. Annual Review of Genetics 42, 165–190. ISSN: 0066-4197, 1545-2948 (Dec. 2008).

6. Martoni F. et al. Insect Phylogeny Structures the Bacterial Communities in the Microbiome of Psyllids (Hemiptera: Psylloidea) in Aotearoa New Zealand. PLOS ONE 18 (ed Kalendar R.) e0285587. ISSN: 1932-6203 (May 2023).

7. McLean A. H. C., Godfray H. C. J., Ellers J. & Henry L. M. Host Relatedness Influences the Composition of Aphid Microbiomes. Environmental Microbiology Reports 11, 808–816. ISSN: 1758-2229 (2019).

8. Martinson V. G., Douglas A. E. & Jaenike J. Community Structure of the Gut Microbiota in Sympatric Species of Wild Drosophila. Ecology Letters 20, 629–639. ISSN: 1461-0248 (2017).

9. Tsuchida T., Koga R. & Fukatsu T. Host Plant Specialization Governed by Facultative Symbiont. Science 303, 1989–1989. ISSN: 0036-8075, 1095-9203 (Mar. 2004).

10. Ferrari J., West J. A., Via S. & Godfray H. C. J. Population Genetic Structure and Secondary Symbionts in Host-Associated Populations of the Pea Aphid Complex: Population Genetics and Symbionts in the Pea Aphid. Evolution 66, 375–390. ISSN: 00143820 (Feb. 2012).

11. Wu H., et al. Abundant and Diverse RNA Viruses in Insects Revealed by RNA-Seq Analysis: Ecological and Evolutionary Implications. mSystems 5 (ed Cristea I. M.) ISSN: 2379-5077 (Aug. 2020).

12. Shi M. et al. Redefining the Invertebrate RNA Virosphere. Nature 540, 539–543. ISSN: 0028-0836, 1476-4687 (Dec. 2016).

13. Edgar R. C. et al. Petabase-Scale Sequence Alignment Catalyses Viral Discovery. Nature 602, 142–147. ISSN: 0028-0836, 1476-4687 (Feb. 2022).

14. Márquez L. M., Redman R. S., Rodriguez R. J. & Roossinck M. J. A Virus in a Fungus in a Plant: Three-Way Symbiosis Required for Thermal Tolerance. Science 315, 513–515. ISSN: 0036-8075, 1095-9203 (Jan. 2007).

15. Xu P. et al. Novel Partiti-like Viruses Are Conditional Mutualistic Symbionts in Their Normal Lepidopteran Host, African Armyworm, but Parasitic in a Novel Host, Fall Armyworm. PLOS Pathogens 16, e1008467. ISSN: 1553-7374 (June 2020).

16. Guinet B. et al. Endoparasitoid Lifestyle Promotes Endogenization and Domestication of dsDNA Viruses. eLife 12, e85993. ISSN: 2050-084X (June 2023).

17. Di Giovanni D. et al. A Behavior-Manipulating Virus Relative as a Source of Adaptive Genes for Drosophila Parasitoids. Molecular Biology and Evolution 37, 2791–2807. ISSN: 1537-1719 (Oct. 2020).

18. Gauthier J., Drezen J.-M. & Herniou E. A. The Recurrent Domestication of Viruses: Major Evolutionary Transitions in Parasitic Wasps. Parasitology 145, 713–723. ISSN: 0031-1820, 1469-8161 (May 2018).

19. Varaldi J. et al. Infectious Behavior in a Parasitoid. Science 302, 1930–1930. ISSN: 0036-8075, 1095-9203 (Dec. 2003).

20. Varaldi J. et al. Artifical Transfer and Morphological Description of Virus Particles Associated with Super-parasitism Behaviour in a Parasitoid Wasp. Journal of Insect Physiology, 1202–1212. ISSN: 00221910 (Nov. 2006).

21. Kageyama D. A Male-Killing Gene Encoded by a Symbiotic Virus of Drosophila. Nature Communications (2023).

22. Patot S., Allemand R., Fleury F. & Varaldi J. An Inherited Virus Influences the Coexistence of Parasitoid Species through Behaviour Manipulation: A Symbiont Mediates Interspecific Competition. Ecology Letters 15, 603–610. ISSN: 1461023X (June 2012).

23. Halbach R., Junglen S. & Van Rij R. P. Mosquito-Specific and Mosquito-Borne Viruses: Evolution, Infection, and Host Defense. Current Opinion in Insect Science 22, 16–27. ISSN: 22145745 (Aug. 2017).

24. McMenamin A. J. & Genersch E. Honey Bee Colony Losses and Associated Viruses. Current Opinion in Insect Science 8, 121–129. ISSN: 22145745 (Apr. 2015).

25. Medd N. C. et al. The Virome of Drosophila Suzukii, an Invasive Pest of Soft Fruit. Virus Evolution 4. ISSN: 2057-1577 (Jan. 2018).

26. Thongsripong P. et al. Metagenomic Shotgun Sequencing Reveals Host Species as an Important Driver of Virome Composition in Mosquitoes. Scientific Reports 11, 8448. ISSN: 2045-2322 (Dec. 2021).

27. Bergner L. M. et al. Demographic and Environmental Drivers of Metagenomic Viral Diversity in Vampire Bats. Molecular Ecology 29, 26–39. ISSN: 1365-294X (2020).

28. Allemand R., Fleury F., Lemaitre C. & Boulétreau M. Dynamique Des Populations et Interactions Competitives Chez Deux Espèces de Leptopilina, Parasitoïdes de Drosophiles, Dans La Vallee Du Rhône (Hymenoptera: Figitidae). Annales De La Societe Entomologique De France (1999).

29. Fleury F., Gibert P., Ris N. & Allemand R. In Advances in Parasitology 3–44 (Elsevier, 2009). ISBN: 978-0-12-374792-1.

30. Carpenter J. A., Obbard D. J., Maside X. & Jiggins F. M. The Recent Spread of a Vertically Transmitted Virus through Populations of Drosophila Melanogaster. Molecular Ecology 16, 3947–3954. ISSN: 0962-1083, 1365-294X (Sept. 2007).

31. Patot S. et al. Prevalence of a Virus Inducing Behavioural Manipulation near Species Range Border. Molecular Ecology 19, 2995–3007. ISSN: 09621083, 1365294X (June 2010).

32. Lawrence P. O. Non-Poly-DNA Viruses, Their Parasitic Wasps, and Hosts. Journal of Insect Physiology 51, 99–101. ISSN: 00221910 (Feb. 2005).

33. De Buron I. & Beckage N. E. Characterization of a Polydnavirus (PDV) and Virus-like Filamentous Particle (VLFP) in the Braconid Wasp Cotesia Congregata (Hymenoptera: Braconidae). Journal of Invertebrate Pathology 59, 315–327. ISSN: 00222011 (May 1992).

34. B Guinet et al. Diversity and Evolution of a Novel Filamentous dsDNA Virus Family Associated with Parasitic Wasps. Virus Evolution (2024).

35. Coffman K. A., Hankinson Q. M. & Burke G. R. A Viral Mutualist Employs Posthatch Transmission for Vertical and Horizontal Spread among Parasitoid Wasps. Proceedings of the National Academy of Sciences 119, e2120048119. ISSN: 0027-8424, 1091-6490 (Apr. 2022).

36. Robertson P. L. A Morphological and Functional Study of the Venom Apparatus in Representatives of Some Major Groups of Hymenoptera. Australian Journal of Zoology 16, 133–166. ISSN: 1446-5698 (1968).

37. David J. A New Medium for Rearing Drosophila in Axenic Conditions. Drosophila Info Service, 36:128 (1962).

38. Yang J. et al. Prevention of Apoptosis by Bcl-2: Release of Cytochrome c from Mitochondria Blocked. *Science (New York*, N.Y.) 275, 1129–1132. ISSN: 0036-8075 (Feb. 1997).

39. Beld M., Sol C., Goudsmit J. & Boom R. Fractionation of Nucleic Acids into Single-Stranded and Double-Stranded Forms. Nucleic Acids Research 24, 2618–2619. ISSN: 0305-1048 (July 1996).

40. Boom R. et al. Rapid and Simple Method for Purification of Nucleic Acids. Journal of Clinical Microbiology 28, 495–503. ISSN: 0095-1137 (Mar. 1990).

41. Meyer M., Stenzel U. & Hofreiter M. Parallel Tagged Sequencing on the 454 Platform. Nature Protocols 3, 267–278. ISSN: 1750-2799 (2008).

42. Lepetit D., Gillet B., Hughes S., Kraaijeveld K. & Varaldi J. Genome Sequencing of the Behaviour Manipulating Virus LbFV Reveals a Possible New Virus Family. Genome Biology and Evolution, evw277. ISSN: 1759-6653 (Jan. 2017).

43. Suvorov A. et al. Widespread Introgression across a Phylogeny of 155 Drosophila Genomes. Current Biology 32, 111–123.e5. ISSN: 09609822 (Jan. 2022).

44. Kumar S. et al. TimeTree 5: An Expanded Resource for Species Divergence Times. Molecular Biology and Evolution 39, msac174. ISSN: 1537-1719 (Aug. 2022).

45. Peters R. S. et al. Evolutionary History of the Hymenoptera. Current Biology 27, 1013–1018. ISSN: 09609822 (Apr. 2017).

46. Blaimer B. B., Gotzek D., Brady S. G. & Buffington M. L. Comprehensive Phylogenomic Analyses Re-Write the Evolution of Parasitism within Cynipoid Wasps. BMC Evolutionary Biology 20, 155. ISSN: 1471-2148 (Dec. 2020).

47. Peres-Neto P. R., Legendre P., Dray S. & Borcard D. Variation Partitioning of Species Data Matrices: Estimation and Comparison of Fractions. Ecology 87, 2614–2625. ISSN: 0012-9658 (Oct. 2006).

48. Wallace M. A. et al. The Discovery, Distribution, and Diversity of DNA Viruses Associated with *Drosophila Melanogaster* in Europe. Virus Evolution 7, veab031. ISSN: 2057-1577 (Jan. 2021).

49. Carton Y., Poirié M. & Nappi A. J. Insect Immune Resistance to Parasitoids. Insect Science 15, 67–87. ISSN: 16729609 (Jan. 2008).

50. Nigg J. C. & Falk B. W. Diaphorina Citri Densovirus Is a Persistently Infecting Virus with a Hybrid Genome Organization and Unique Transcription Strategy. Journal of General Virology 101, 226–239. ISSN: 0022-1317, 1465-2099 (Feb. 2020).

51. Schoonvaere K., Smagghe G., Francis F. & de Graaf D. C. Study of the Metatranscriptome of Eight Social and Solitary Wild Bee Species Reveals Novel Viruses and Bee Parasites. Frontiers in Microbiology 9, 177. ISSN: 1664-302X (Feb. 2018).

52. Yinda C. K. et al. Cameroonian Fruit Bats Harbor Divergent Viruses, Including Rotavirus H, Bastroviruses, and Picobirnaviruses Using an Alternative Genetic Code. Virus Evolution 4. ISSN: 2057-1577 (Jan. 2018).

53. Cross S. T. et al. Partitiviruses Infecting Drosophila Melanogaster and Aedes Aegypti Exhibit Efficient Biparental Vertical Transmission. Journal of Virology 94 (ed Pfeiffer J. K.) e01070–20. ISSN: 0022-538X, 1098-5514 (Sept. 2020).

54. Webster C. L. et al. The Discovery, Distribution, and Evolution of Viruses Associated with Drosophila Melanogaster. PLOS Biology 13 (ed Malik H. S.) e1002210. ISSN: 1545-7885 (July 2015).

55. Webster C. L., Longdon B., Lewis S. H. & Obbard D. J. Twenty-Five New Viruses Associated with the Drosophilidae (Diptera). Evolutionary Bioinformatics 12s2, EBO.S39454. ISSN: 1176-9343, 1176-9343 (Jan. 2016).

56. van Mierlo J. T. et al. Novel Drosophila Viruses Encode Host-Specific Suppressors of RNAi. PLoS Pathogens 10 (ed Schneider D. S.) e1004256. ISSN: 1553-7374 (July 2014).

57. Martinez J., Lepetit D., Ravallec M., Fleury F. & Varaldi J. Additional Heritable Virus in the Parasitic Wasp Leptopilina Boulardi: Prevalence, Transmission and Phenotypic Effects. Journal of General Virology 97, 523–535. ISSN: 0022-1317, 1465-2099 (Feb. 2016).

58. Hermanns K., Zirkel F., Kurth A., Drosten C. & Junglen S. Cimodo Virus Belongs to a Novel Lineage of Reoviruses Isolated from African Mosquitoes. Journal of General Virology 95, 905–909. ISSN: 0022-1317, 1465-2099 (Apr. 2014).

59. Dhaygude K., Johansson H., Kulmuni J. & Sundström L. Genome Organization and Molecular Characterization of the Three *Formica Exsecta* Viruses—FeV1, FeV2 and FeV4. PeerJ 6, e6216. ISSN: 2167-8359 (Feb. 2019).

60. Ballinger M. J., Bruenn J. A., Hay J., Czechowski D. & Taylor D. J. Discovery and Evolution of Bunyavirids in Arctic Phantom Midges and Ancient Bunyavirid-Like Sequences in Insect Genomes. Journal of Virology 88 (ed Doms R. W.) 8783–8794. ISSN: 0022-538X, 1098-5514 (Aug. 2014).

61. Obbard D. J., Shi M., Roberts K. E., Longdon B. & Dennis A. B. A New Lineage of Segmented RNA Viruses Infecting Animals. Virus Evolution 6, vez061. ISSN: 2057-1577 (Jan. 2020).

62. Spear A., Sisterson M. S., Yokomi R. & Stenger D. C. Plant-Feeding Insects Harbor Double-Stranded RNA Viruses Encoding a Novel Proline-Alanine Rich Protein and a Polymerase Distantly Related to That of Fungal Viruses. Virology 404, 304–311. ISSN: 00426822 (Sept. 2010).

63. Schoonvaere K. et al. Unbiased RNA Shotgun Metagenomics in Social and Solitary Wild Bees Detects Associations with Eukaryote Parasites and New Viruses. PLOS ONE 11 (ed Lu R.) e0168456. ISSN: 1932-6203 (Dec. 2016).

64. Kondo H., Chiba S., Maruyama K., Andika I. B. & Suzuki N. A Novel Insect-Infecting Virga/Nege-like Virus Group and Its Pervasive Endogenization into Insect Genomes. Virus Research 262, 37–47. ISSN: 01681702 (Mar. 2019).

65. Mondotte J. A. et al. Evidence For Long-Lasting Transgenerational Antiviral Immunity in Insects. Cell Reports 33, 108506. ISSN: 22111247 (Dec. 2020).

66. Koonin E. V., Dolja V. V. & Krupovic M. Origins and Evolution of Viruses of Eukaryotes: The Ultimate Modularity. Virology 479–480, 2–25. ISSN: 00426822 (May 2015).

67. Raghwani J. et al. Seasonal Dynamics of the Wild Rodent Faecal Virome. Molecular Ecology, mec.16778. ISSN: 0962-1083, 1365-294X (Nov. 2022).

68. Lopez-Ferber M. & Comendador M. A. Transmission of the Hereditary S Character in Drosophila Simulans: Effects of Aging and Temperature Treatments. Journal of Invertebrate Pathology 59, 264–270. ISSN: 00222011 (May 1992).

69. Olazcuaga L. et al. A Whole-Genome Scan for Association with Invasion Success in the Fruit Fly Drosophila Suzukii Using Contrasts of Allele Frequencies Corrected for Population Structure. Molecular Biology and Evolution 37 (ed Singh N.) 2369–2385. ISSN: 0737-4038, 1537-1719 (Aug. 2020).

70. Prenter J., MacNeil C., Dick J. T. & Dunn A. M. Roles of Parasites in Animal Invasions. Trends in Ecology & Evolution 19, 385–390. ISSN: 01695347 (July 2004).

71. Martinez J., Fleury F. & Varaldi J. Heritable Variation in an Extended Phenotype: The Case of a Parasitoid Manipulated by a Virus: Heritable Variation in Extended Phenotype. Journal of Evolutionary Biology 25, 54–65. ISSN: 1010061X (Jan. 2012).

72. Carton Y., Bouletreau M., van Alphen J. J. M. & van Lenteren J. C. In The Genetics and Biology of Drosophila, Vol. 3 348–394 (Academic Press, 1986).

73. Varaldi J., Patot S., Nardin M. & Gandon S. In Advances in Parasitology 333–363 (Elsevier, 2009). ISBN: 978-0-12-374792-1.

74. Chabert S., Allemand R., Poyet M., Eslin P. & Gibert P. Ability of European Parasitoids (Hymenoptera) to Control a New Invasive Asiatic Pest, Drosophila Suzukii. Biological Control 63, 40–47. ISSN: 10499644 (Oct. 2012).

75. Kacsoh B. Z. & Schlenke T. A. High Hemocyte Load Is Associated with Increased Resistance against Parasitoids in Drosophila Suzukii, a Relative of D. Melanogaster. PLoS ONE 7 (ed DeSalle R.) e34721. ISSN: 1932-6203 (Apr. 2012).

